# PlanktonScope: Affordable modular imaging platform for citizen oceanography

**DOI:** 10.1101/2020.04.23.056978

**Authors:** Thibaut Pollina, Adam G. Larson, Fabien Lombard, Hongquan Li, Sebastien Colin, Colomban de Vargas, Manu Prakash

## Abstract

The planktonic communities within our oceans represent one of the most diverse and understudied ecosystems on the planet. A major hurdle in describing these systems is the sheer scale of the oceans along with logistical and economic constraints associated with their sampling. This is due to the limited amount of scientifically equipped fleets and affordable equipment. Here we demonstrate a modular approach for building a versatile, re-configurable imaging platform that can be adapted to a number of field applications, specifically focusing on oceanography. By using a modular hardware/software approach for building microscopes, we demonstrate high-throughput imaging of lab and field samples while enabling rapid device reconfiguration in order to match diverse applications and the evolving needs of the sampler. The presented versions of PlanktonScope are capable of autonomously imaging 1.7 ml per minute with a 1.5 µm resolution, and are built with under $400 in parts. This low cost enables new applications in laboratory settings such as the continuous imaging of suspension cultures, and in-field settings with the ability to scale up for long-term deployment on an international fleet of sailing boats enabling citizens based oceanographic research

## Introduction

Planktonic communities are the foundations of the global ecological network in the ocean (Fenchel 1988). Together, these communities fix from 30 to 50% of the world’s carbon emissions (le B. Williams, Thomas, and Reynolds 2008). Despite being an essential component of the biosphere, we know astonishingly little about the level of diversity within these communities, or the scope of impact that human activities are having on them. This ecosystem is under the direct influence of microplastics (Long et al. 2015), urban, agricultural and industrial waste (Lebreton et al. 2017), (Rocha et al. 2015), acoustic perturbations (Howe et al. 2019), and nuclear waste (Kautsky et al. 2016), as well as indirect influences such as climate change (Hays, Richardson, and Robinson 2005). Marine ecologists also face a continual challenge of cataloging a growing list of newly discovered species as many remain unknown, or are very poorly characterized and therefore often unidentifiable. Monitoring the health of such a vast number of species under such varied conditions, and over spatial scales that encompass entire oceans, remains difficult.

Quantitative imaging methods, such as flow imaging microscopy, are an efficient way to monitor both the diversity and quantity of micro and meso-scale plankton communities (Lombard et al. 2019)) in conjunction with the many environmental or anthropogenic stressors (Kautsky et al. 2016) that influence themSeveral high-throughput and highly automated imaging instruments such as the IFCB (Olson and Sosik 2007) and the FlowCam (Buskey and Hyatt 2006) have been developed for this task. However, these instruments are expensive, bulky, and not suitable for small boats, which prevent their adoption on a larger scale.

Additionally, the current strategy for sampling the ocean relies heavily on large and expensive oceanographic expeditions, together with repeated geochemical and biological measurements at fixed but sparse stations. Consequently, most of the ever-evolving oceanic habitats remain largely undersampled. However, scientific fleets are not the only entities exploring the ocean. Thousands of citizen sailors traverse the oceans each year in sailboats exploring and living within these vast ecosystems.

The eagerness of these potential citizen scientists along with the value of their thousands of transects creates a niche for scalable tools that can enable a global survey of the oceans by generating reliable data. Frugal tools become an effective way to tackle the problem of cost-, and spatiotemporal-limiting oceanography. For example, Foldscope (Cybulski, Clements, and Prakash 2014) is a cost-effective, portable microscope with over a million copies around the world, and has successfully enabled a global community of microscopists to share their data and discoveries. Similarly, there exists a need for a low-cost portable microscope to be used directly at sea that can involve the vast community of sailors continually traversing bodies of water across the planet.

Since modularity is a natural way to make complexity manageable and to accommodate uncertainty in the evolution of design (Efatmaneshnik and Ryan 2016), we used this approach to construct a complex instrument composed of simple modules. Every module encapsulates a simple function allowing scientists and makers to build off of this platform. This strategy allows the device to be easily upgraded instead of replaced as a whole, providing an affordable way to take on unforseen future applications.

The modular approach for building a versatile, re-configurable imaging platform can be adapted to a number of field and laboratory applications, particularly, but not at all limited to, oceanography. We demonstrate the efficiency of our system in obtaining high-throughput imaging of lab and field samples while enabling rapid reconfiguration in order to match the evolving needs of oceanographers.

For costs under $400, the presented designs allows for adaptation to a number of imaging applications. By connecting the modules together, mechanically and electronically, as well as the optical and fluidic paths, these designs can host a whole breadth of new functional layers embedded in physical modules.

The basis of the largest ecosystem on earth lies within the planktonic organisms invisible to the naked eye. To understand this world we need to magnify not only this hidden world, but also our approach. Deploying a high-throughput microscope platform on a global scale will bring light to the habitats under-surveyed by the large and more infrequent research cruises. Connecting these platforms with a network of climate-researchers, ecologists, citizen scientists, and many others across the planet will bring further relevance to each individual measurement.

## Results

### Open imaging platform

Here we present a flow-based imaging platform in two configurations: V.1) a totally modular, compartmentalized configuration enabling multi-functionality and complete adaptability, and V.2) a compact version designed for robust performance and portability. Both achieve optical magnification of 0.75X and a pixel size of 1.5µm/px. The travel distance of the focus stage is about 3.2cm for a step size of 0.15µm. For a flow-rate estimated at 3ml/min and a flow-cell of 500µm thick, we can image a volume of 1.7ml/min. The platform is built around an open-hardware principle. The components are off-the-shelf and readily accessible from numerous vendors at low cost. The corresponding open-software strategy utilizes existing libraries for the interfacing and image processing in addition to a flow-based visual programming platform to enable any user to rapidly customize acquisition and processing steps.

#### V.1: The compartmentalized version (Fig. 1)

The fully modular version is based on six triangular units, each of which is a separate functional layer that couple together through shared optical and electronic paths : 1) a single board computer coupled to its camera sensor, 2-3) two reversed M12 lenses separated in two modules (an objective lens and a tube lens), 4) a motorized stage and focus delta platform for sample manipulations, 5) a module embedding independent programmable rings of LEDs for the illumination module, and 6) a laser cut peristaltic pump (Fig. 1A). The modularity is rooted in an interface designed to connect modules mechanically, electronically, and optically, all in numerous easily alterable configurations (Fig. 1B). The prototype can be used either in the lab (Fig. 1C) or in the field (Fig. 1D).

**Figure 1.**
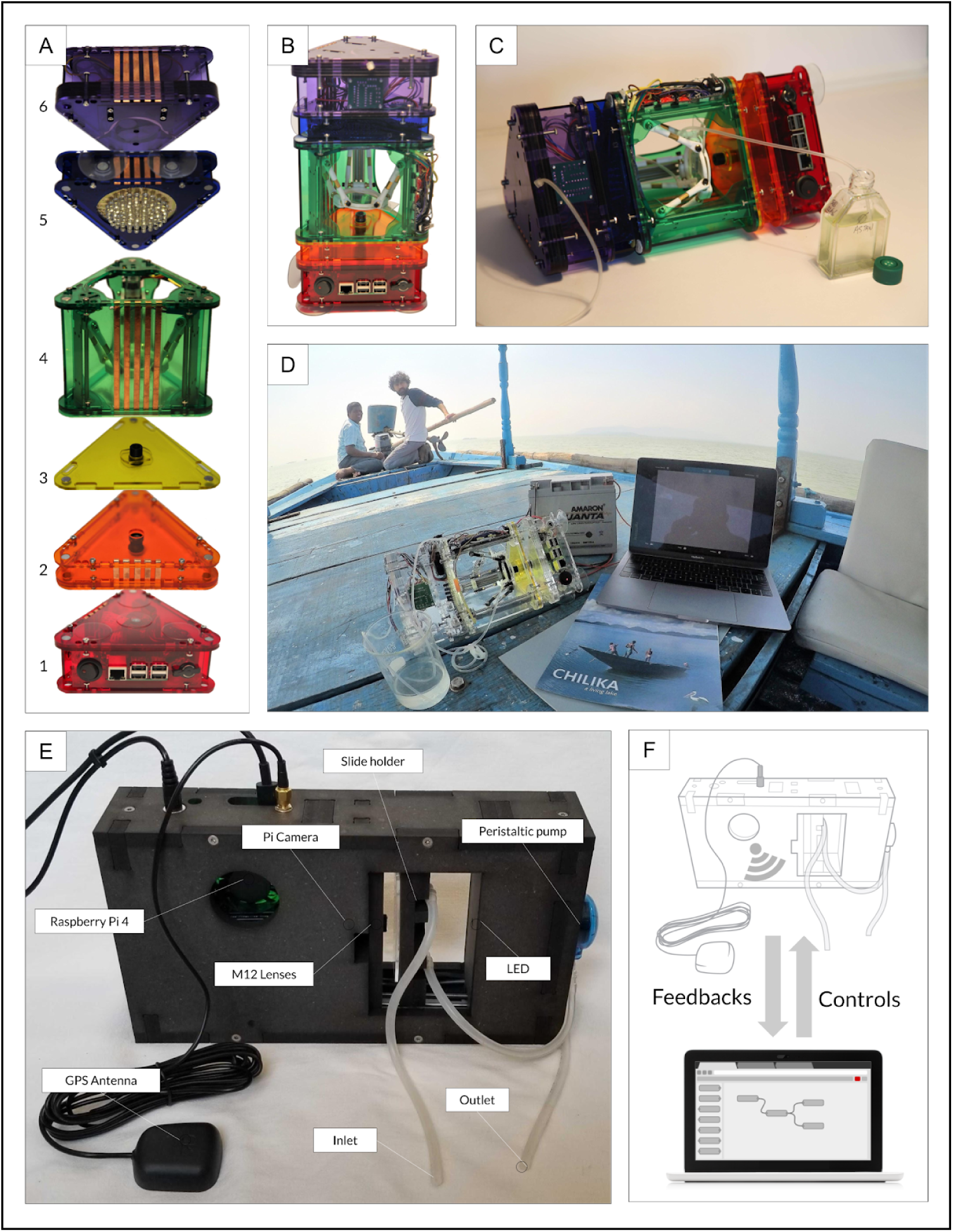
Modular and monolithic Planktonscope design compared. (A) Modular stackable flow-through microscope design with blocks that stack together (bottom to top) : Computational/imaging sensor (1), tube lens (2), objective lens (3), delta stage (4) for sample manipulation and focus including flow cell mount, illumination (5) and the pump (6). The platform can be re-assembled and is held together by a stack of magnets. (B) Planktonscope can be used in vertical configuration for static imaging or (C) horizontal configuration for flow through samples. (D) Deployment of the fully modular version of the PlanktonScope on board a traditional fishing boat in lake Chilika (Orissa, India), operating autonomously on a 12V car battery. (E) Monolithic portable Planktonscope with fixed configuration. (F) Planktonscope is controlled via smartphone or laptop allowing real-time feedback during data collection and processing.

**Figure 2.**
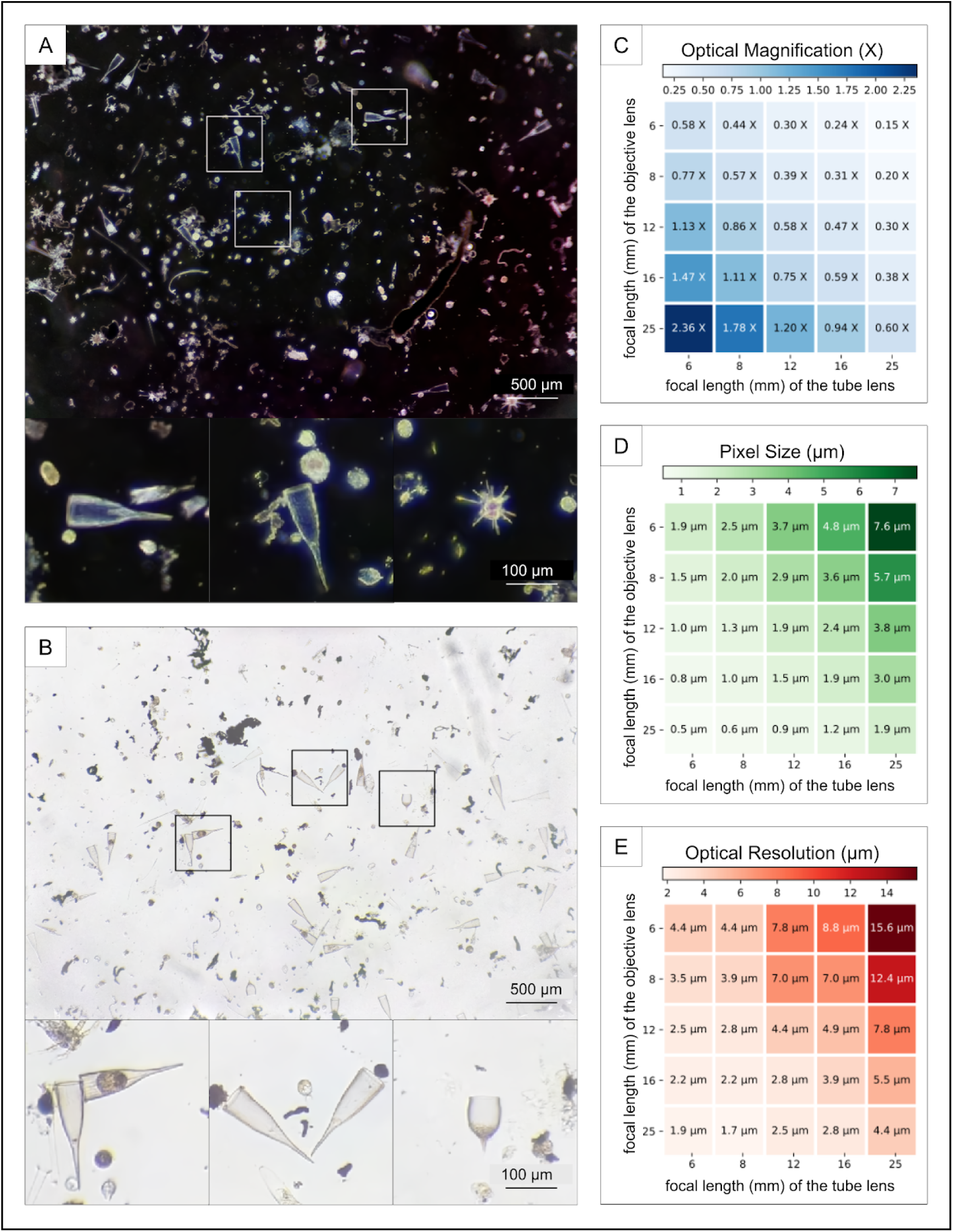
Optical characterization for various modular configurations. (A) Image of typical plankton tow under dark-field illumination. Insets depict zoom in on specific species. (B) Image of typical plankton tow under bright-field illumination. Insets depict zoom in on specific species. Optical magnification (C), pixel resolution and (D) optical resolution of the different optical configuration from different M12 lens configurations.

#### V.2: The flow-through version (Fig. 1E)

The second prototype (Fig. 1E) is an example of how the platform can be specialized for flow-through acquisition, complete with a robust electrical connectivity allowing for stable deployment in field conditions. A single board computer controls a focus actuator holding the flow-cell, a single LED for bright-field illumination, a peristaltic pump, and a GPS.

### Open HardWare

#### Mechanical understanding of the prototypes (Fig. S1)

The designs presented here are built around a laser cut framework. Both designs are parametric, enabling the use of different thicknesses of the chosen material. These range from acrylic and recycled plastic, to wood, metal, or fiberboard. This machining method allows rapid design iteration and enables a precise, low-cost method for aligning and spacing optical components.

#### V.1: Modular version

In the modular version, to physically couple the units, three neodymium magnets are incorporated into the corners of the interface between modules (Fig. 1A) enabling both proper alignment and quick reconfiguration. Following this interface, the prototype can be intermachine compatible as each module can be readily 3D printed.

The microscope can be used in vertical or horizontal configurations, placed upright or inverted, depending on the need or constraints of the experimenter. For example, a vertical manual mode enables exploration of a static sample that can be placed on a glass slide, flow-cell, or petri dish (Fig. 1B). The delta stage gives the user the ability to track an organism with high precision, or quickly survey the area of the sample holder. A horizontal autonomous mode gives the user the ability to continuously image liquid samples passing through a flow-cell at a desired rate. This enables continual imaging of a large volume (Fig. 1C).

#### V.2: Flow-through version

The flow-through version simplifies the assembly of the modular prototype by using a minimal structure to position and align the components. Designed to increase the robustness and stability of the prototype for field deployment or in-situ installation, a level of modularity is maintained by allowing the lenses and flow-cell to be quickly swapped. While the modular version requires a considerable time of 10 hours for the machining, soldering, and assembly, the flow-through version drastically reduces the build complexity enabling a complete machining/assembly in less than 4 hours.

#### Electronic modularity and autonomy

Both prototypes are based on a Raspberry Pi single board computer which enables a centralized method to control the electronics, acquire and process the images, and serve as the user/machine interface(Fig. S1).

The magnetic coupling of the modular unit enables electronic connectivity through the contact of copper ribbons that connect each module at their interface to form a custom BUS for I^2^C connection and power. The different independent microcontrollers, here Arduinos, receive queries as actuators from the Raspberry Pi and send logs as sensors back to the Raspberry Pi.

Instead of utilizing a custom BUS which is time consuming to establish, the flow-through version utilizes Pi HATs (Hardware Attached on Top) for both assembling and deploying code in the Arduinos. The Pi HATs enable the rapid addition of numerous off-the-shelf specialized boards. Three HATs are utilized here to serve different functions: one for cooling the CPU of the Raspberry Pi, one for controlling the focus stage and the pump, and a final HAT supporting a GPS for geolocalization. Thanks to the massive community built around Raspberry Pi, hundreds of other possibilities exist for new modules and more functions built on top of this platform.

Both instruments can be powered either through standard wall AC power or from battery cells for field deployment (Fig. 5A). For an acquisition frequency of 0.5 Hz and a standard Li or polymer battery of 20,000mAh, the flow-through version can collect continuously for more than 5 hours.

#### Illumination choice

The illumination in the modular version is built of 5 different concentric rings composed of 1, 6, 12, 24 and 32 white ultra bright LEDs having a narrow angle of 17°. The light intensity of each ring can be tuned separately to offer a broad range of illumination modes. Two main modes are pure dark-field where the two external rings are alight (Fig. 2A) and pure bright-field where the most central LED is used (Fig. 2B). In the following results, we opted to use the maximum light intensity of the central LED to maximize the depth of field in the flow-cell.

The flow-through version uses a single LED which provides an almost collimated light source due to the narrow angle of illumination. This achieves a large depth of field for imaging samples with a large size variance.

#### Low cost and expandable optical configuration

The recent improvements of cheap, high quality sensors have led to the concurrent development of high performance and frugal lenses in a compact form factor. They are termed ‘M12’ corresponding to the metric tapping dimension. By orienting two M12 lenses toward each other we construct an affordable and reconfigurable solution to project the image of a microscopic object to a camera sensor(Fig. 2).

By using different focal lengths for both lenses, measuring the size of the field of view, and calculating the resolution of each combination, we obtained a comparative matrix of 25 different optical configurations. We observe (Fig. 2C) a gradient from low magnification (0.15X) to high magnification (2.36X) respectively offering a pixel size (Fig. 2D) of 0.5µm or 7.6µm. Since each version allows magnetic swapping of both lenses, all the described optical configurations are readily interchangeable.

#### Stage and focus

The modular version embeds a layer which combines two important functions in microscopy into a compact form factor: focusing and exploring a sample. This linear delta design, well known for its important development and use in 3D printers, uses 3 vertical linear stepper motors that hold a platform, each with 2 arms. The platform made of two separable magnetic bodies can host a broad range of sample holders: a slide, a petri dish, or a flow-cell. Focusing is made possible by controlling 3 independent drivers wired to simultaneously move the stepper motors up or down. The travel distance of the platform measures about 3.2cm with a step size of 0.15µm. This allows fine control of movement to accurately track and image micron sized objects. For the price of about $30, this represents an affordable way to construct a motorized XYZ stage.

Since the flow-through version is specialized in fluidic acquisition, there is no need for a stage. The flow-cell can be actuated for fine focus using 2 synchronous linear stepper motors offering a step size of 0.15µm on a travel distance measuring about 2.5cm.

#### Two flow-through strategies

The fluidic strategy can be distinguished in two different modes, a continuous flow mode and a stop flow mode. In continuous mode the peristaltic pump is continuously rotating at low flow-rate while the camera is taking images at a given frame-rate. The stop-flow method synchronizes the rotation of the pump to the image capture in order to stop the pump when each image is taken. Since Pi Camera is based on a rolling shutter, the imaged objects undergo a morphological deformation when imaged under continuous flow. In addition, peristaltic pumps have a pulsed flow which is difficult to characterize, making post-acquisition correction difficult.

#### Stop flow enables high quality imaging

Since stop-flow synchronizes the acquisition and the pump, the objects are stationary when imaged, canceling any morphological deformation and allowing quantitative analysis. Moreover, this enables a longer exposure, increasing the resolution and reducing the need for powerful illumination. This lower frame-rate strategy enables the capture of the full camera sensor for a larger field of view than via the continuous mode.

### Open Software (Fig. S4 software architecture)

The entire software architecture (Fig. S4) is based on existing programs and python libraries such as Node-RED for the Graphical User Interface and first layer of programming interface, MorphoCut for handling the image processing from the raw images to the online platform, and EcoTaxa for classification and annotation.

#### User/Machine interface

By utilizing the headless configuration for the Raspberry Pi, we removed the need for a dedicated monitor, mouse and keyboard, enabling control of the instrument from any device able to access a browser over a WiFi connection (Fig. 1F). This strategy enables any user to immediately interact with the device without OS or software compatibility issues. The user can then access a browser based dashboard powered by Node-RED for an interactive collection of the metadata crucial for acquisition as well as rapid state modification of the actuators.

#### Flow based programming strategy

The back-end of the GUI is also based on Node-RED, a flow-based development tool for visual programming which is provided by default on any Raspberry Pi software suite. Node-RED provides a web browser-based flow editor, which can be used to create JavaScript functions. Elements of applications can be saved or shared for re-use. The strategy makes it more accessible to those with limited experience in scripting. This visually modifiable program can easily be shared over a json text file.

#### Image processing

The current workflow has image processing done automatically after the acquisition of a defined batch of images. They are then processed by MorphoCut, a python-based library designed to handle large volumes of imaging data.

Several operations (Fig. 3A) are applied on the raw images acquired in fluidic mode: first an average background from five frames surrounding the current one is calculated for subtraction (1), current raw frame (2), current frame with background removed (3), threshold (4), labeling (5). This way, large data sets can be compressed on-board by storing only relevant regions of interest.

**Figure 3.**
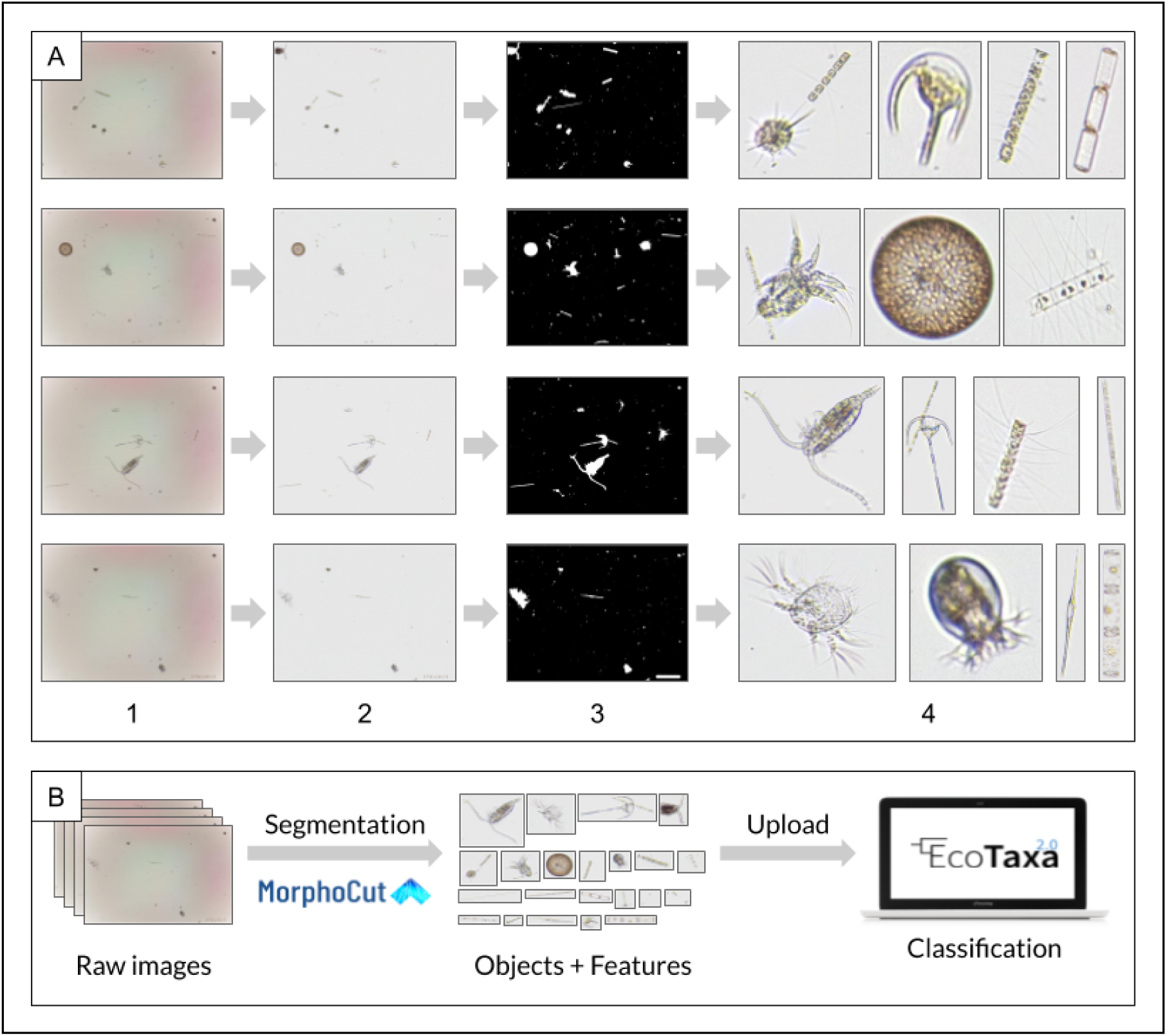
Image processing pipeline for fluidic analysis (A) Workflow used to segment the objects imaged in a frame and to extract features including an average background from 5 frames surrounding the current one (1), current raw frame (2), current frame with background removed (3), threshold (4) extract objects and label. Large data sets collected can be compressed on-board by storing only relevant regions of interest. (B) Overview of the image processing pipeline utilizing *Morphocut* (https://github.com/morphocut/morphocut) for data cleanup uploaded in *Ecotaxa*.

MorphoCut extracts different features on each region of interest that are stored, along with contextual metadata generated by the user on the GUI, in a compressed file ready to be uploaded on EcoTaxa (Fig. 3B). This creates a uniform data format already utilized by marine researchers worldwide.

### Benchmarking

#### Imaged volume and frame-rate

This pump in the modular system is a peristaltic pump composed of a stack of 5 acrylic layers forming a closed chamber inside which 3 “rollers” can spin around the motor axis compressing a tube along the internal wall. The speed of the motor as well as the diameter of the compressed tube determines the flow-rate which is about 3 ml/min at maximum speed. Since the field of view is smaller than the section of the flow-cell, the volume seen on the imaging sensor is smaller than the volume present in the section of the flow-cell. The optical configuration described below paired with a flow-cell of 5mm long and 500µm deep allows one to image a volume of 2.33µl per frame for a volume of 4.05µl present in the complete volume of the flow-cell. In order to avoid repeatedly imaging the same object, we calculated a maximum frame-rate of 12 frames/second. With these settings, about 1.68ml sample is imaged per minute and the device will take less than 9 minutes to image 15ml of liquid sample from 27ml passing through the flow-cell.

The compact version uses off-the-shelf peristaltic pumps for flow. Several models can be easily incorporated by small modifications to the laser cut mount on the CAD file.

#### Monitoring lab culture morphology (Fig. 4)

The Planktonscope is designed with rapid adaptability in mind. In this manner, it is designed to be transported quickly from designated use in the field to controlled data collection in a laboratory setting. To benchmark its ability to function as a lab culture monitoring system we used a continual flow mode to image monocultures of Pyrocystis noctiluca and Coscinodiscus wailesii.

**Figure 4.**
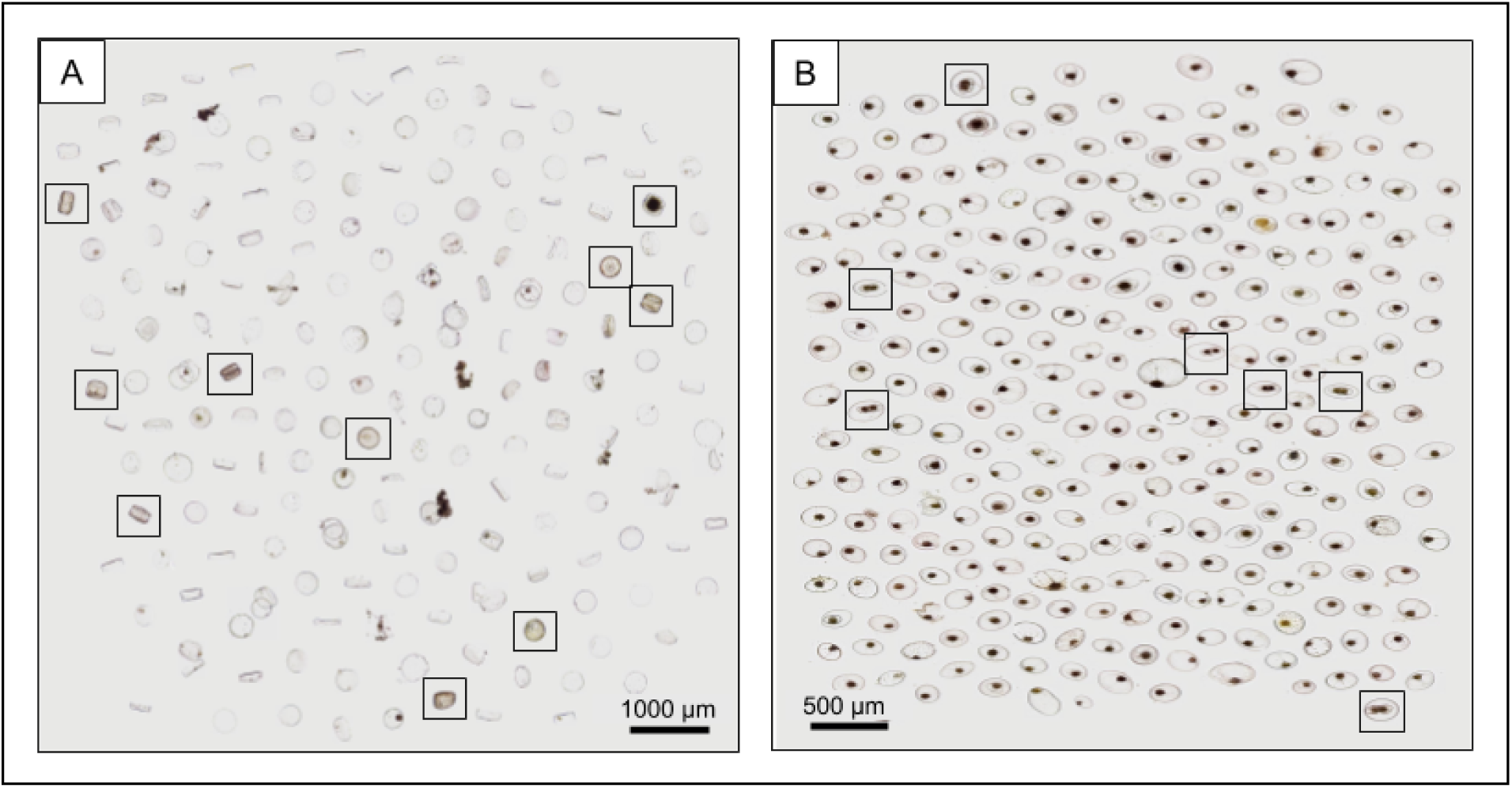
Non-destructive continuous monitoring of lab cultures using a Planktonscope allows for cell state to be measured at single cell resolution. **A)** Coscinodiscus wailesii cultures were monitored over a period of 6 hours. Creating a simple montage allows the user to easily quantify living or dead cells at different time points. **B)** Pyrocystis noctiluca cultures were monitored over a period of 6 hours during their conditioned night to day transition. Dividing cells are easily identifiable.

**Figure 5.**
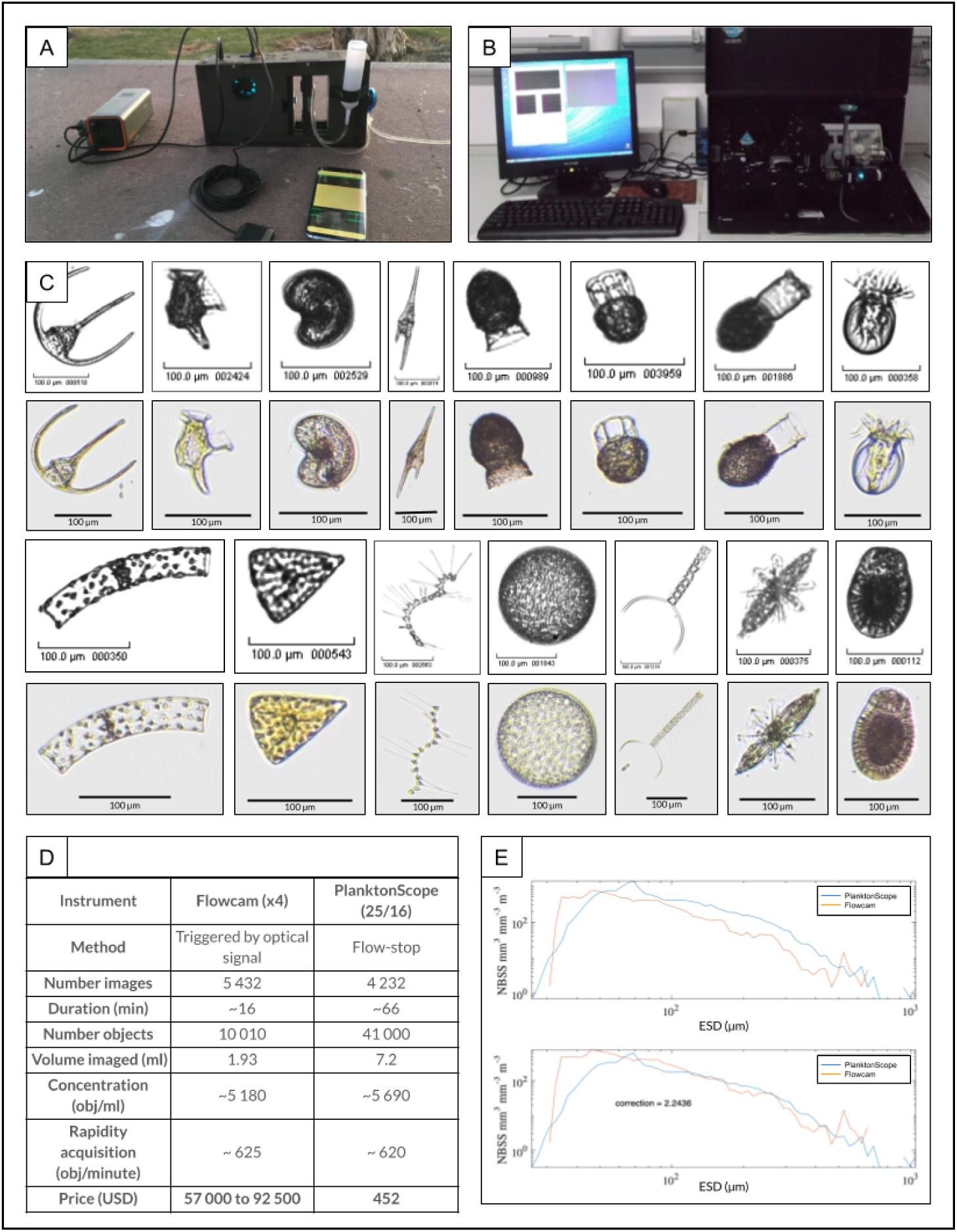
One to one comparison of plankton scope with another traditional flow through imaging systems (Flowcam) Ultra portable configuration of the Planktonscope operated on a cell phone charger and controlled through a user interface on a smartphone. (B) Typical setup of a FlowCam on a laboratory bench. (C) One to one comparison of the same sample (plankton tow, Villfranche) passed through a Planktonscope and flowcam. A few representative images are chosen from the two data sets - first row from left to right : *Ceratium spp., Dinophysis caudata, Peridiniales spp., Ceratium furca, Codonaria spp., Dictyocysta spp., Codonellopsis spp., Undellidae spp.* Second row from left to right : *Guinardia spp., Licmophora spp., Asterionellopsis spp., Coscinodiscophyceae spp., Chaetoceros spp., Acantharea, unknow sp.* (D) Table comparing efficiencies for both trigger based optical image collection and flow-stop base wide field of view imaging and computational segmentation. We find that when normalized for the total number of objects detected - Planktonscope performed equally well compared to flow cam (625 objects detected per minute detected with Flowcam compared to 620 objects detected per minute with Planktonscope). (E) Comparison of total planktonic organisms (objects) sampled with different collection methods and analyzed with different optical/imaging methods as a function of the size of organisms (expressed as equivalent spherical diameter; ESD). Total organism biovolume per size classes were expressed as normalized biovolume size spectra (NBSS) by dividing the total biovolume within a size class by the biovolume interval of the considered size class. NBSS is representative of the number of organisms within a size class. Results were obtained by imaging a plankton net sample with Flowcam with a 4X lens and PlanktonScope. All data are raw counts and converted to biovolume using ellipsoidal calculations. The low count at the smaller size range of each observation corresponds to an underestimation of an object’s number due to both limited capabilities of each imaging device for small objects and net under sampling for small objects utilizing the plankton tow.

We continually flowed a culture of diatoms through the system to monitor culture viability. After segmentation it was straightforward to distinguish and quantify a ratio of dead to living cells (Fig. 4A). By imaging the large transparent dinoflagellate Pyrocystis noctiluca, we can observe (Fig. 4B) the cell morphology evolving over time. In this manner we can easily build a diagram of temporal phenotypes. As circadian clocks function as major drivers of behavior in many marine protists, these types of controlled continual monitoring could serve as a source of informative, non-invasive, and easily accessible information on any cultured strain.

#### FlowCam comparison (Fig. 5)

As another benchmarking measure we passed identical samples through the Planktonscope and a FlowCam to create a quantitative comparison of the instruments.

As we can observe in Fig. 5C, both instruments configured with similar magnification provide enough resolution to allow a classification down to a genus level and often to a species level.

### Field exploration

#### Planktonic biodiversity survey (Fig. 6)

This Planktonscope platform has been designed to be deployed in the field and used by experienced microscopists and citizen scientists alike. It was therefore crucial to test the apparatus in various field conditions. We tested its robustness, verified the simplicity of use, and checked its capability to acquire high quality and reproducible data. In order to qualitatively present a survey of the ecology at each field site we created collages of the most frequently extracted objects. Thus, one can draw up a representative panel of those species most prevalent in each ecosystem (Fig. 6). It is this strategy that could be scaled up to survey the massive tracts of ocean that are being explored by hundreds of sailors every day on transects across the globe.

**Figure 6.**
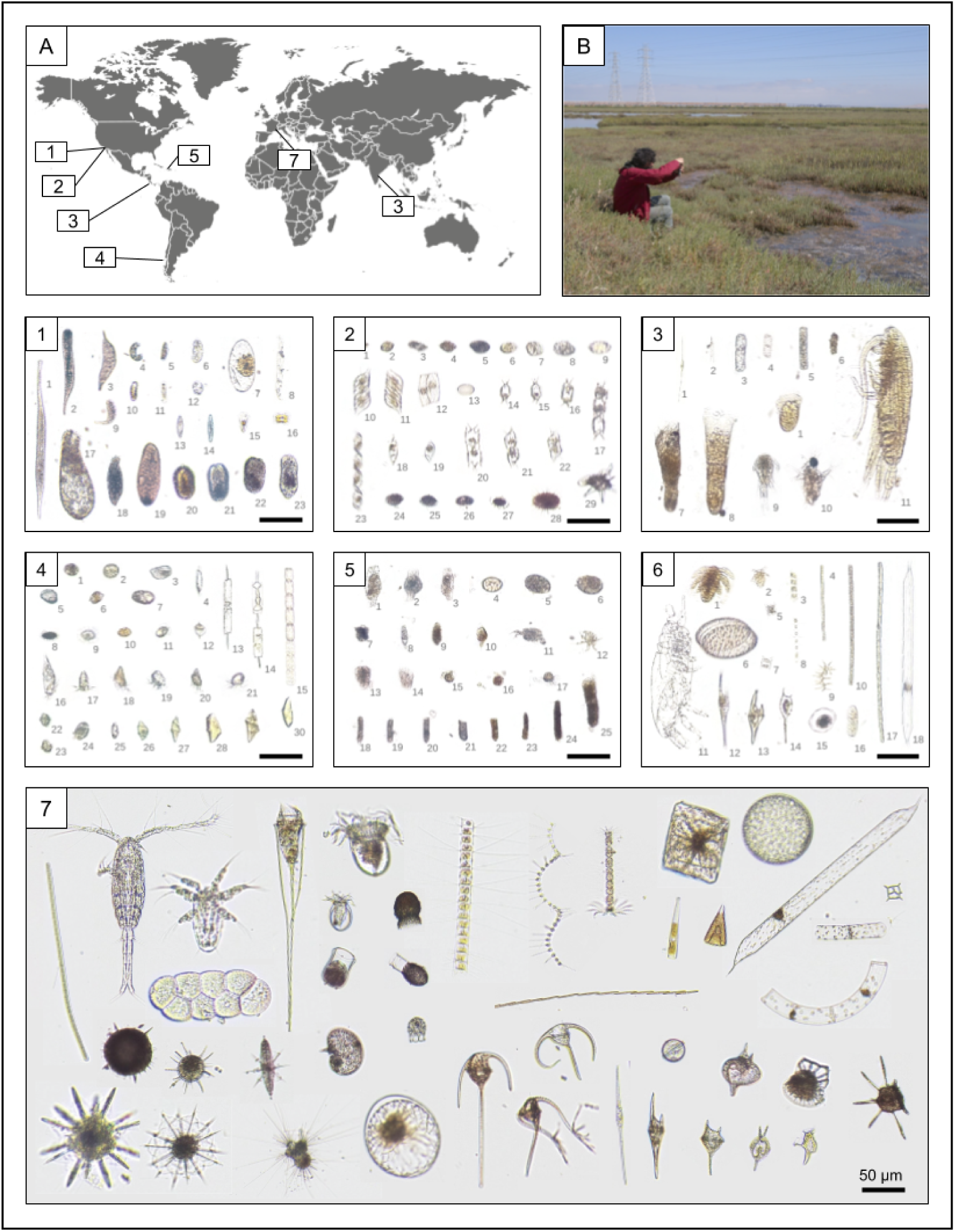
Field testing plankton scope at seven field sites around the world (A) with sampling and imaging done in the field (B) for most samples. Composite montages were made to display the objects identified with highest frequency in each ecosystem, creating a visual representation of local biomass. **1)Palo Alto Baylands Nature Preserve (USA)** - 1: Tracheloraphis, 2: Tracheloraphis, 3: Ciliate, 4: Unidentified, 5: Pennate diatom, 6: Ciliate, 7: Pyrocystis sp., 8: Gyrosigma sp., 9 10: Ciliate, 11: Pennate diatom, 12: Unidentified, 13: Pennate diatom, 14: Navicula sp., 15: Unidentified, 16: Amphiprora gigantea, 17: Enchelyodon, 18-23: Ciliate. **2) Monterey Bay (USA)** - 1-9: Unidentified, 10-12: Pennate diatom, 13: Centric diatom, 14-17: Odontella longicruris, 18 19: Unidentified diatom, 20-22: Odontella longicruris, 23: Unidentified diatom, 24-26: Armored dinoflagellate (Protoperidinium?), 27 28: Unidentified Dinoflagellate (resting cyst ?), 29: Ornithocercus. **3) Isla Secas (Panama)** - 1: Nitzschia longissima, 2 3: Unidentified, 4: Centric diatom, 5: Unidentified, 6: Copepod fecal pellet, 7: Ciliate, 8: Unidentified, 9: Copepod, 10: Crustacean larvae, 11: Calanoid copepod. **4) Comau Fjord (Chile)** - 1-6: Unidentified, 7: Unarmored dinoflagellate, 8 9: Unidentified Dinoflagellate (resting cyst), 10: Prorocentrum compressum, 11: Dinophysis sp., 12: Protoperidinium sp., 13 14: Ditylum brightwellii, 15: Detonula pumila, 16-21: Ciliate, 22-24: Lepidodinium chlorophorum, 25-30: Gyrodinium sp. **5) Isla Magueyes (Puerto-Rico)** - 1: Copepod larva, 2: Nauplius larva, 3: Chaetoceros sp., 4: Oscillatoria sp., 5: Eucampia zodiacus, 6: Coscinodiscus sp., 7: Eucampia zodiacus, 8: Chaetoceros sp., 9: Unidentified, 10: Oscillatoria sp., 11: Calanoid copepod, 12: Ceratium furca, 13: Ceratium sp., 14: Ceratium lineatum, 15: Pyrocystis sp., 16: Unidentified, 17: Oscillatoria sp., 18: Proboscia alata. **6) Chilika Lake (India)** - 1-10: Unidentified, 11: Crustacean larva, 12: Nauplius larva, 13-24: Unidentified, 25: Ciliate **7) Villfranche (France)**

The Planktonscope is ultra-portable and battery powered, able to be transported and used for the duration of a cruise or installed in-situ with the use of a dedicated power supply. It is also easily integrated into current sampling pipelines to enrich contextual parameters already being gathered.

#### Planktonic distribution study (Fig. 7)

Beyond a basic survey, we next wished to implement Planktonscope to approach a specific ecological question. The unique ecosystem present in a Patagonian fjord provided an interesting framework.

**Figure 7.**
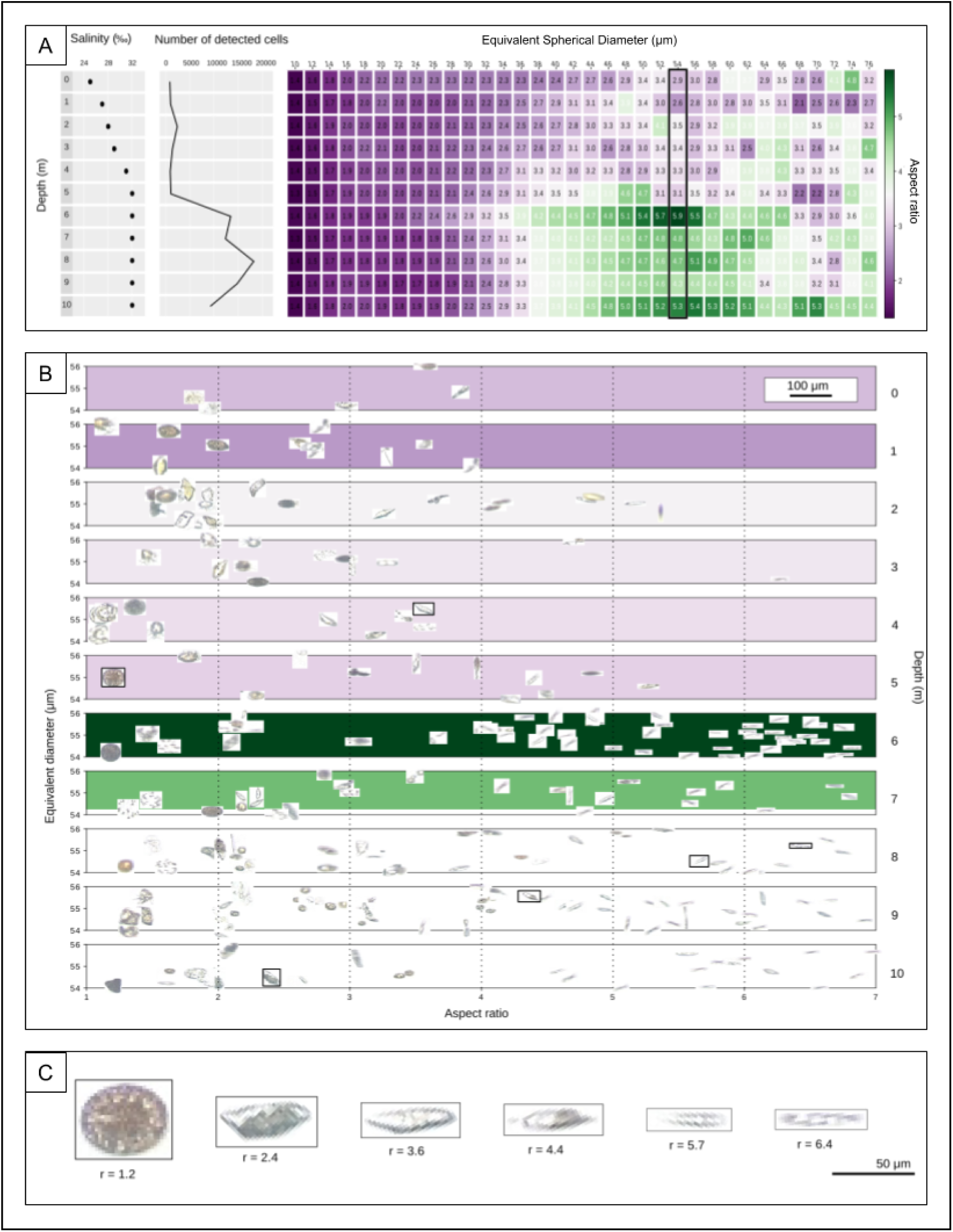
Planktonscope deployed in creating an ecosystem snapshot organized along the vertical axis in a Chilean fjord. (A). The extreme salinity gradient found in the Patagonian fjords create a unique habitat for planktonic organisms to navigate. Using depth dependent sampling and the Planktonscope we attempted to describe correlations of salinity with plankton abundance per meter of depth. Samples were collected via a siskin bottle at different depths and imaged under a planktonscope. Plot depicts vertical snapshot of an ecosystem from 0m (surface water) to 10m (depth) with number of identified objects and equivalent diameter (10 to 76 µm) as a function of the depth (0 to 10m).The measurement of the elongation per equivalent diameter is based on 94,262 objects detected in total. Color bar represents aspect ratio from purple (small aspect ratio) to green (large aspect ratio). (B). Display of distribution of objects with mean size of 54 µm as a function of the depth (0 to 10m). (C) Illustration of objects with a gradual aspect ratio from 1.2 to 6.4 marked in B.

The Comau Fjord in southern Chile receives 5 meters of rain per year, with numerous freshwater rivers and streams feeding into the saltwater bay. This provokes a vertical salinity gradient that evolves seasonally as well as with weather events such as wind and rain (Buskey and Hyatt 2006). A typical vertical stratification salinity gradient is visible where the salinity gradually increases from 25‰ at the surface to a plateau of 32% below the pycnocline at a 5 meter depth (Figure 4A). We used Planktonscope to discover if this salinity gradient results in concurrent stratification of the planktonic organisms.

The drop in salinity was accompanied by a large increase (>10fold) in concentration of detected objects in the meter of saline water immediately beneath the pycnocline. This number falls off around 9 meters below the surface, or around 4 meters below the pycnocline (Fig. 4A).

By measuring the aspect ratio of every fraction size for each meter, we observed a correlation between the increasing number of detected objects and their aspect ratio. Most of the detected objects collected below the pycnocline and with an equivalent diameter bigger than 36µm and smaller than 64µm have an aspect ratio of around 4 (Fig. 4A).

By further exploring the vertical stratification of objects with fraction size from 53 to 55µm, where the aspect ratio is on average about 5.9 at 6 meters, we find many elongated objects (Fig. 4B) with similar morphology.

As shown in figure 4C, objects representing a gradient in terms of aspect ratio from a very round cell such as a centric diatom to more elongated cells that resemble species from the genus Gyrodinium.

## Discussion

It is becoming more apparent that planktonic abundance, geographic range, and community composition are directly linked to the evolving structure and health of oceanic ecosystems (Drinkwater et al. 2003)(Hughes 2000). These biota form the basis of a global food web, and disruption of these plankton communities can be catastrophic ecologically and socioeconomically. As primary producers plankton communities also serve critical roles in the global carbon cycle. The dynamics of their distribution serve as sensitive and reliable indicators of the changing climate(Hays, Richardson, and Robinson 2005).

Further deciphering the geochemical and ecological webs that sustain the 140 million square miles of global oceans will require thousands of researchers of various disciplines to generate and organize massive amounts of data. To collect this amount of data it will require tools capable of being made, used, and modified on a massive scale.

By designing an inexpensive imaging platform that can produce high quality oceanographic data, we demonstrate the versatility of a modular approach in which the same device can serve different purposes. The Planktonscope is a low cost, rapidly modifiable, and high resolution digital microscope designed to enable large scale survey of the world’s oceans. The microscope is robust enough to perform autonomously in field conditions while maintaining the capability to robustly produce laboratory quality micrographs. By utilizing the principles of open source hardware and software, we believe we can greatly expand our knowledge of the largely unexplored oceanic ecosystems. We demonstrate here a number of applications but emphasise the foundation of Planktonscope lies in a community driven platform that is continually improved and modified to suit different needs.

### Visual data collection

There are relatively few continual time series that visually document marine ecosystems. Much of the data on our planet’s planktonic biodiversity is composed of bulk counts or bulk sequences from mixed populations without visual representations of the sampled specimens. However, much of organismal behavior and underlying cellular processes are beyond the reach of counts and genomic surveys alone. To convert this data into deeper knowledge of biodiversity we need to observe these cells and how their behavior influences the environment. Symbiosis and predation are good examples of the important interactions missed by sequencing (Colin et al. 2017). There is a growing need to supplement the current oceanographic measurements with visual representation. A frugal high throughput microscope could play roles in many different types of studies.

### Cost leads to scale

An unfortunate but common reality that limits the long-term survey of planktonic communities is the high cost and erratic funding situations associated with marine research. The significant resources involved in maintaining quality oceanographic measurements often cripple the ability to fund studies through time. Over 100 years ago, innovative strategies such as Hardy’s continuous plankton recorder, using ships of opportunity over dedicated scientific fleets, brought great success and incredible amounts of data by using existing networks of different surface ships (“The Continuous Plankton Recorder: Concepts and History, from Plankton Indicator to Undulating Recorders” 2003). These instruments are still delicate, specialized, and expensive however, limiting their deployment as well as the compilation and interpretation of the data. However, almost 20,000 sailors a year utilize the wind as a way to explore the oceans. We can incorporate the enthusiasm and reach of this community into an experimental design that involves explorers to take data throughout their journey with a low foot-print, enabling a green participatory science. Current trends in affordable electronics and distributed manufacturing make it possible to put data collection in the hands of thousands of users across the planet. Computer vision and automated image processing make cataloging and storing massive amounts of data more straightforward than ever before. Most importantly, the globalization of scientific networks makes it possible for motivated researchers spread throughout the world to share inspiration, technical advice, and support.

### In-situ plankton monitoring

As demonstrated in the data from the Comau fjord, plankton dynamics in many ecosystems are subject to rapid change in response to climate events and human activity. In Chile, seasonal blooms cause frequent damage to local fisheries and the communities that depend on them. Single blooms have created losses of over 800 million dollars locally, as well as contributing to a growing public health crisis. Tools such as the Planktonscope could be deployed on a large scale. Correlation with other geochemical data collected from coupled sensors could act as automated plankton monitors, serving as sentinels against unexpected harmful algal blooms.

### Continual culture monitoring

Cultivation of marine cultures is notoriously challenging and in many cases infeasible for extended periods. However valuable behavioral and morphological data can be collected through continuous visual monitoring for even limited periods of time. Thousands of images, with millions of features, could help identify diel behavior, or characterize cell or life-cycle dependent morphological changes. Our limited data set of diatom and dinoflagellate cultures demonstrates how a visual time series performed in the laboratory could bring insight or depth to behavioral or ecological studies.

Several studies have displayed the unique power of visual inspection of planktonic communities throughout time(Brownlee, Olson, and Sosik 2016). We hope tools such as the Planktonscope will help expand this detailed exploration of our oceans by equipping and uniting more researchers around a platform that evolves with each new question. A tool simple and affordable enough to be made anywhere, yet accurate and reliable enough to provide quality scientific data and is supported by an open source scientific community via updates and customization due modular design could enable global communities to participate in oceanographic research.

## Materials and methods

### Prior development

The proof of concept of this instrument was a monolithic fluidic microscope made of Thorlab components encapsulated in a metallic box. It was capable of continuously imaging a liquid sample under bright-field thanks to the coupled Raspberry Pi/Pi Camera, 2 M12 Lenses, and integrated fluidic path. The liquid sample was pulled through the field of view of an ibidi channel slide via a peristaltic pump. This prototype was given to a French sailor named Guirec Soudée to use on board his sailboat during his return journey to Bretagne, France from San Francisco California. By collaborating with the Fablab UBO Open Factory, this original version has been completely reinvented to serve the challenge of modularity in a microscope.

### Design thinking

Because the original modular design is made of different units that can be stacked on top of each other, we decided to utilize a parametric design to optimize a physical interface common to all modules using Autodesk Fusion 360 (v2.0.5688). Different parameters define the dedicated areas of the interface such as the electronic connection area, the magnetic linkage, and the optical path. The thickness of the material plays an important role to characterize the interface as well as the outer dimensions of all the electronics used inside the instrument. This shareable online 3D environment provided by Fusion 360 contains the main 3D model as well as other models that form the electronic and optical parts. Most of these models have been generated by measuring the existing objects but some have been downloaded from an online library named GrabCad. Once the different iterations of the 3D model were ready to be machined, the sketches were extracted as DXF from Fusion 360 and nested in Adobe Illustrator CC (version 22.1) in order to fit the dimensions of the sheets of used material. The parts are then machined on a 3 mm thick acrylic by a laser cutter machine (RS-1610L) at UBO Open Factory. The laser cutter machine has an optimal resolution of 25 µm. The assembly was performed manually and took ∼5 hours.

### Content of the modules

The price of the material to assemble one unique Planktonscope V.1 is about $200. The Planktonscope V.2 is about $400 (Table S1).

#### Power, computational and sensor modules

The PlanktonScope is directly powered through one multi-functional module dedicated to the computation and sensor. It receives 12V either by a regular AC power adapter for lab experiments or a battery for field deployment.

For the Planktonscope V.1, a custom BUS made of 6 electronic wires dedicated to power the other modules provides 12V, 5V and Ground wires. The three other wires consist of the I2C, SDA, SCL and a dedicated Ground enabling the exchange of data between the different modules. The camera sensor is a Pi Camera v2.1 embedded in the module. It is positioned facing up in order to collect the image coming from above. For V.1, under this module and on one side are 3 suction cups allowing an improvement of the vertical and horizontal stability of the microscope in the field.

#### Optical modules

The optical train is defined by two inverted S-mount lenses (M12 lenses) that are both encapsulated in different detachable modules. The two modules have been designed to enable a rapid change of each M12 lens used as a couple. The alignment is set by the insertion holes cut and positioned by the laser cutter machine. The distances of the M12 lenses to each other and to the sensor are defined by rotating the M12 lenses in the holes tapped using a M12×0.5 hand thread tap from Thorlabs.

#### Stage and focus modules

This module is made to focus and explore a sample using a linear delta design largely documented thanks to the evolution of 3D printers. Three vertical Allegro A3967-based Easy-driver stepper motors are each equipped with two M3 diagonal push rods that all maintain a central platform. This platform is made of two horizontal parts that are magnetically coupled and can be separated in order to insert different types of sample mounts such as a glass slide, petri dish, or flow cell. Each stepper is driven by a A4988 driver powered with 9V and controlled by a common Arduino mini pro present in the module. In order to control the location of the sample maintained by the platform, an inverse kinematic is necessary to transform a X/Y/Z desired displacement in a delta motion. In this paper, the code embedded in the Arduino has been simplified to control only the focus by moving the three stepper motors simultaneously. This Arduino has a defined I2C address allowing the Raspberry Pi to iteratively set a new focal position.

#### Illumination module

The illumination is made of individual white LEDs 5 mm LTW2S - 17000 mcd forming five concentric rings. From the center to the external ring, the number of LEDs used is reciprocally 1, 6, 12, 18 and 24. Since the LEDs aren’t individually driven, each ring of LEDs are controlled separately by an Arduino Mini Pro with a defined I2C address. The Arduino is capable of applying the value of the analog intensities for each ring by receiving a setting from the Raspberry Pi. To simplify the wiring in this module, a custom PCB has been created by Eagle Version 9.0.1. The flow-through version utilizes a single LED for illumination.

#### Pump module

The pump for passing the sample through the field of view within a flow cell is encapsulated in a single module. This peristaltic pump is made by stacking several layers of laser cut acrylic. Through this process a peristaltic pump is formed by allowing 3 rollers connected to a stepper motor to compress a 3 mm neoprene tube. The use of laser cut parts allows the user to change the diameter of the tube used for compression in the pump.

[Supplementary figure]

### Optical characterization

The fact that the two lenses are detachable and easily swappable allows us to describe each configuration by imaging the USAF 1951 resolution target with different pairs of M12 lenses. By selecting five different M12 lenses, we collected images of the resolution target using 25 possible configurations. The illumination was set to use only the LED in the central ring for basic bright-field. The PiCamera was set to take a picture with 1080p corresponding to 1920×1080px. The different M12 lenses used to image were selected based on their similarities in terms of sensor format set to 1.5”, their optical resolution of 5 Mp, and their IR filter. The variable parameter used to determine the different configurations is then set only by their focal length (Table 3).

For a single configuration, a ruler was imaged in order to back calculate, via ImageJ, the actual size of the field of view along with the optical magnification for each pair of M12 lenses. The pixel resolution of each optical configuration was then calculated from the size of the field of view and the width in pixels of the image.

The resolution of each configuration has been measured by finding the smallest separable groups and elements from the resolution target. The resolution was calculated as follows :

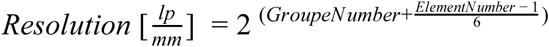

We used the following equation to retrieve the resolution in microns (µm) :

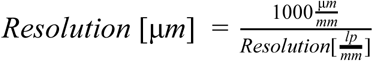

In order to determine the appropriate fraction size appropriate for each optical configuration, we divided the height of the field of view by 3 for the upper limit and consider a minimum equivalent diameter of 32px for the lower limit.

### Image acquisition and image processing

The physical configuration of the PlanktonScope remained the same for the different experiments presented in this paper.

For the optical configuration, the pair of the M12 lenses are 16 mm for the tube lens and 12 mm for the objective lens. The field of view measures 2 879 µm wide and 1 619 µm high. For the fluidic path, the peristaltic pump is set at a flow-rate of 3 ml/min and the flow-cell is a rectangle shaped borosilicate glass capillary (VitroTubes), 5 mm wide, 500 µm deep and 5 cm long. The volume imaged in one frame is about 2.33 µL, while the whole volume passed in the capillary is about 4.05 µL.

The processing of the images from V.1 was done with a desktop computer in order to process the data faster. For V.2, the processing was done on the Pi. For each acquisition of 2000 frames, a cleaning step was used to remove particles attached to the flow-cell while also correcting for chromatic aberration. A rolling ball algorithm using 20 frames around the considered frame (10 frames before and 10 frames after) was implemented.

From the cleaned frames, different image operations such as dilation, closing, and erosion were performed using OpenCV (version 4.0.0). Frames were converted into binary images from which we select and crop the region of interest corresponding to the objects. Different features were extracted from the binary extracted objects using Scikit-image (version 0.16.1) such as the eccentricity, the equivalent diameter, the euler number, the extent, the area, the filled area, the major axis length, minor axis length, the orientation, the perimeter, and the solidity.

### Monitoring morphology of lab cultures

The culture of Pyrocystis noctiluca was established from the strain UTEX LB 2504 and cultured in F/2 medium (Bigelow Labs) at 20C while maintaining a 12/12H light/dark cycle.

The culture of Coscinodiscus wailesii was established from the strain CCMP2513 and cultured in L1 medium (Bigelow Labs) at 20C while maintaining a 12/12H light/dark cycle.

### International field sites

The instrument has been deployed at six locations representing different ecosystems. The same sampling protocol (except for the Palo Alto Baylands Nature Preserve, see below) was performed using a 20 µm mesh plankton net with a diameter of 30 cm and a 10 minutes pull at 2 knots. All tows were surface tows. The samples were filtered with a mesh sieve to remove the particles bigger than 500 µm. For the Palo Alto Baylands Nature Preserve, since the site is shallow and quite turbid, the sample was collected directly using a 50mL falcon tube from the subsurface and filtered using the 500 µm mesh sieve. All samples were collected during daytime.

The acquisition was done using a frame-rate set at 8 frames/second, which corresponds to a volume of 1.12 ml imaged per minute. We took 2000 frames per sample, at 5 minutes total, the volume imaged was 5.6 mL per sample. From all the segmented objects, we manually selected the objects most likely to correspond to living organisms in order to avoid terrigenous sediment as all the field sites are coastal.

### Vertical stratification of planktonic communities in Comau Fjord

To investigate the diversity and distribution of plankton in the stratified fjord waters we employed a horizontal water sampler from LaMotte (CODE 1087) to sample the vertical distribution of plankton from 0-10 meters below the water’s surface. From the 1,200 mL samples collected for every depth, we conserved 15ml for flow-through imaging. The salinity at every depth was measured using a hand refractometer from Atago.

For the acquisition, the frame-rate was set at 8 frames/second and 2000 frames were taken for each depth corresponding to 5.6 mL imaged per depth.

## Supporting information

Planktonscope construction manual

## Acknowledgements

We acknowledge all members of PrakashLab for insightful comments and suggestions throughout this work. We thank Jorge Mardones and Lara Zamora (IFOP, Chile) and Gurudeep Rastogi (Wetland Research and Training Center, Chilika Development Authority) for support during field testing. We acknowledge Rainer Kiko for introducing us to MorphoCut (Github https://github.com/morphocut/morphocut). A.G.L. is a Simons Fellow supported by the Simons Foundation. M.P. acknowledges funding support from Schmidt Futures, Moore Foundation and NSF Center for Cellular Construction NSF STC award DBI 1548297, CZ BioHub and HHMI-Gates Faculty Fellows Program.

## Supplementary Material

**FIGURE S1:**
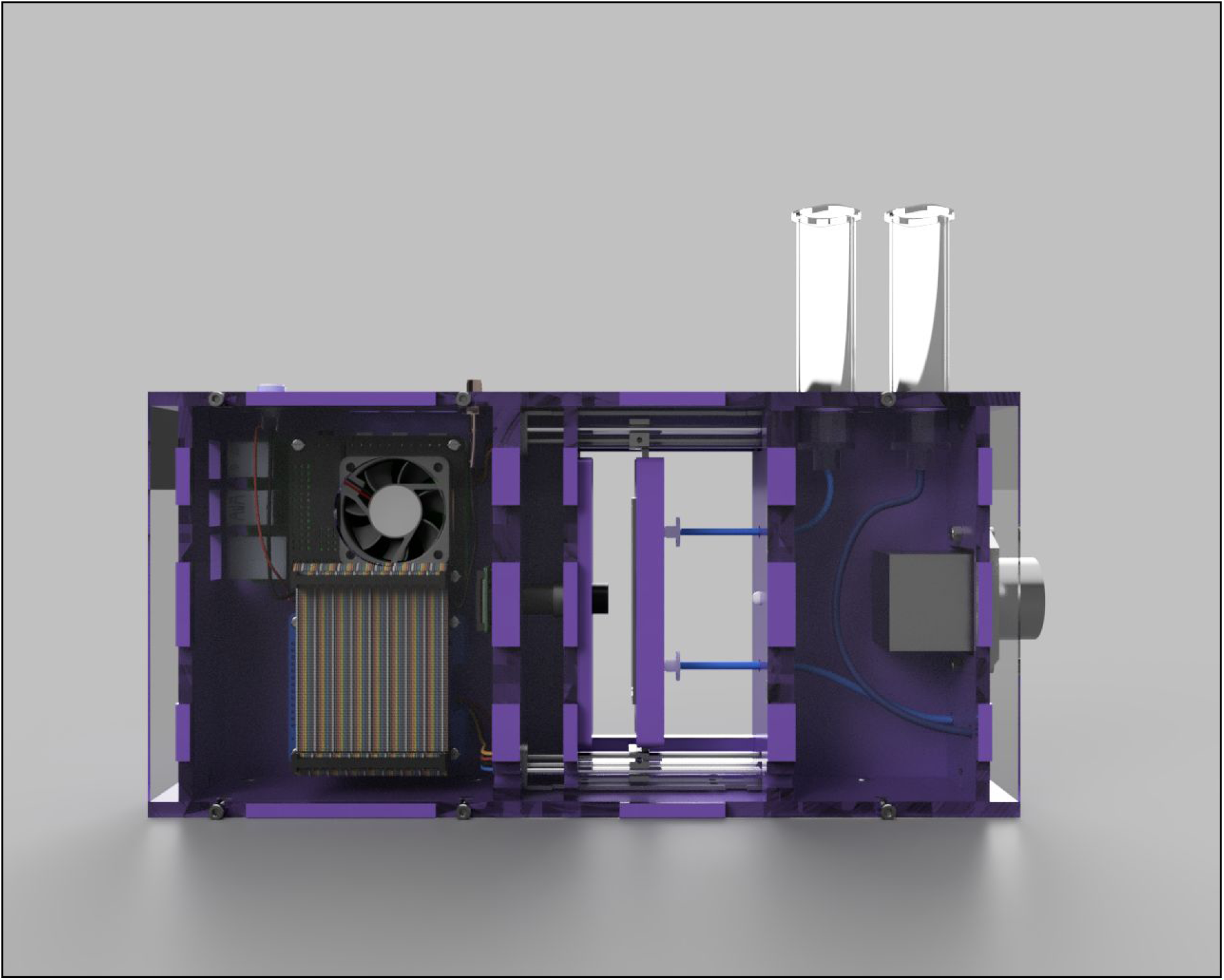
CAD. The design for the Planktonscope is available for download on the Planktonscope.org website or through the Planktonscope Github repository. Detailed walkthroughs for assembly and other resources are also available.

**FIGURE S2:**
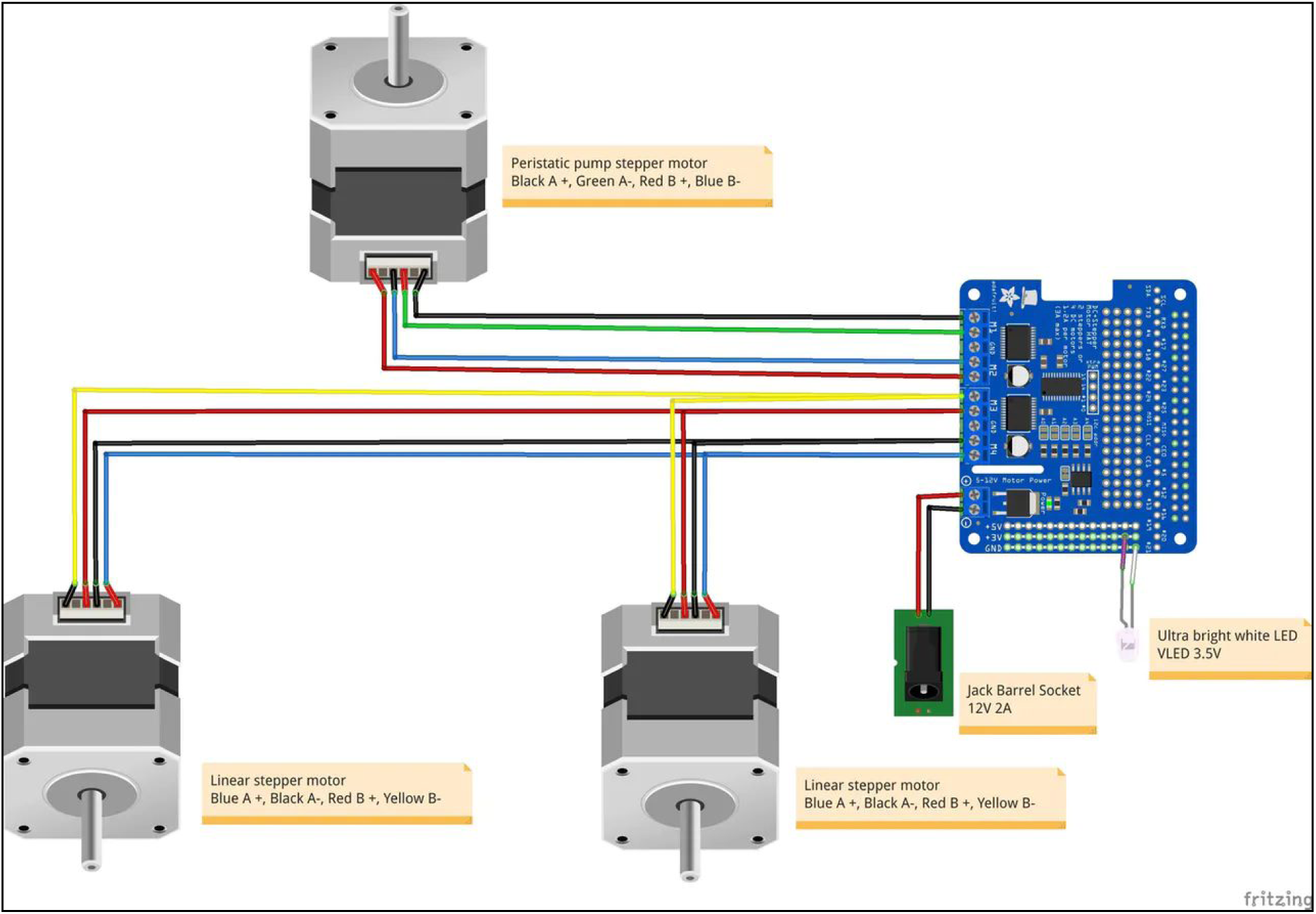
Electronic layout. The current wiring diagram for the Planktonscope. More technical details and guides are available on the Planktonscope.org website.

**FIGURE S3:**
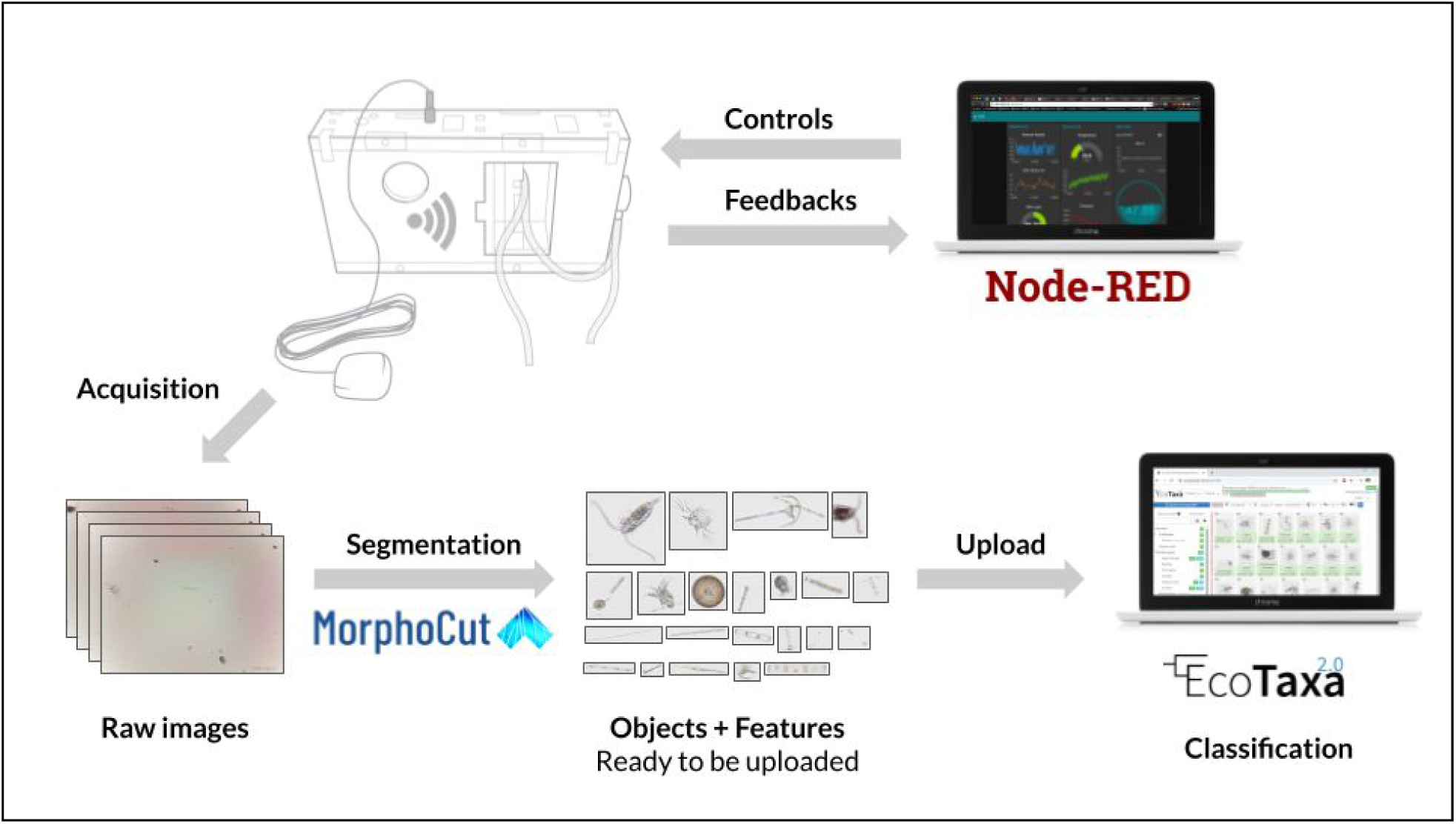
Software architecture. The current software for running the Planktonscope consists of Python scripts for managing hardware and acquisition controls, all accessible via a Node-RED dashboard transmitted via a Wifi signal from the Raspberry Pi. The images are then background subtracted, segmented, and extracted with Morphocut (https://github.com/morphocut/morphocut) libraries then uploaded to EcoTaxa for classification.

**FIGURE S4:**
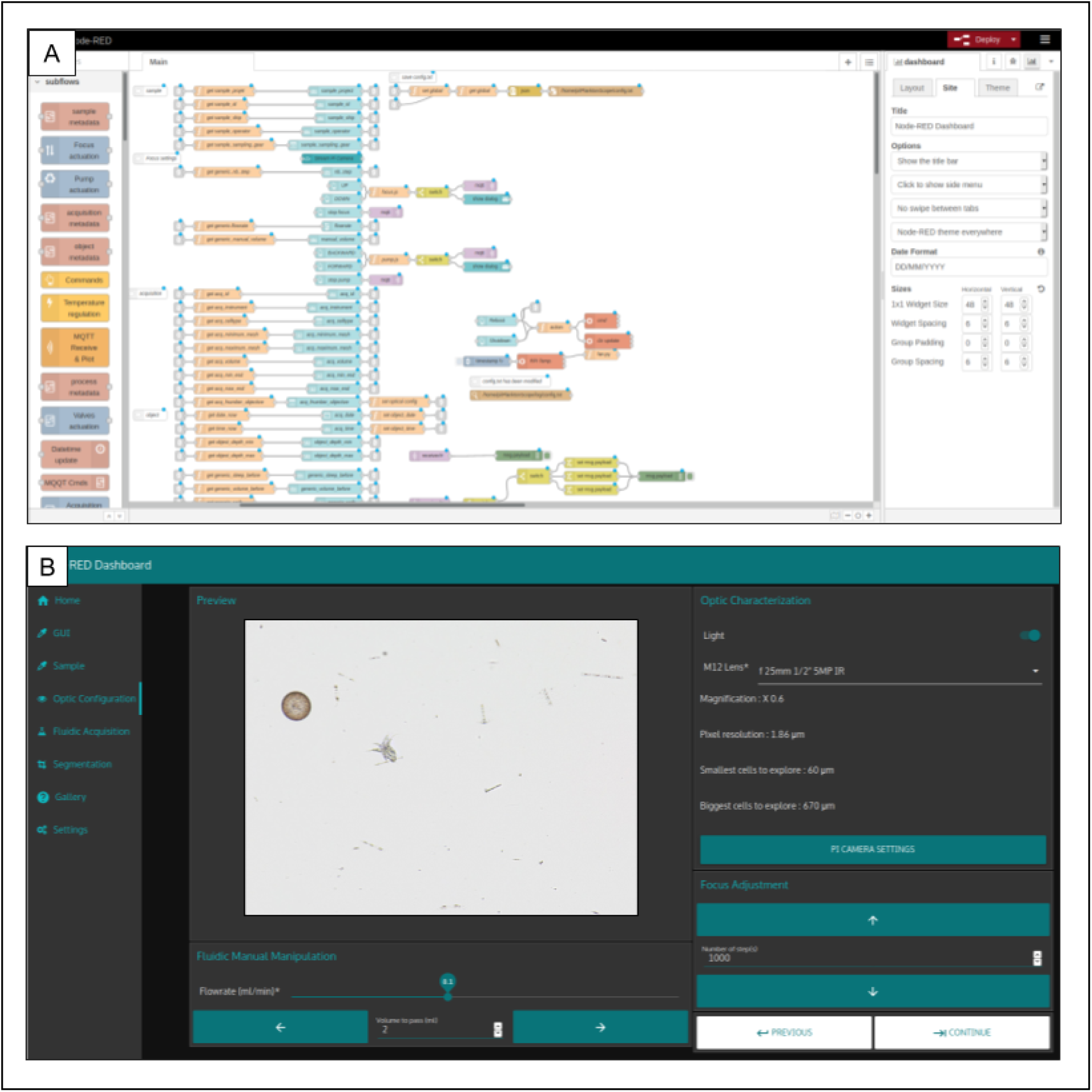
Node-RED modular architecture and software GUI. Snapshot of Node-RED programming and a GUI used to control Planktonscope in real time via any browser; including desktop PCs or a cellphone.

**FIGURE S5:**
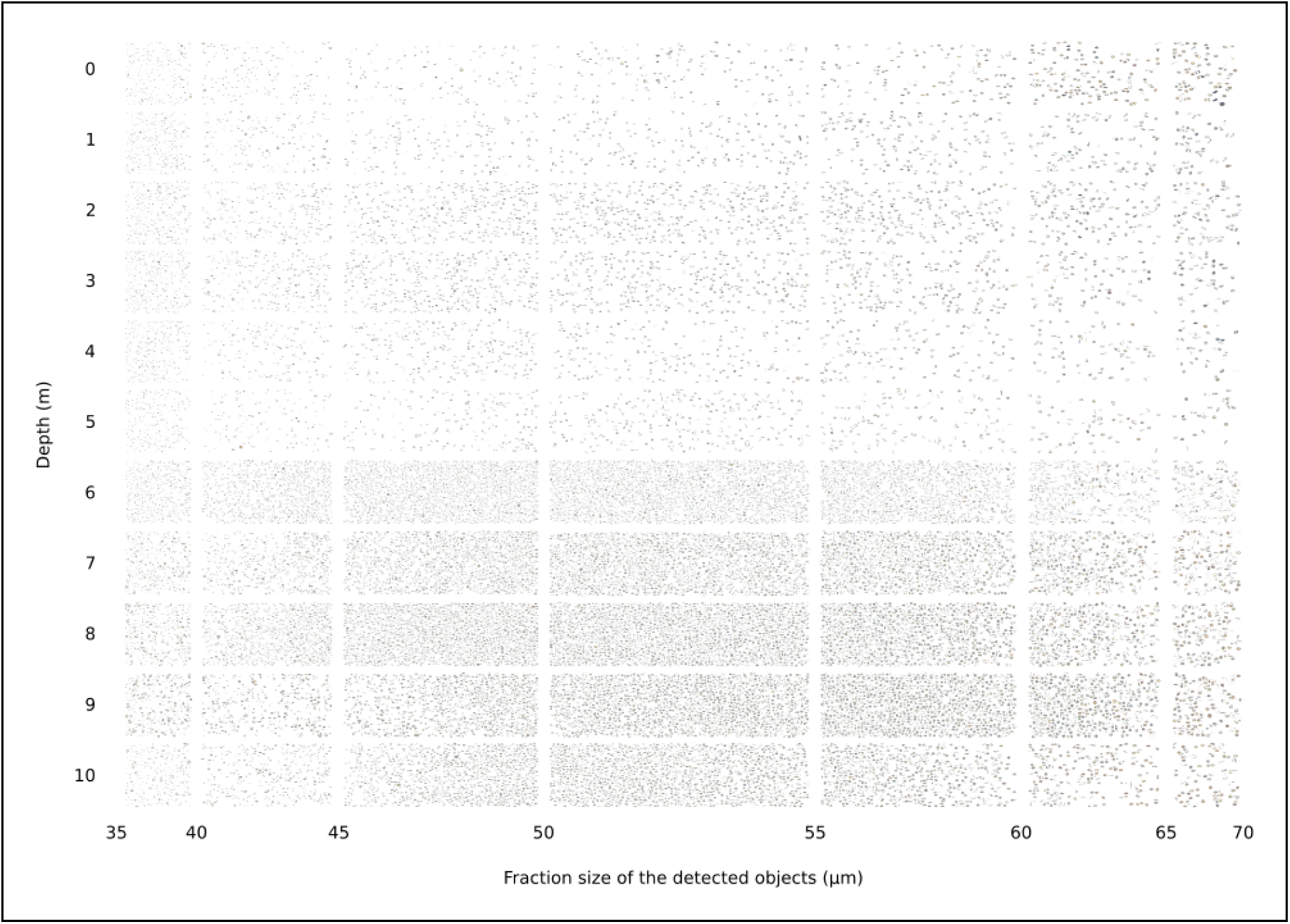
Collage of the detected objects for each depth (from 0 to 10 meter) in function of the fraction size (from 35 to 70µm) The data from the experiment in the Chilean fjord consisted of thousands of images of plankton with different sizes and features changing as a function of depth. Using a collage with actual images of planktonic organisms as data points serves as a tool to display qualitative and quantitative information in one figure.

## SUPPLEMENTARY MATERIAL 1 : Description of field sites

### Palo Alto Baylands Nature Preserve

The **Palo Alto Baylands Nature Preserve** is the largest tract of undisturbed marshland remaining in the San Francisco Bay (Rosso, Ustin, and Hastings 2005). As a bay salt marsh, the field site is a unique mixture of tidal and freshwater habitats. The sample site is quite shallow, about 20cm on average, hosting many large ciliates such as Tracheloraphis, Enchelyodon and Heterotrichs. Among them, we detected diatoms *Amphiprora gigantea*, Gyrosigma sp. and different penne diatoms (Fig. 4A).

### Monterey Bay

Located on the Central Coast of California, **Monterey Bay** is renowned for its complex habitat, diverse seafloor, deep submarine canyons, and a strong seasonal upwelling that creates a constantly evolving ecosystem. We observed different species of pennate diatoms and identified many as *Odontella longicruris*. Additionally, we can observe other dinoflagellates, armored or in resting cysts (Fig. 4B).

### Isla Secas

The **Isla Secas** (Dry Islands) is a small archipelago in the Gulf of Chiriqui on the Pacific coast of Panama consisting of about sixteen islands. This coastal archipelago offers seasonal sanctuary to two different populations of migrating humpback whales. The region was largely populated with zooplankton such as copepods, crustacean larvae, as well as diatoms (*Nitzschia longissima*) (Fig. 4C).

### Comau Fjord

The **Comau Fjord** is surrounded by mountains reaching an altitude of 2,000 meters, and water of maximum depth around 500 meters. This unique geological zone receives a large amount of rainfall distributed throughout the year (5000 mm/year). This fjord is a notable example of seasonal harmful algal blooms ((León-Muñoz et al. 2018)). We detected many diatoms such as (*Ditylum brightwellii, Detonula pumila*), many small dinoflagellates (Gyrodinium spp., Protoperidinium sp., Dinophysis sp., *Lepidodinium chlorophorum, Prorocentrum compressum*, Pronoctiluca sp.) and at least one species of ciliate (Fig. 4D).

### Isla Magueyes

**Isla Magueyes** is home to the second largest coral reef system in the world, as well as coastal mangroves which provide large amounts of nutrients to the waters, while also being in close proximity to deep water. Sampling the outer reef, we observed a typical zooplanktonic presence of copepods as well as its larvae. We determined the presence of many different diatoms (Coscinodiscus sp., Eucampia zodiacus, Chaetoceros spp and Proboscia alata), dinoflagellates represented by the genius Ceratium (Ceratium sp., Ceratium furca and Ceratium lineatum) and finally the presence of several specimens of Oscillatoria (Fig. 4E).

### Chilika Lake

As the largest coastal lagoon in India and the second largest coastal lagoon in the world, the **Chilika Lake** is a very precious and unique ecosystem. This gigantic brackish water lagoon is important for many reasons. It is a ground for migratory birds, home for several threatened species, and sustains more than 150,000 fisher–folk ((Sekhar 2007)). The water we sampled from the surface had large amounts of debris as well as Tintinnids and numerous larvae (crustacean and nauplius) (Fig. 4F).

## SUPPLEMENTARY MATERIAL 2 : Specifications of the PlanktonScope

**Optical magnification :** from **0.15X** to **2.36X**, we used **0.75X**

**Pixel size :** from **7.62**µ**m/px** to **0.48**µ**m/px**, we used **1.5**µ**m/px**

**Optical resolution :** from **15.63**µ**m** to **1.95**µ**m**, we used **2.76**µ**m**

**Travel distance** of the platform : **3.2cm** (with a step size of **0.15**µ**m**)

**Types of illumination** : **bright-field** & **dark-field**

**Maximum frame-rate** to avoid repeated objects : from **60fps** to **0.2fps**, we used **12fps**

**Nb frames per minute :** from **2334fpm** to **146fpm**, we used **741fpm**

**Volume imaged per min** : from **0.5ml** to **8.8ml**, we used **1.7ml** (for a flow-cell of **500um** thick)

**Flow-rate :** from **0.2ml/min** to **5ml/min**, used **3ml/min**

**SUPPLEMENTARY TABLE 1:** V.2 Bill of materials.

| Category | Quantity | Name | Details |
| --- | --- | --- | --- |
| Electronic | 1 | Raspberry Pi 4 B (4GB) | The single board computer from Raspberry Pi with 4GB of memory |
|  | 1 | Adafruit Stepper Motor HAT | This module controls 2 steppers: the focus and the pump stepper motors |
|  | 1 | Adafruit Ultimate GPS HAT | This module stores the date & time as well as logs GPS coordinates |
|  | 1 | Yahboom Cooling Fan HAT | This module cools and enables visual feedback for functions with LEDs |
| Electronic | 1 | Hammer Header Male 2x20 | Only one is needed |
|  | 1 | Stacking Header 2x20 | Only one is needed |
|  | 2 | Pitch IDC Sockets 2x20 | Only two are needed |
|  | 10cm | GPIO Ribbon IDC 40P | Only 10 cm is needed |
| Electronic | 1 | Flex Cable for Pi Camera | A small flex cable is included along with the Pi Camera however, a longer one >50cm is needed. |
|  | 1 | DC Power Jack Socket | Only per unit is needed |
|  | 1 | GPS Active Antenna | Make sure the IPEX to SMA cable is included |
|  | 1 | Micro HDMI Cable | This optional cable can be used for development purpose |
| Electronic | 1 | Power Supply 3A (USB) | Needs to provide 3A 5V |
|  | 1 | Power Supply 1A (USB) | Needs to provide 1A 5V |
|  | 1 | USB Type-C to USB-A 2.0 | To power the Raspberry Pi |
|  | 1 | USB 5v to DC 12v Step Up | Make sure this USB 5V / DC 12V step up converter limits the current at 0.8A |
| Electronic | 1 | Stepper Motor Peristaltic Pump | Only one is needed |
|  | 2 | Linear Stepper Motor | Make sure to select two linear steppers |
|  | 1 | MicroSD Card + Adapter | The minimum size is 32GB but can go up to 512GB |
|  | 1 kit | Heat sink kit for Raspberry Pi | Only one is needed |
| Fluidic | 1 kit | µ-Slide I Luer Variety Pack | Other flow cells can be mounted, but currently ibidi are used |
|  | 2 | HSW 20ml Syringe | Other syringes can be used |
|  | 1 kit | ÜberStrainer Set 3 | These help keeping the flow cell from being clogged |
|  | 1m | Silicone Tubing ID 1.6mm | Used for the fluidics |
| Fluidic | 2 | Luer Lock Connector Male 1.6 mm | These are used to secure the fluidic path |
|  | 2 | Luer Connector Male 1.6 mm | These are used to secure the fluidic path |
| Optic | 1 | LED white 5mm | A simple white LED will do |
|  | 1 kit | Arducam M12 Lens Kit | This is an affordable way to try many different optical configurations |
|  | 1 | M12 Lens 25mm IR 1/2" 5MP | This is not included in the Arducam kit, and is used as the tube lens |
|  | 1 | Pi Camera V2.1 | This will be the sensor |
| Hardware | 1 | M2.5 Standoff Assortment Kit | 6mm and 15mm standoffs |
|  | 1 | M2 M3 M4 Screw Assortment Kit | M2x8mm and M3x12mm screws and M3 nuts |
|  | 1 | CR1220 Battery | CR1220 battery that will be placed in the Adafruit Ultimate GPS HAT |
|  | 16 | Magnets 6 x 2 mm | Neodymium magnets to connect functional layers |

**SUPPLEMENTARY TABLE 2:** Characteristics of M12 lenses.

| f | IR | Nb of Mega Pixel | Size (inches) |
| --- | --- | --- | --- |
| 6 | Yes | 5 | 1/2.5 |
| 8 | Yes | 5 | 1/2.5 |
| 12 | Yes | 5 | 1/2.5 |
| 16 | Yes | 5 | 1/2.5 |
| 25 | Yes | 5 | 1/2 |

**SUPPLEMENTARY TABLE 3:** Size of the field of view.

| Size of the field of view<br>Width x height (mm) |  | f-number of the objective lens |  |  |  |  |
| --- | --- | --- | --- | --- | --- | --- |
|  |  | 6 | 8 | 12 | 16 | 25 |
| f-number of the tube lens | 6 | 6.36 x 3.58 | 4.79 x 2.69 | 3.25 x 1.83 | 2.50 x 1.41 | 1.56 x 0.88 |
|  | 8 | 8.31 x 4.68 | 6.44 x 3.62 | 4.27 x 2.40 | 3.32 x 1.87 | 2.06 x 1.16 |
|  | 12 | 12.25 x 6.89 | 9.37 x 5.27 | 6.36 x 3.58 | 4.92 x 2.77 | 3.07 x 1.73 |
|  | 16 | 15.62 x 8.79 | 11.69 x 6.58 | 7.90 x 4.44 | 6.19 x 3.48 | 3.90 x 2.19 |
|  | 25 | 25.00 x 14.06 | 18.68 x 10.51 | 12.41 x 6.98 | 9.80 x 5.51 | 6.09 x 3.43 |

## SUPPLEMENTARY FILES

**Assembly_Instructions.pdf**

