## Supplementary material for "PlanktonScope: Affordable modular imaging platform for citizen oceanography": Planktonscope construction manual

### PlanktonScope v.2

#### Instructions

Make sure you have your screwdriver kit, soldering iron, and components ready. Also, remember to flash the Planktonscope image disk on the SD card before installing the Raspberry Pi. If you are not familiar with any process, such as soldering, tapping, or wiring, try and read or watch a few introductory videos on the topics. Soldering deals with high heat and potentially toxic materials, so make sure to use the proper precautions.

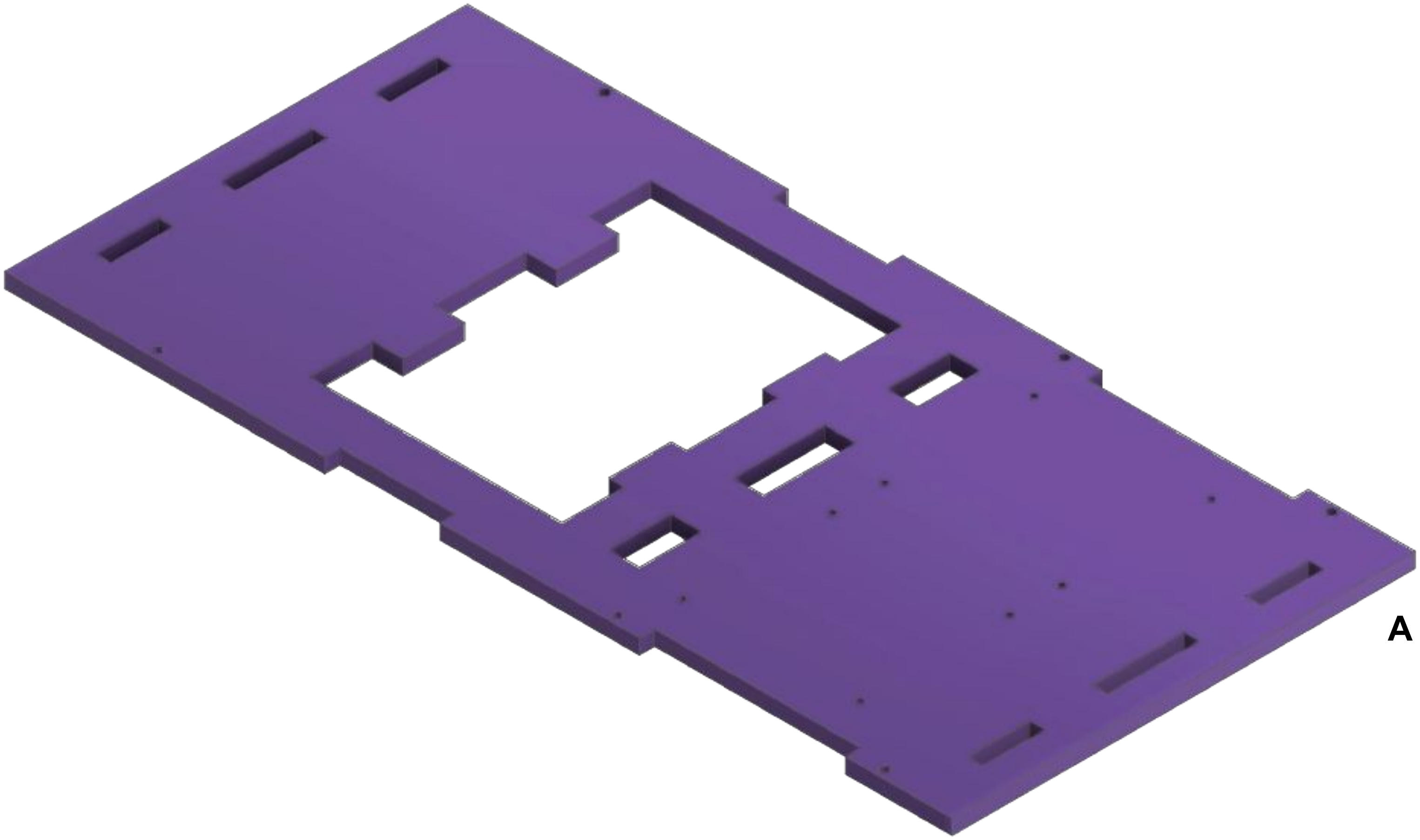


Laser cut all components using the .ai file ensuring all cuts are complete. The current design should have a **5mm** material thickness. Start by placing laser cut base **A** on a flat workspace.

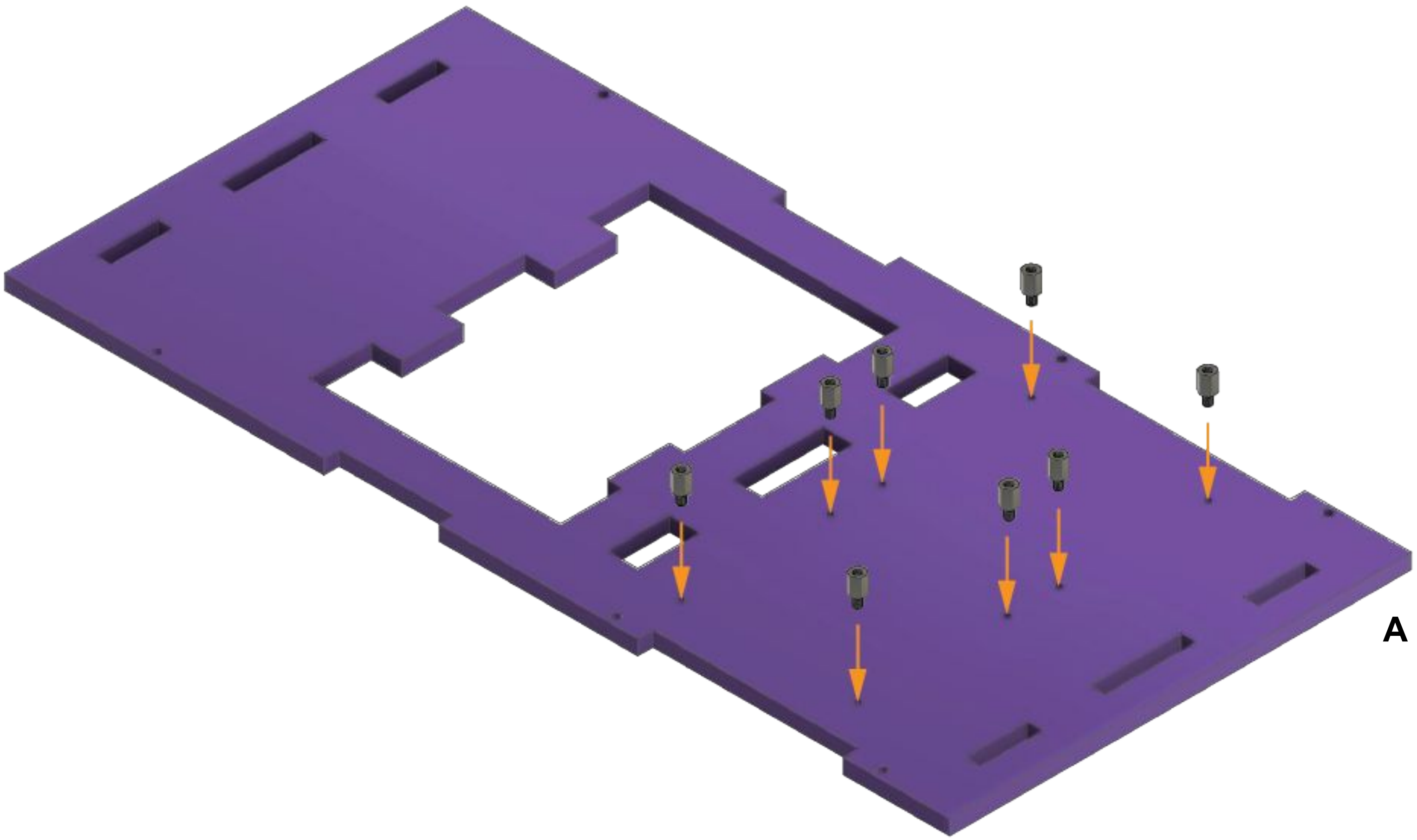


Place 8 standoffs (M2.5 6mm) into the designated holes on the laser-cut base **A.**

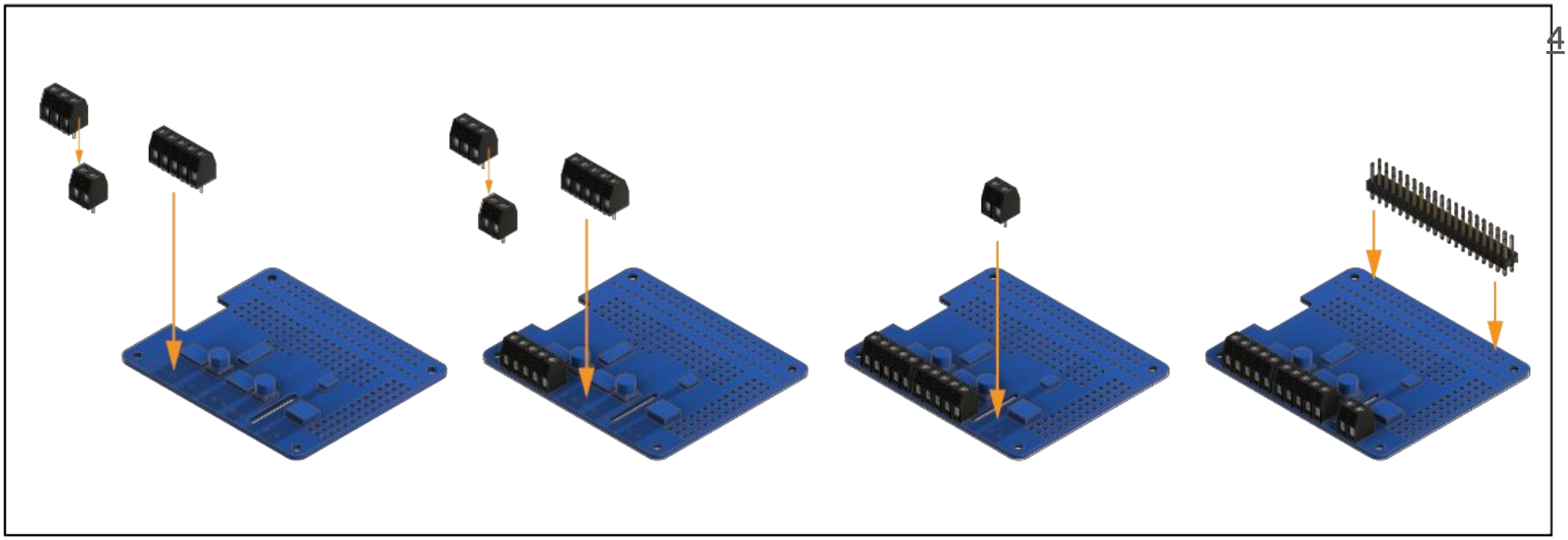

Insert and solder the terminal blocks and headers onto the motor driver PCB.

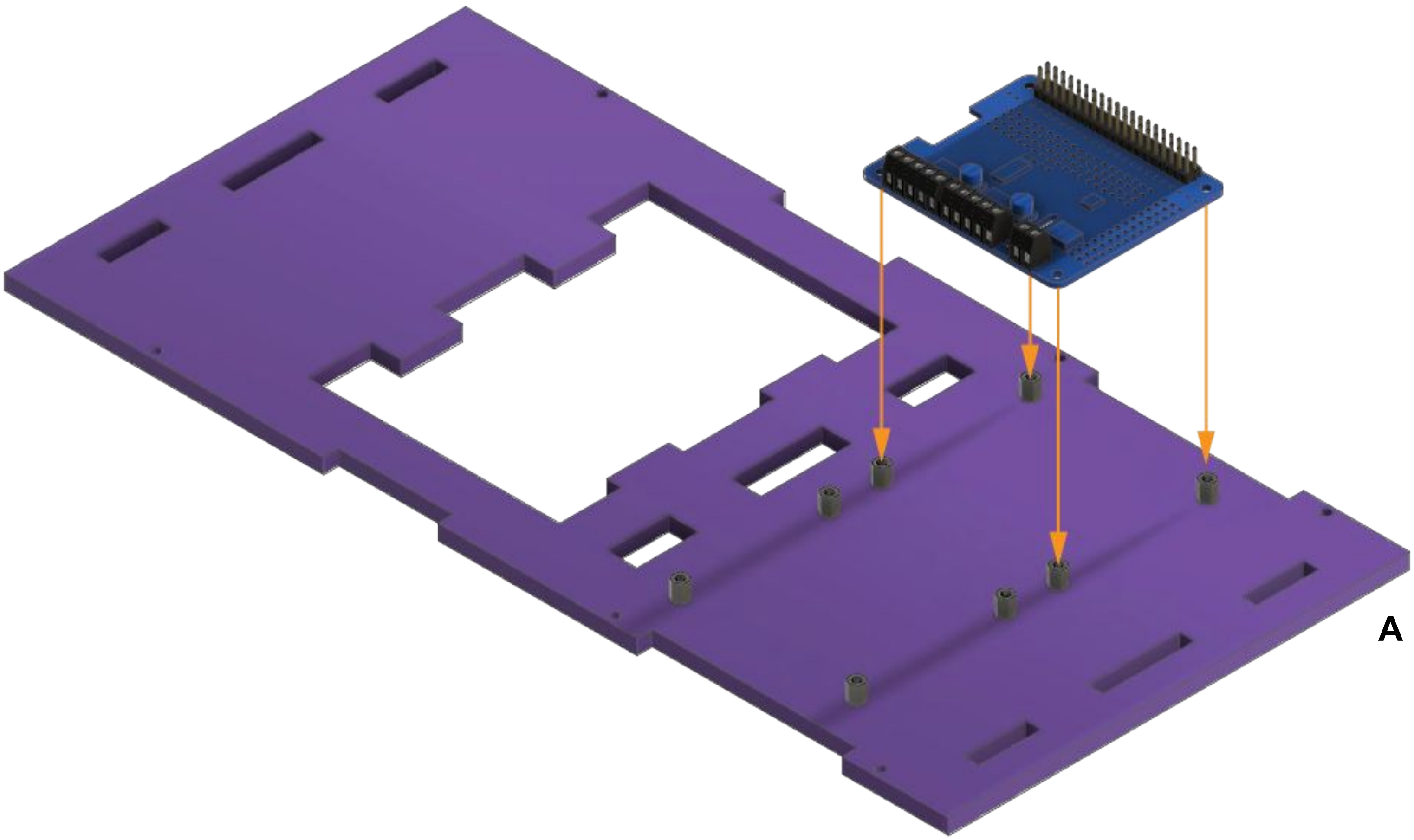


Place the motor driver PCB on to the indicated standoffs on **A**

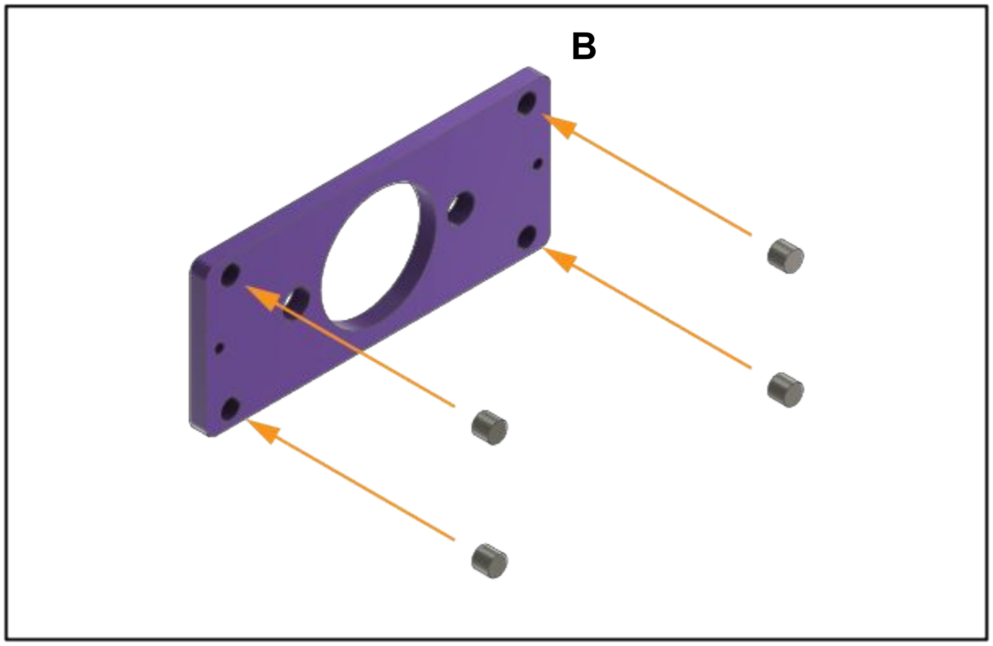


Insert the magnets into the laser cut sample stage **B**

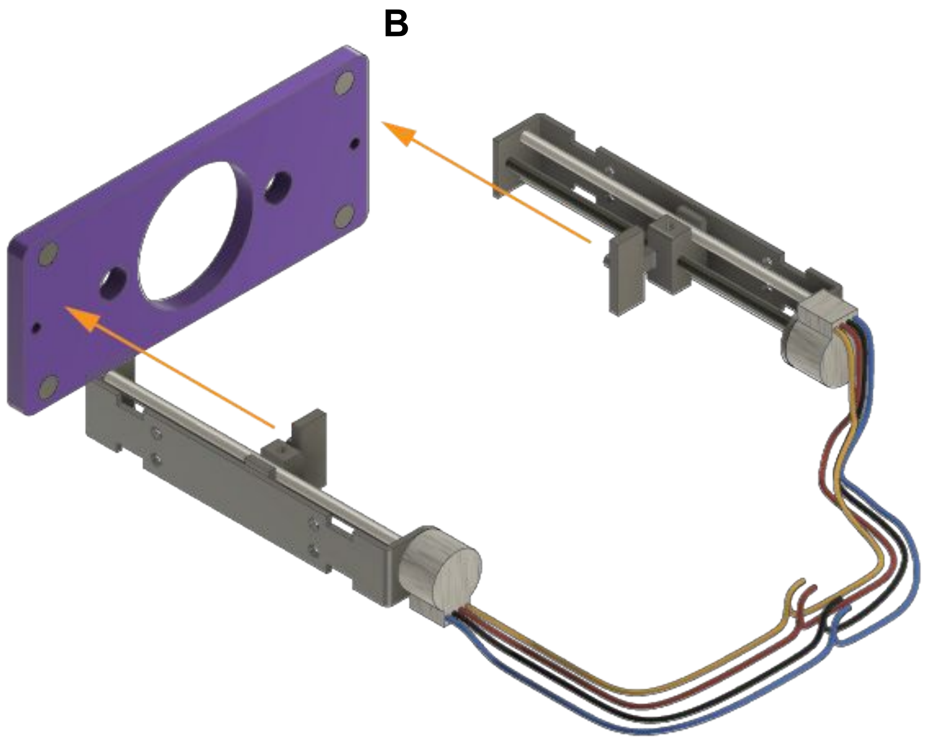


Connect the attachment points on the stepper motors to the sample stage **B.**
Secure with super glue or epoxy

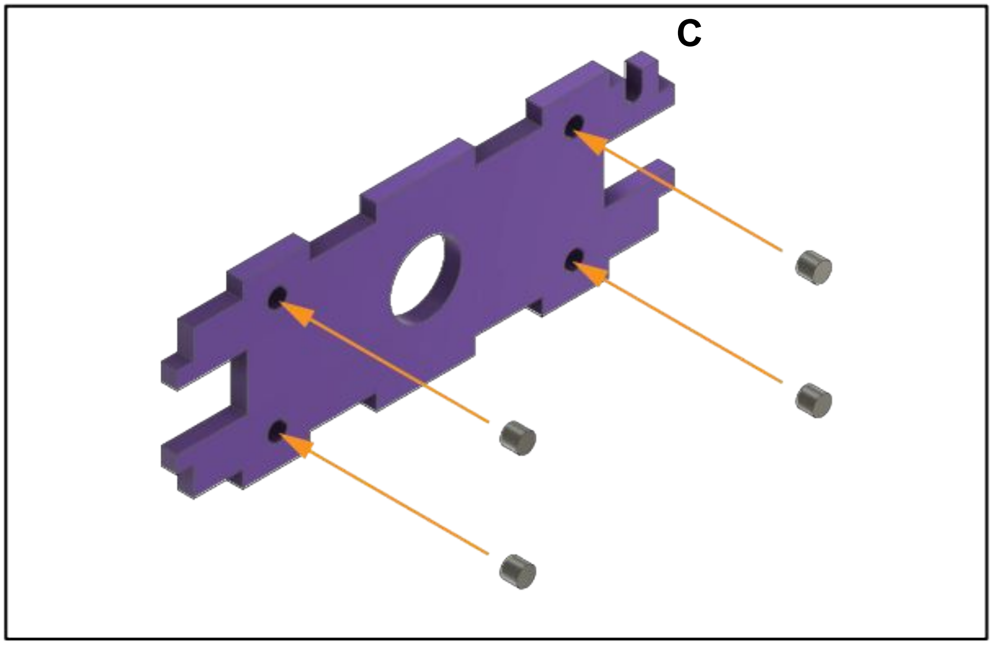


Insert the magnets into the laser cut mount **C**

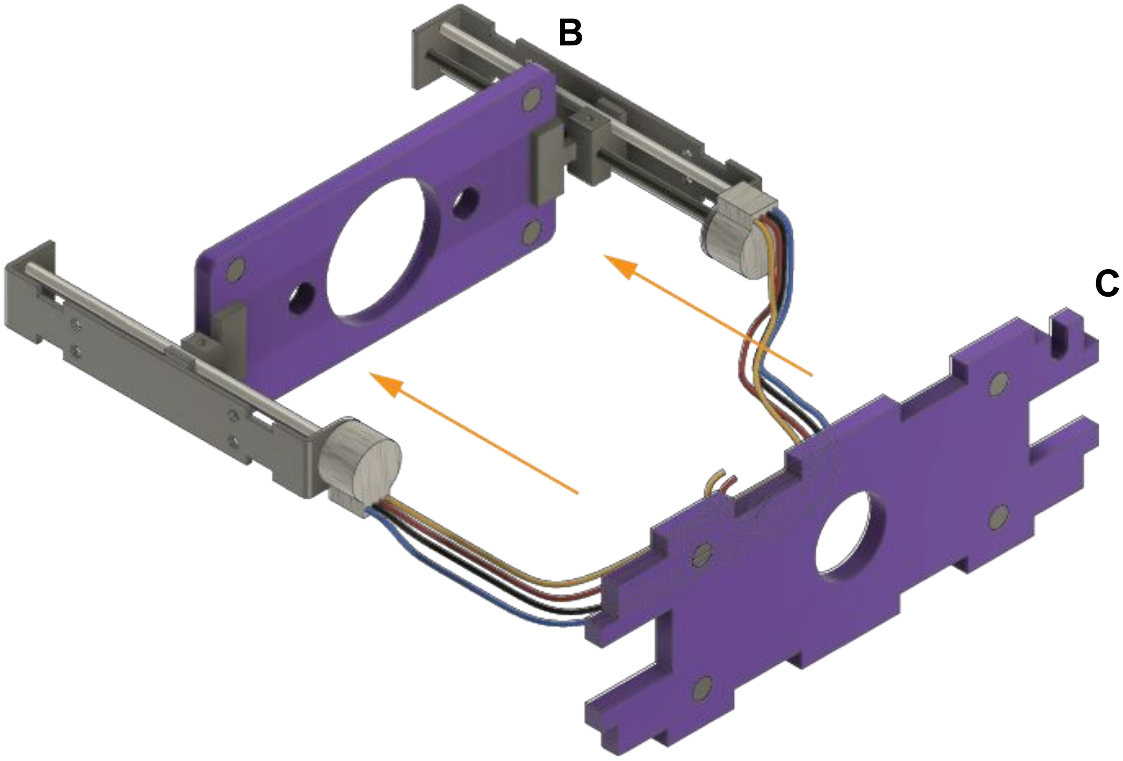


Arrange mount **C** as shown

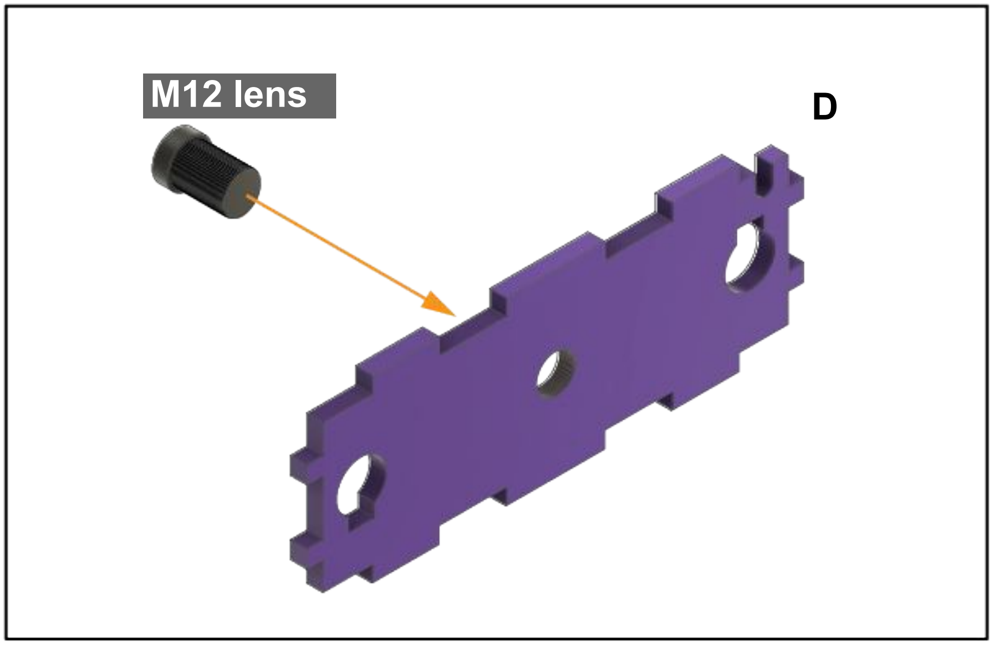


Tap the lens mount **D** with the 12mm tap*

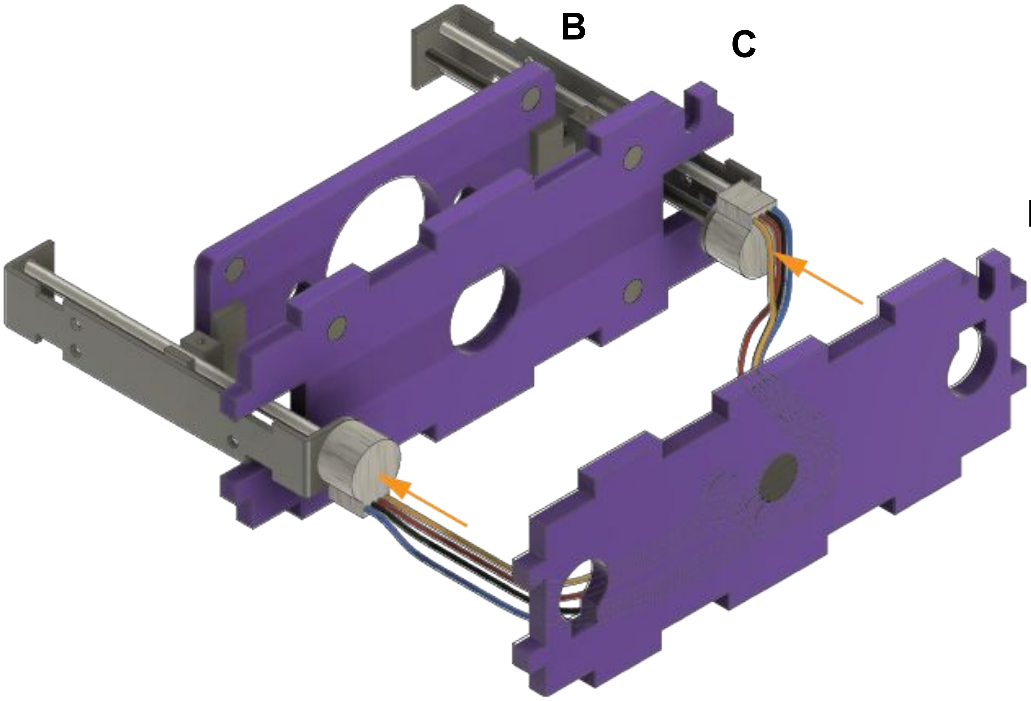


Insert the bases of the stepper motors into the laser cut lens  mount **D**

*the taps in the acrylic mounts will align and secure the m12 lenses and form the optical path.. Attention should be paid to tapping the pieces as straight as possible.

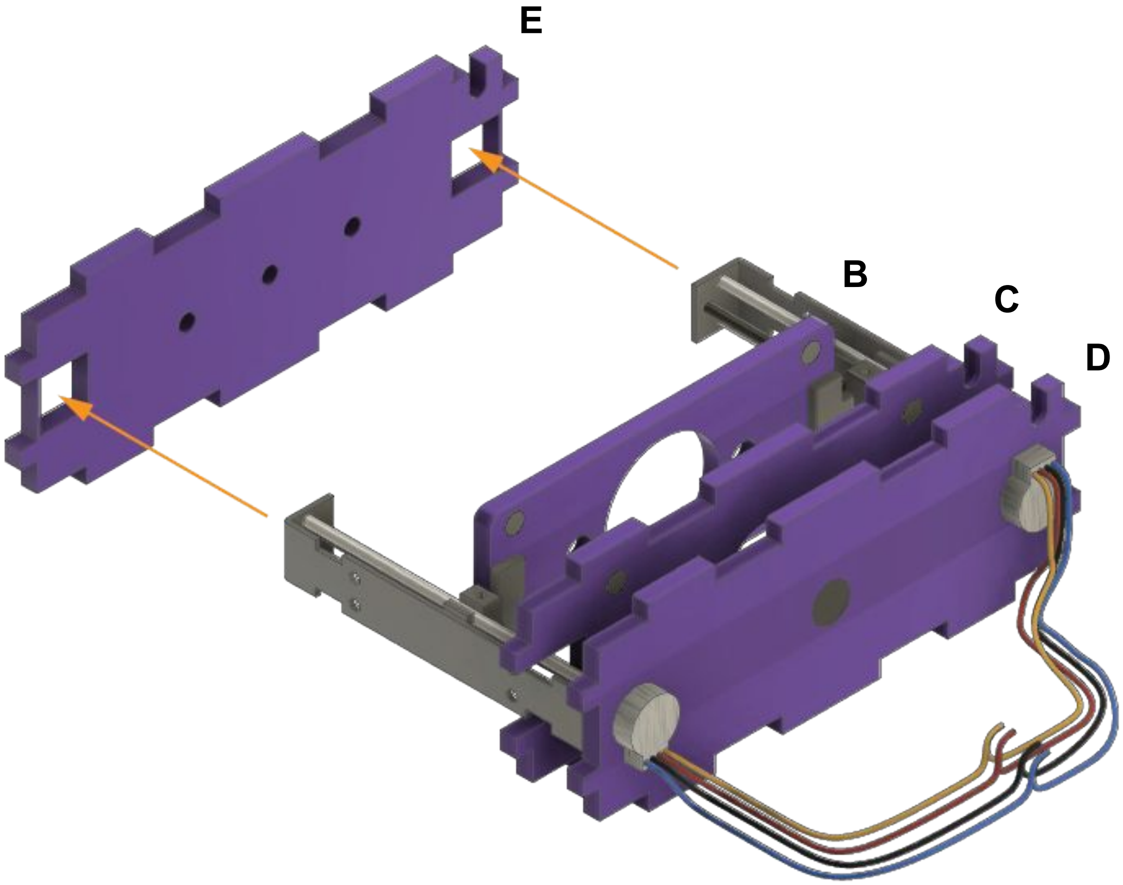


Insert the other end of the stepper motor driver into its laser cut mount **E**

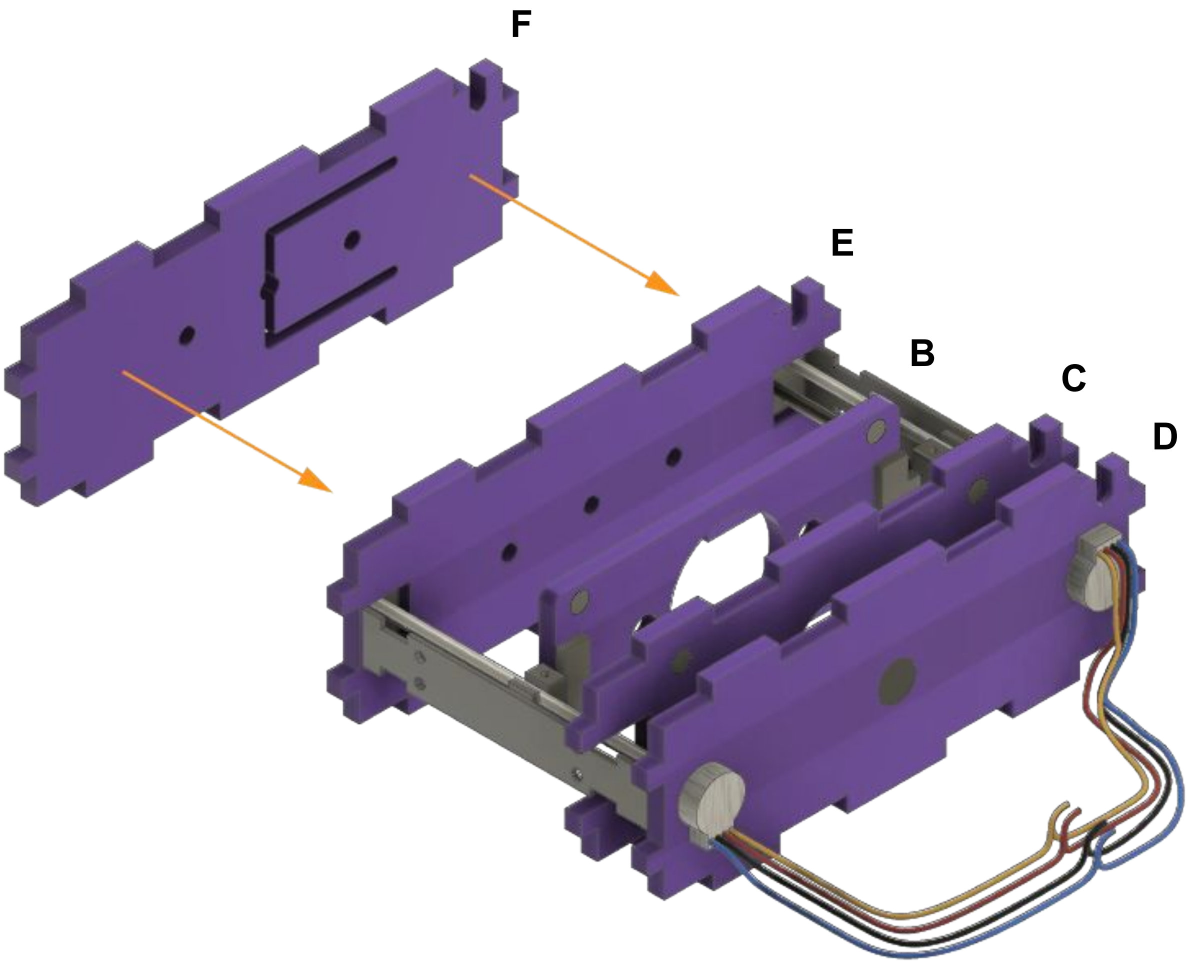


Attach the illumination mount **F,** securing with acrylic cement or superglue

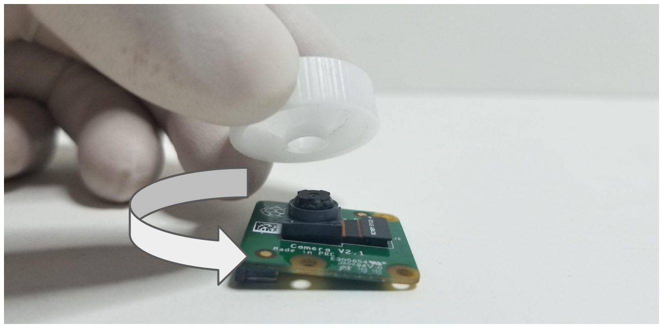

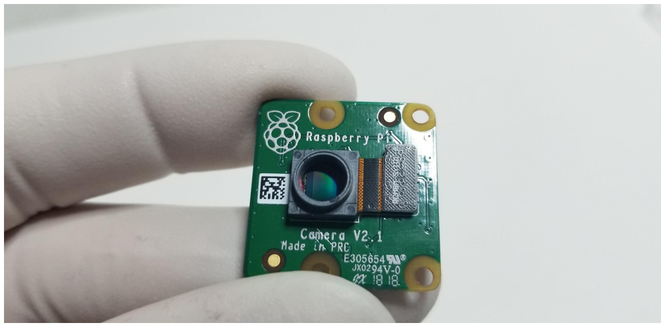

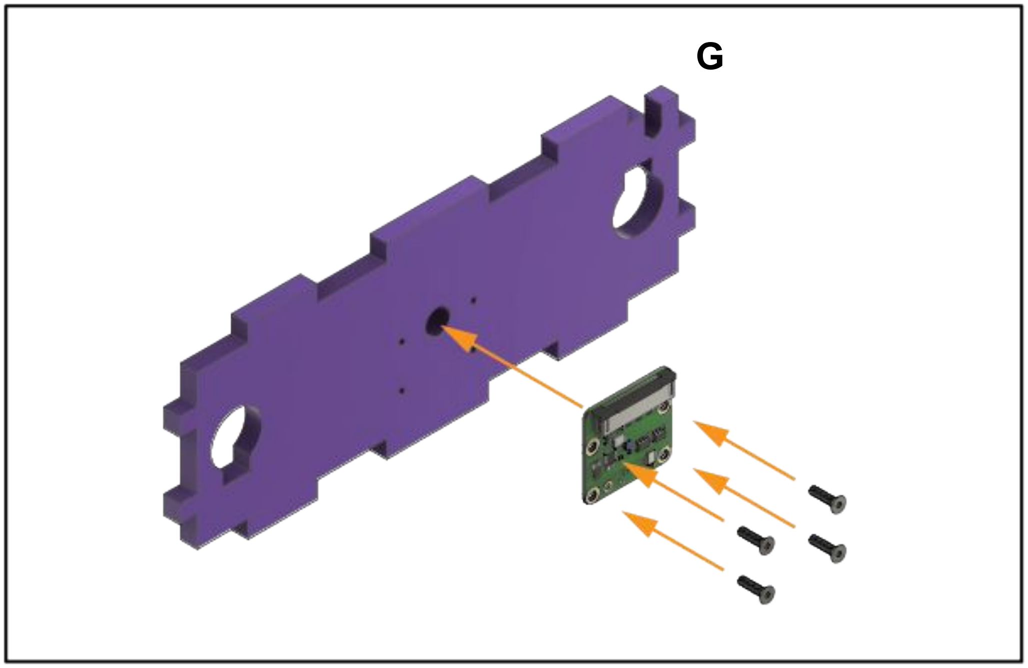


Carefully remove the lens from the PiCam and insert the camera into the laser cut mount being careful to avoid getting oil or dust on the sensor

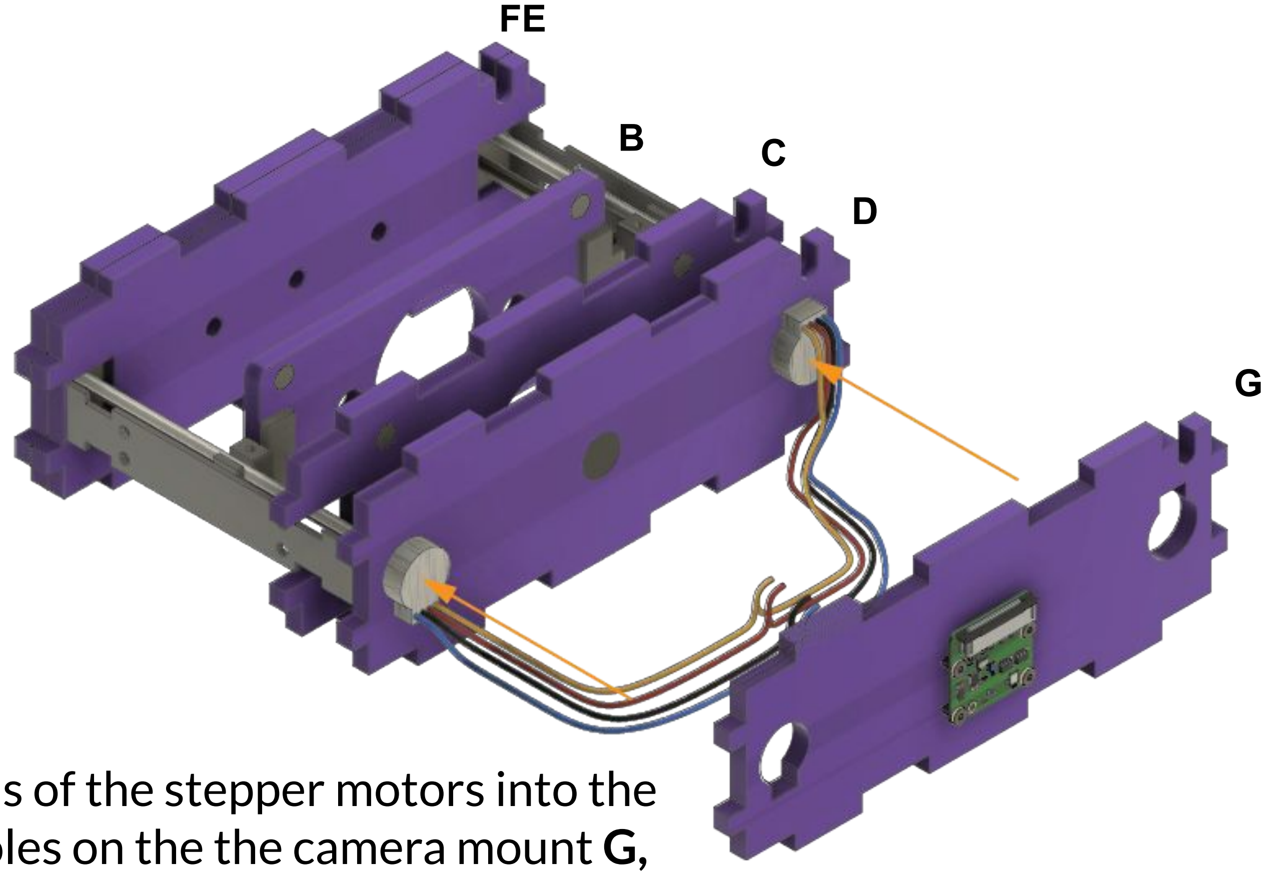


Insert the ends of the stepper motors into the designated holes on the the camera mount **G,** first feeding through the wires

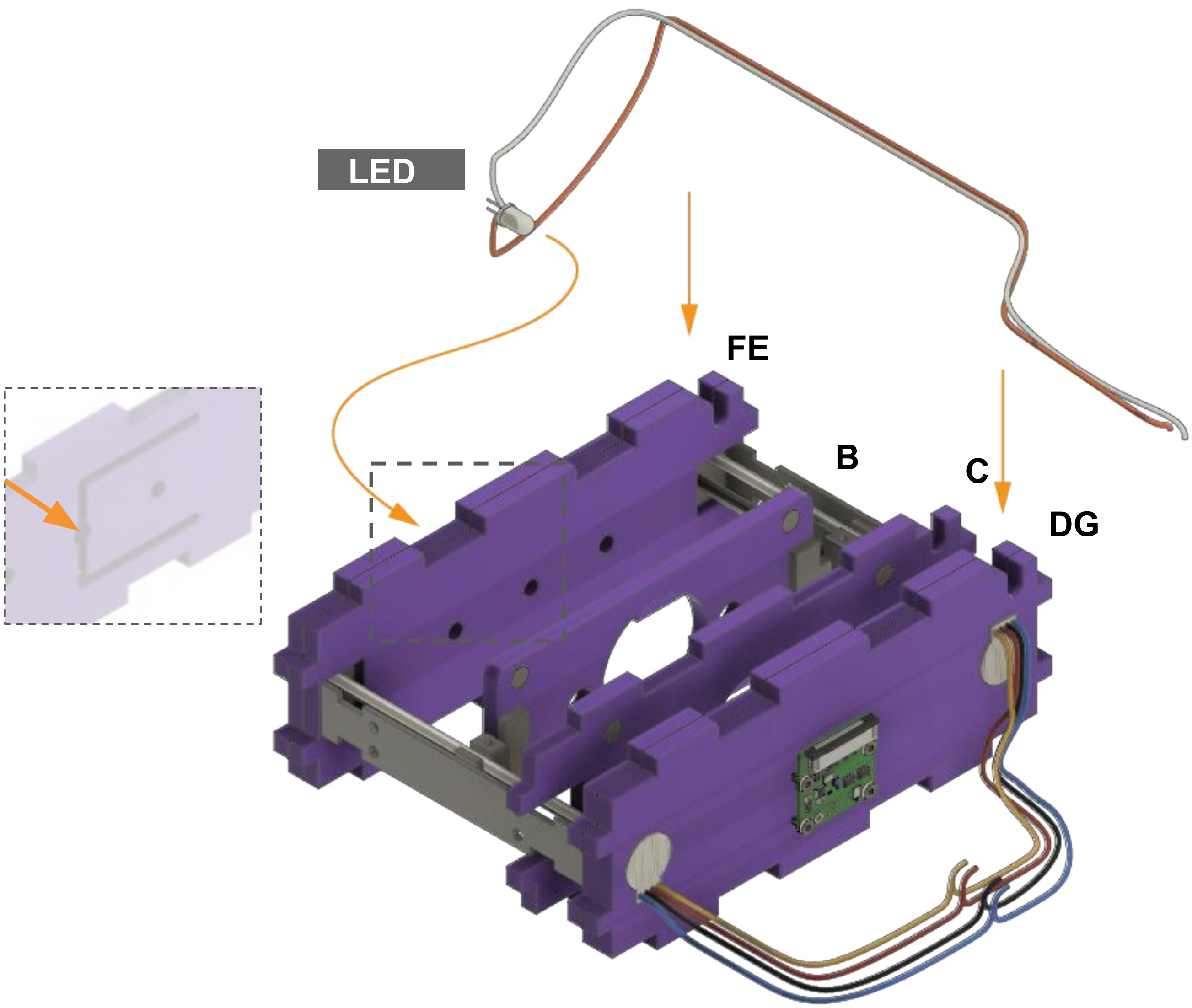


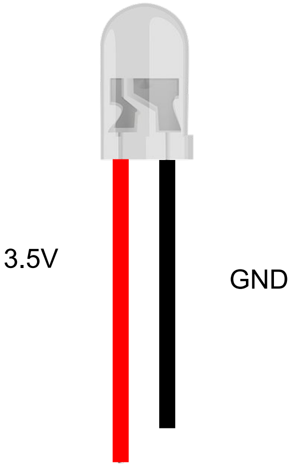


Insert the ends of the stepper motors into the designated holes on the illumination mount **FE**

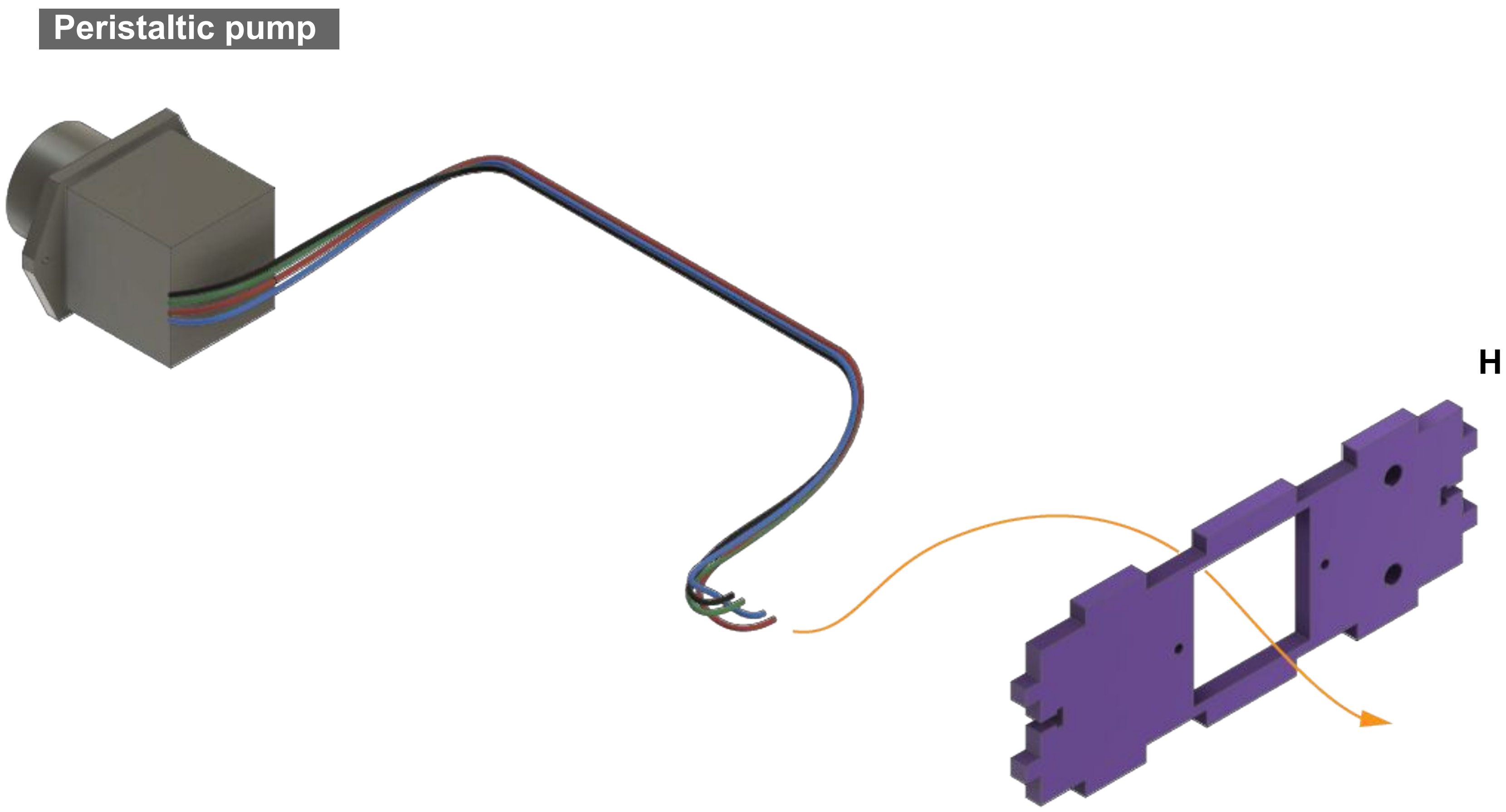


Mount the peristaltic pump on to the pump mount **H,** threading wires through

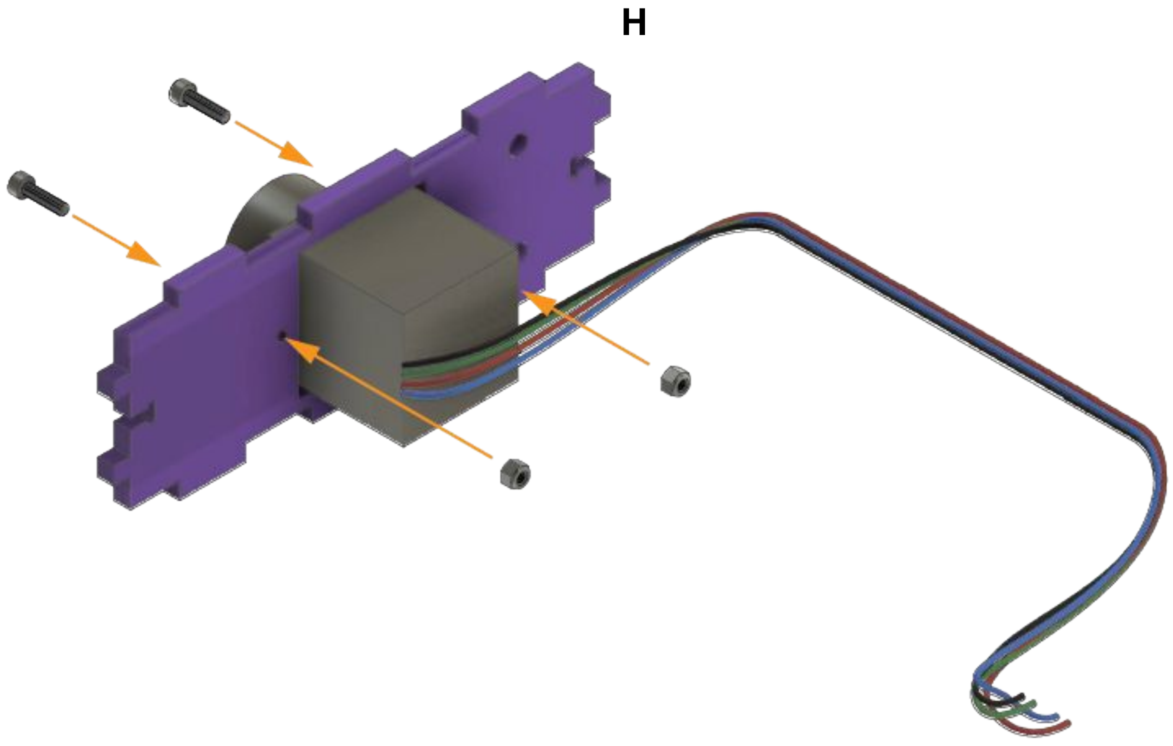


Secure the peristaltic pump to the pump mount **H** with screws and nuts

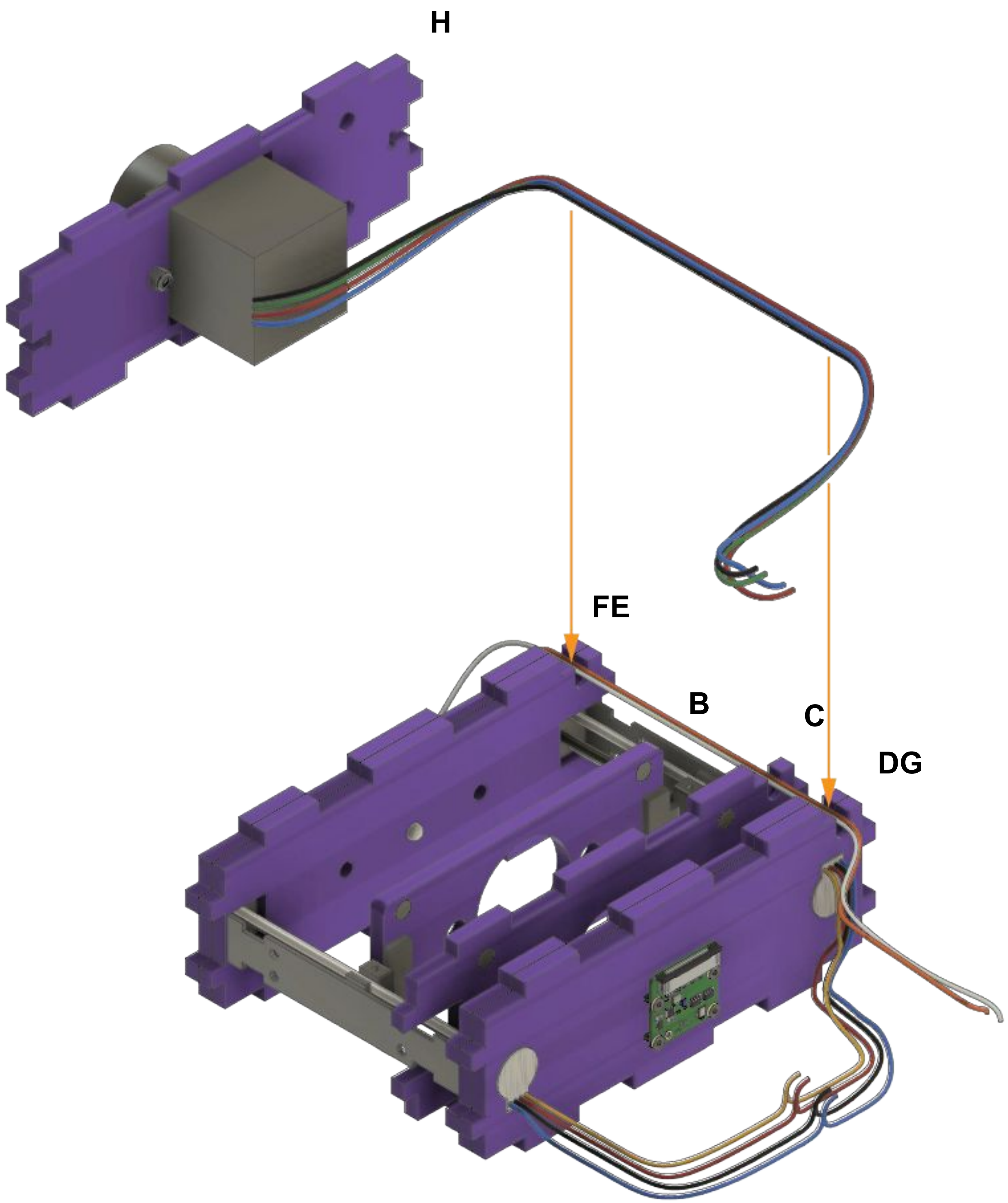


Feed the pump wires through the designated grooves of the assembled motor body

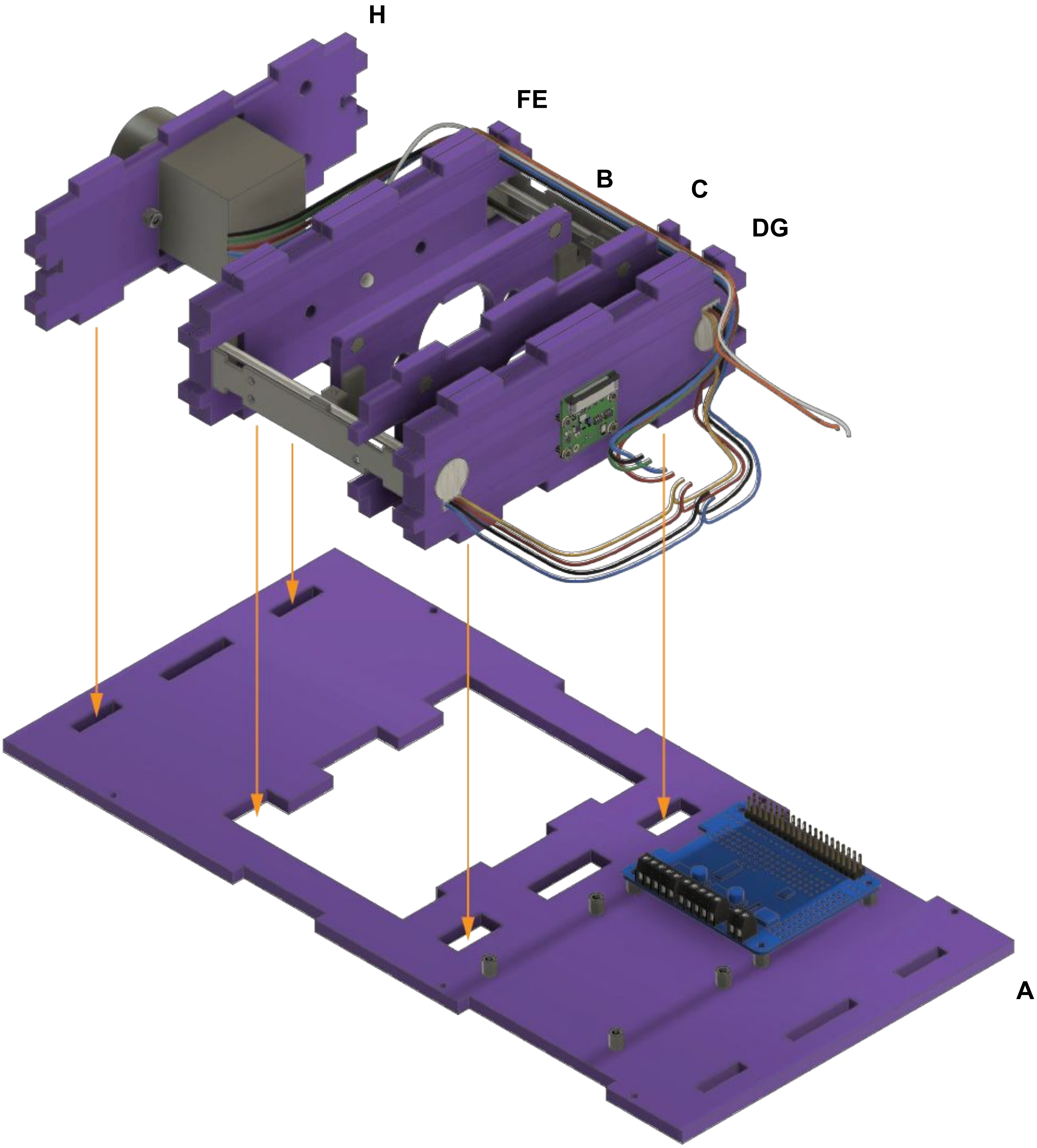


Insert the tabs of the assembled motor body on to the base **A**

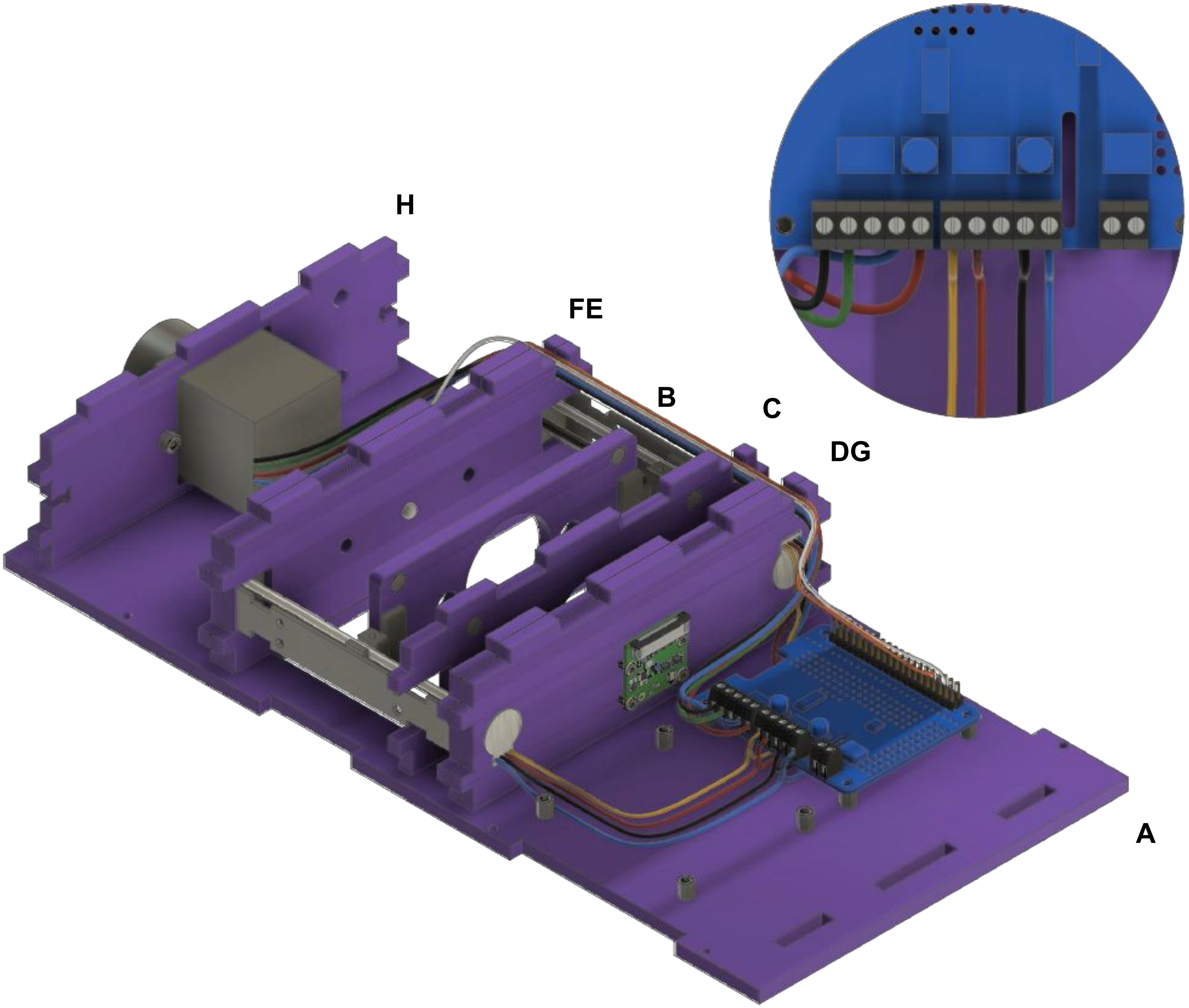


Insert the wires into their designated terminal mounts as shown and secure them by tightening the screws

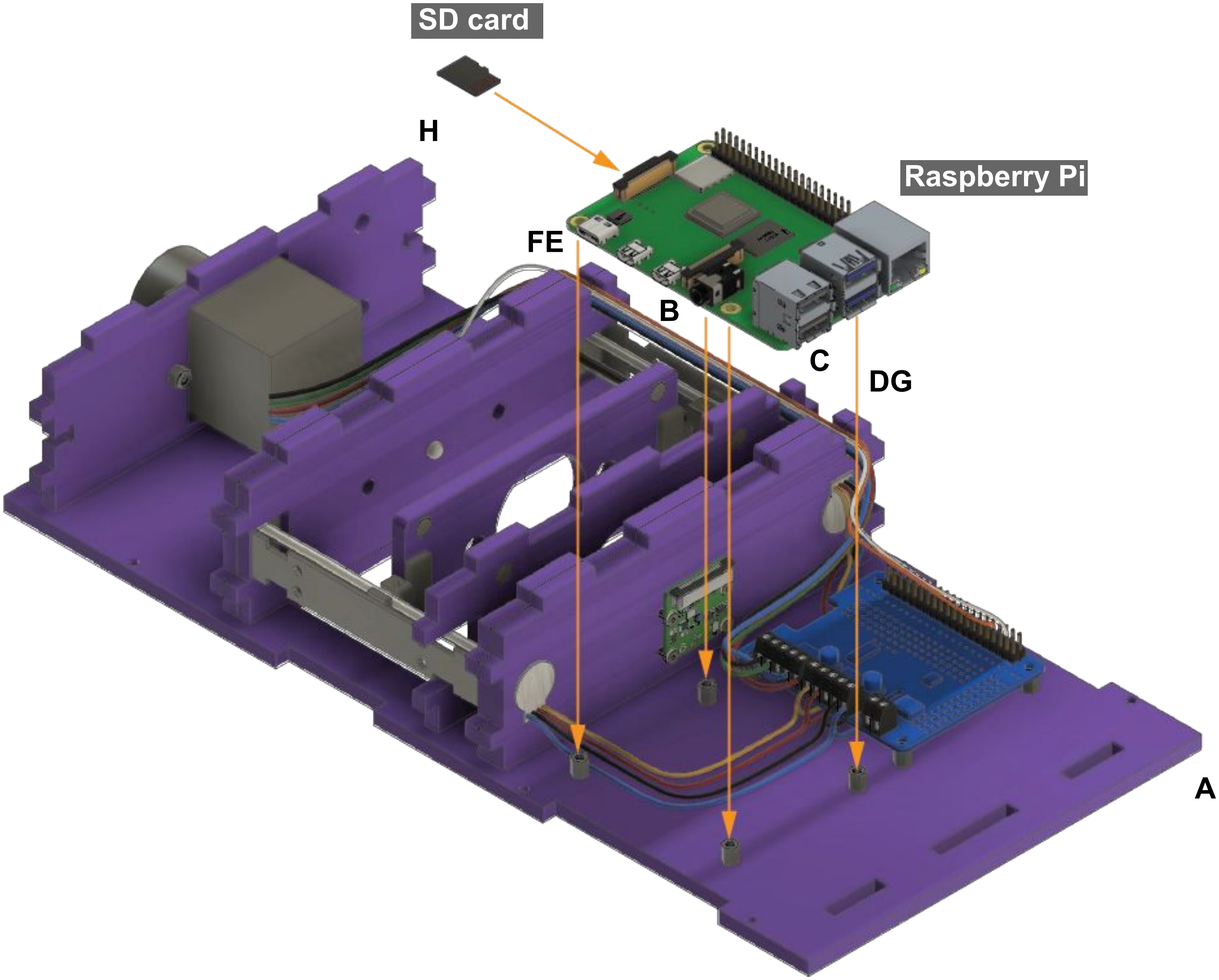


Mount the Raspberry Pi containing the *flashed* SD card on the standoffs attached to the laser cut base **A**

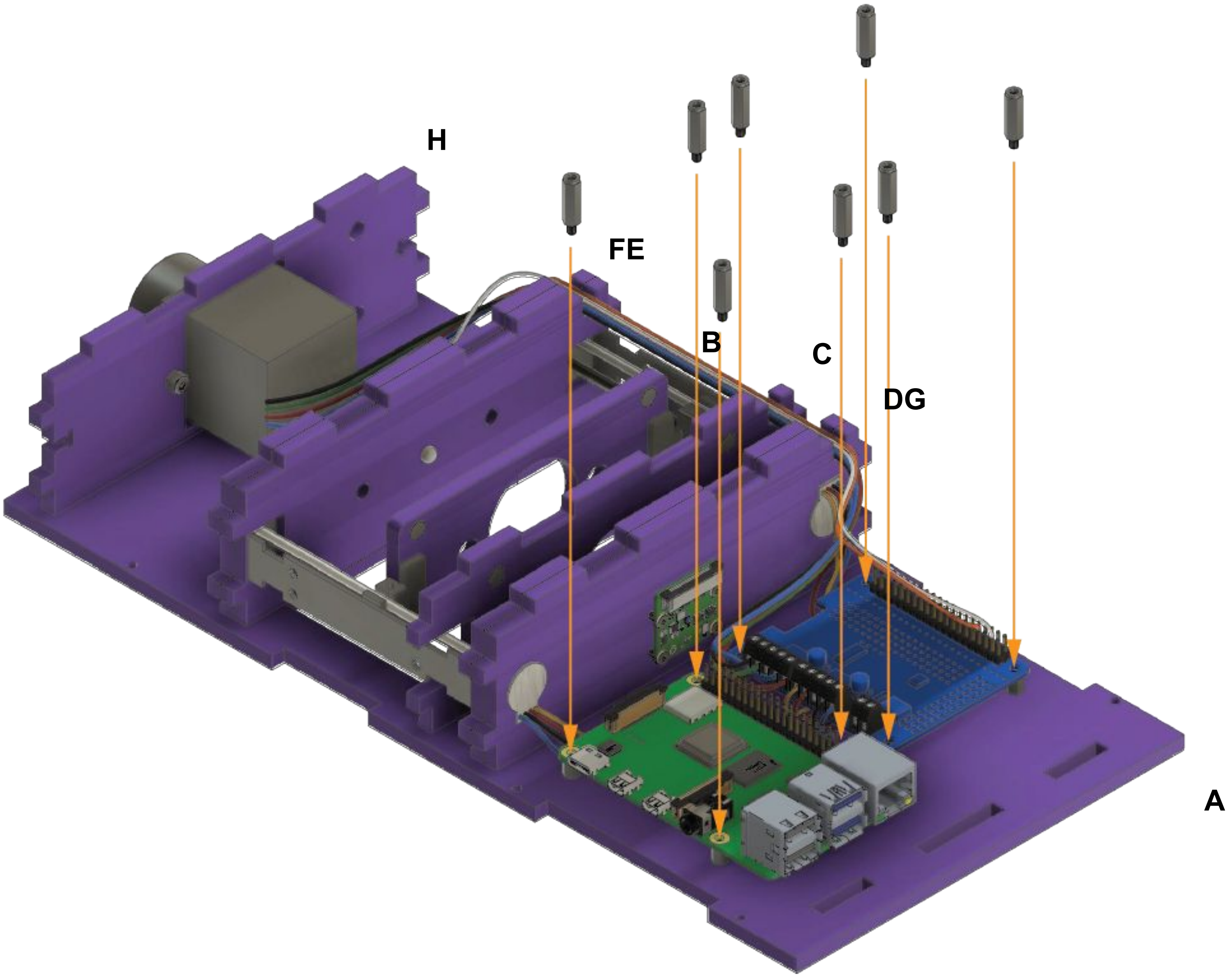


Add 8 standoffs (M2.5 15mm) into the indicated spots on base **A**

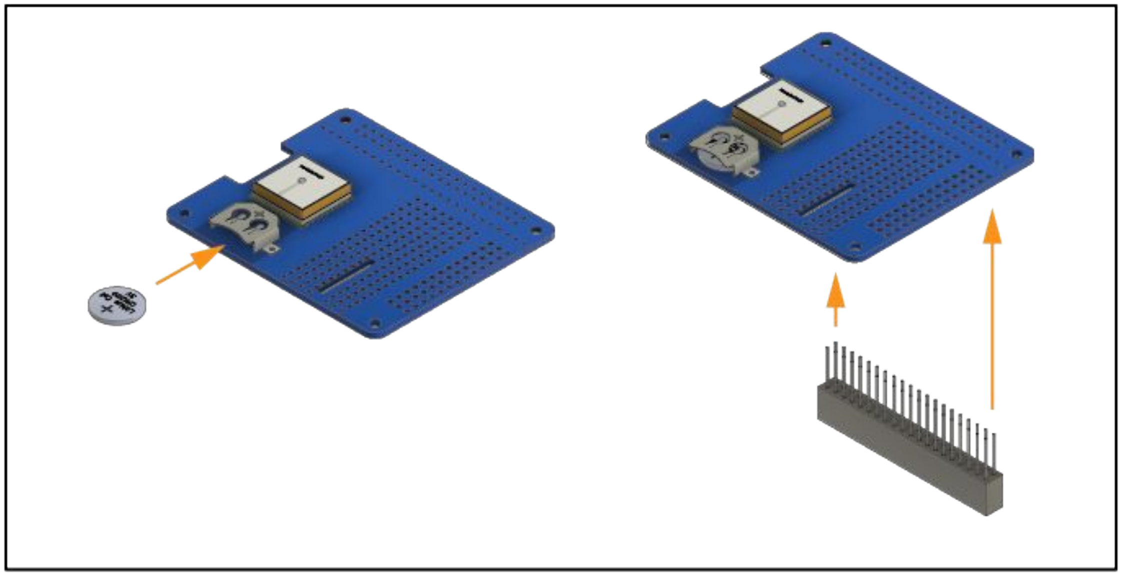

Insert the battery to power the GPS HAT and solder the terminal mounts in place

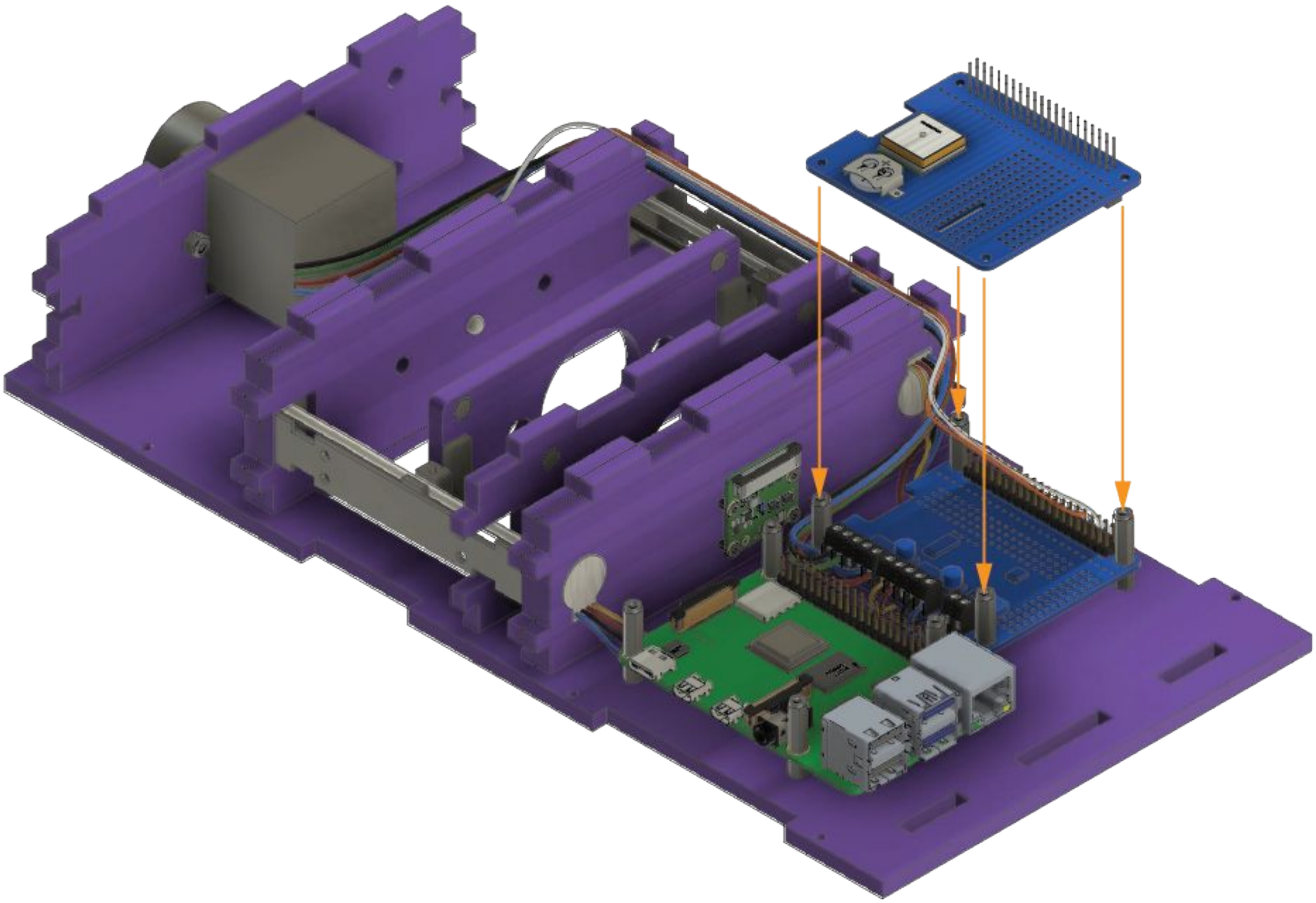

Mount the GPS HAT over the motor driver PCB using the standoffs attached to the laser cut base **A**

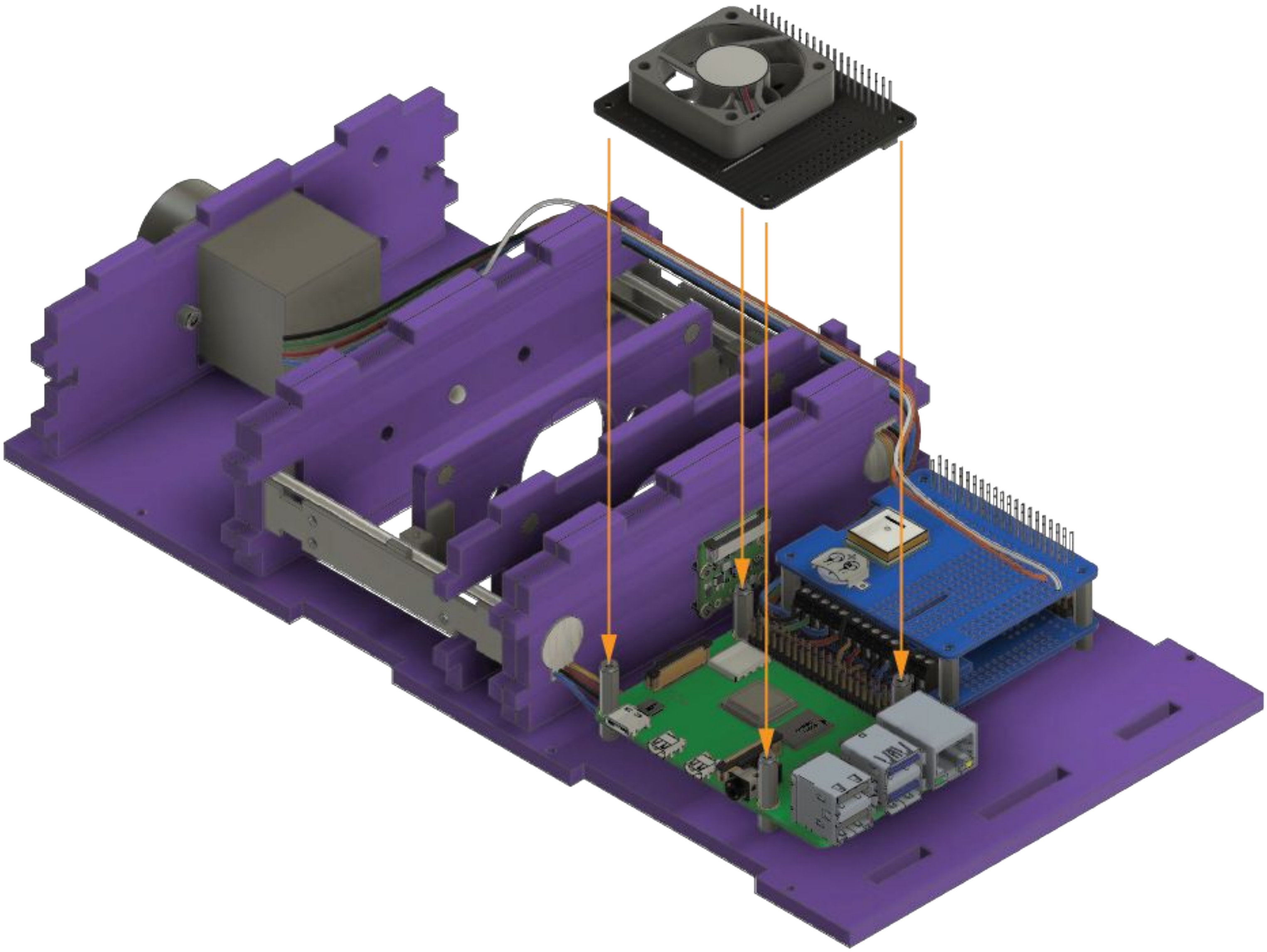

Place the cooling fan HAT above the Raspberry Pi by mounting it to the standoffs on base **A**

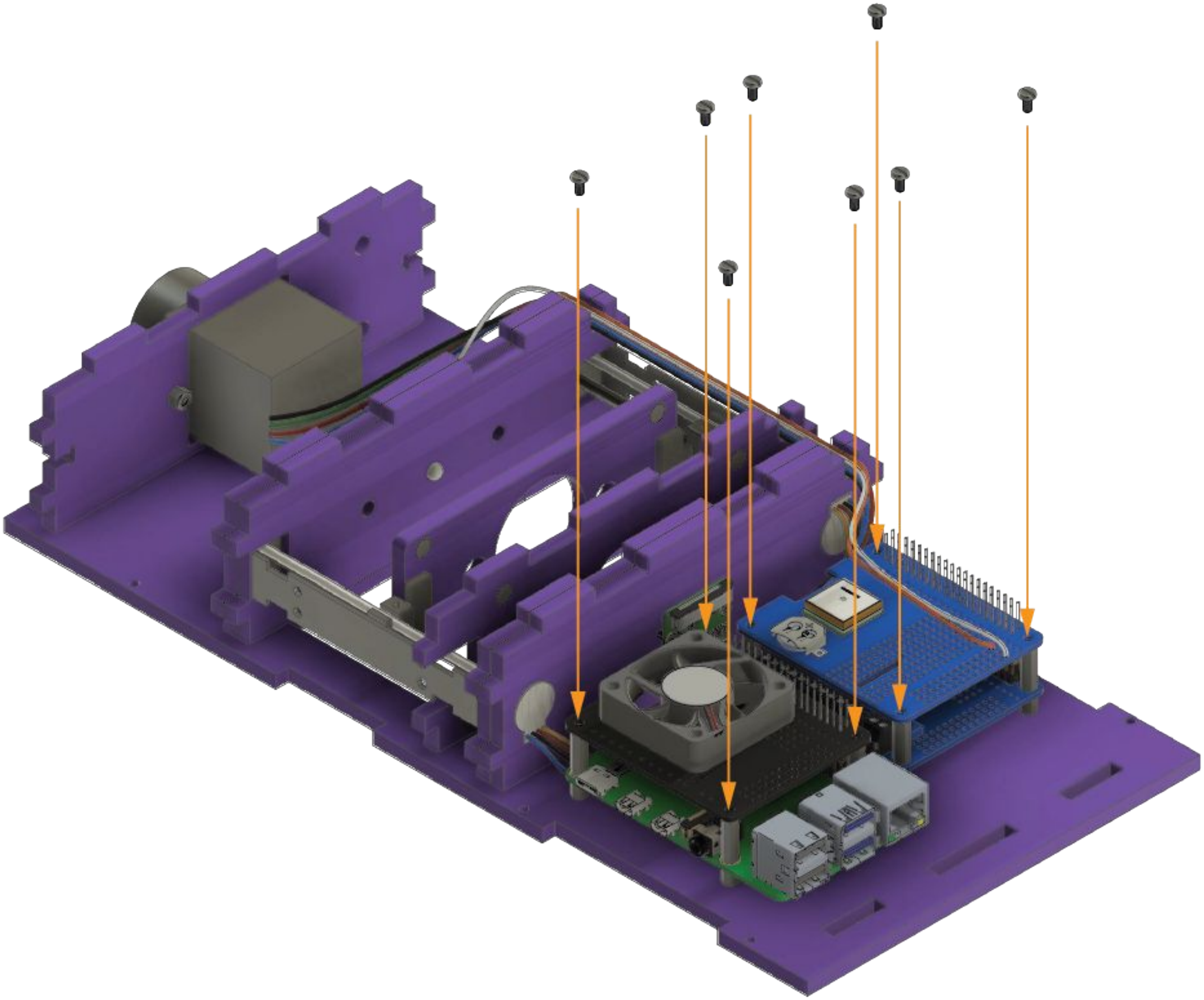

Secure the cooling fan HAT and GPS HAT by tightening the 8 screws to the standoffs on base **A**

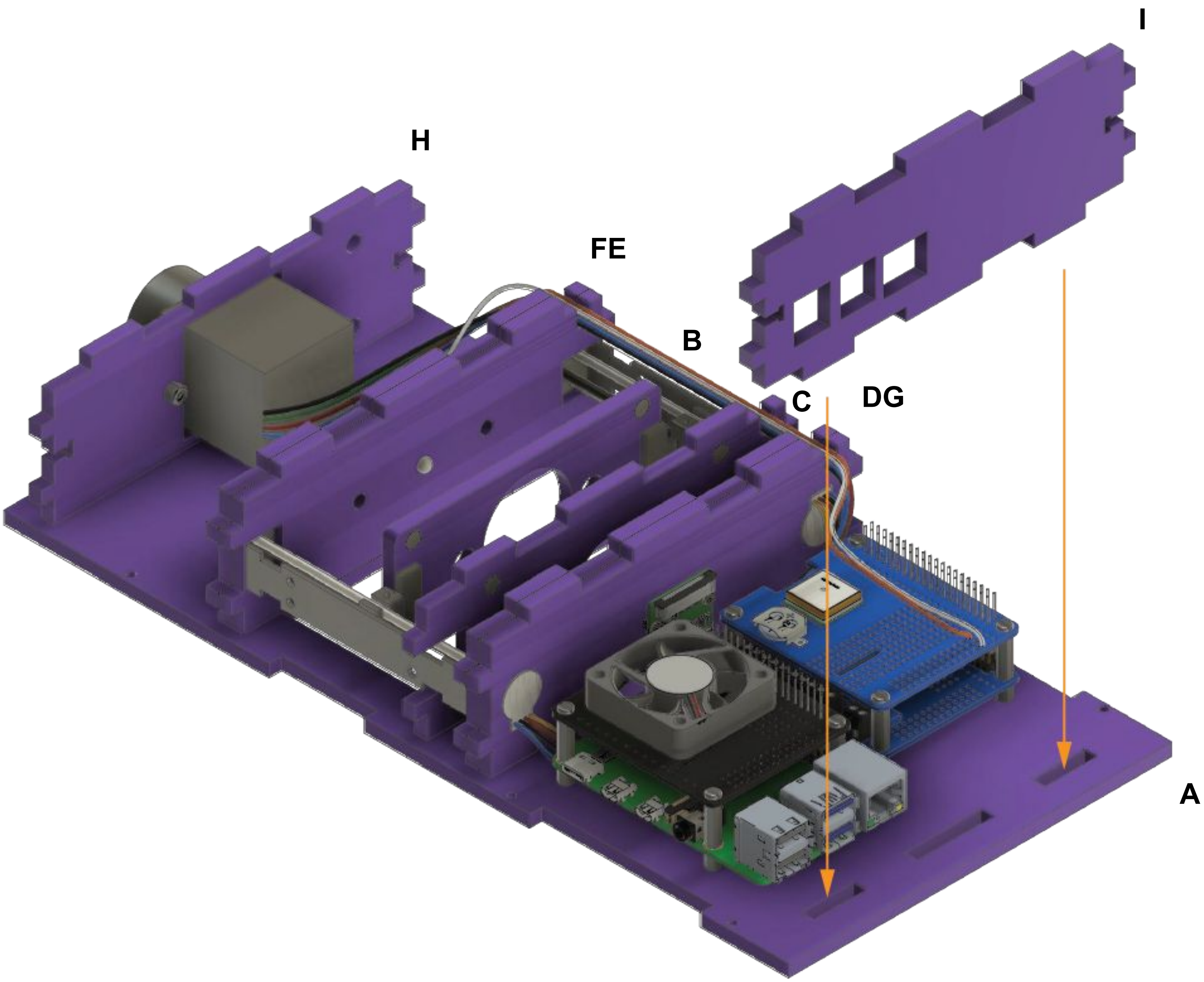


Insert the laser cut border **I** into base **A**

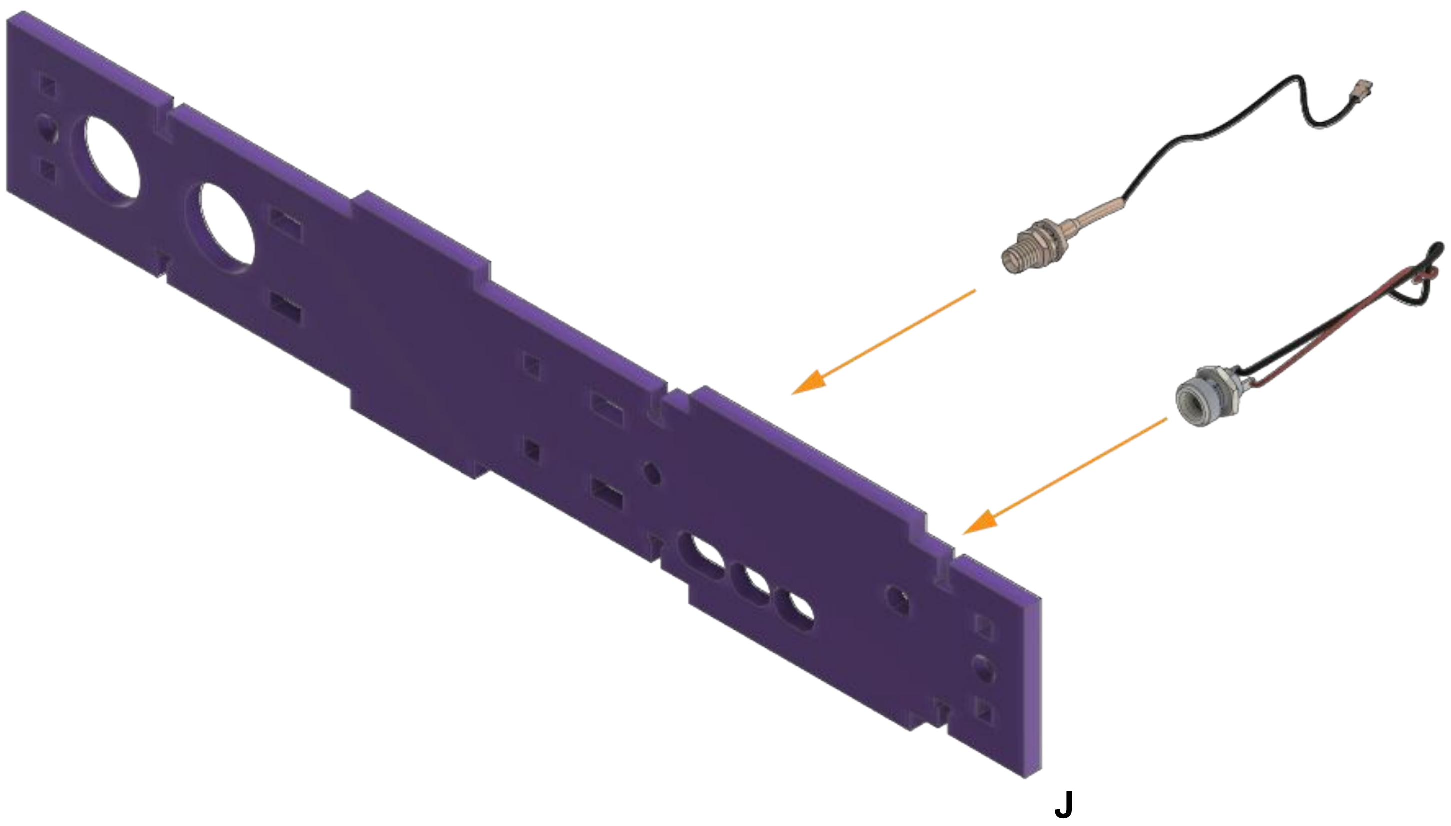


Insert the power and GPS connectors into side plate **J**

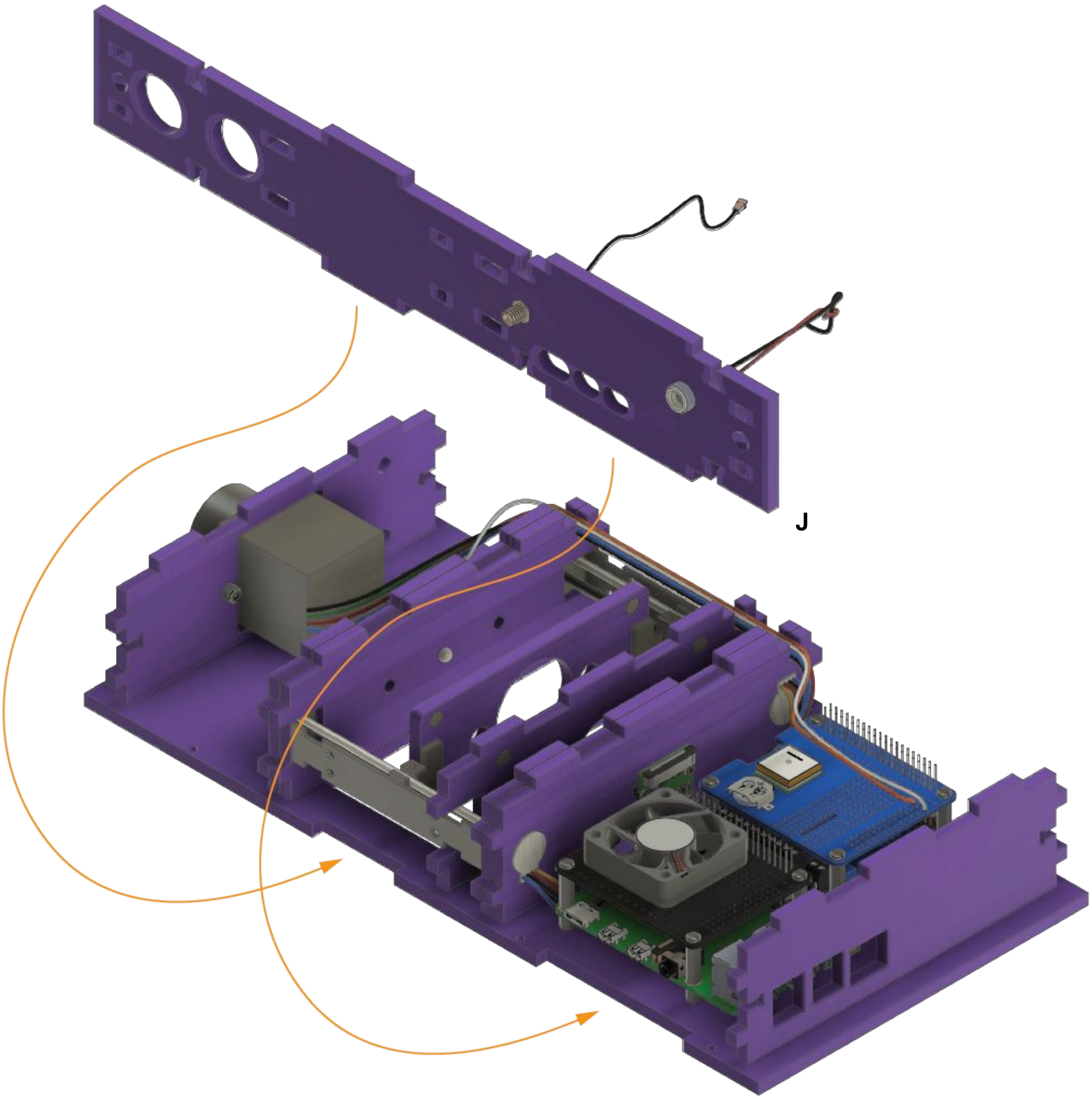


Place the side plate **J** into the designated slots on the base

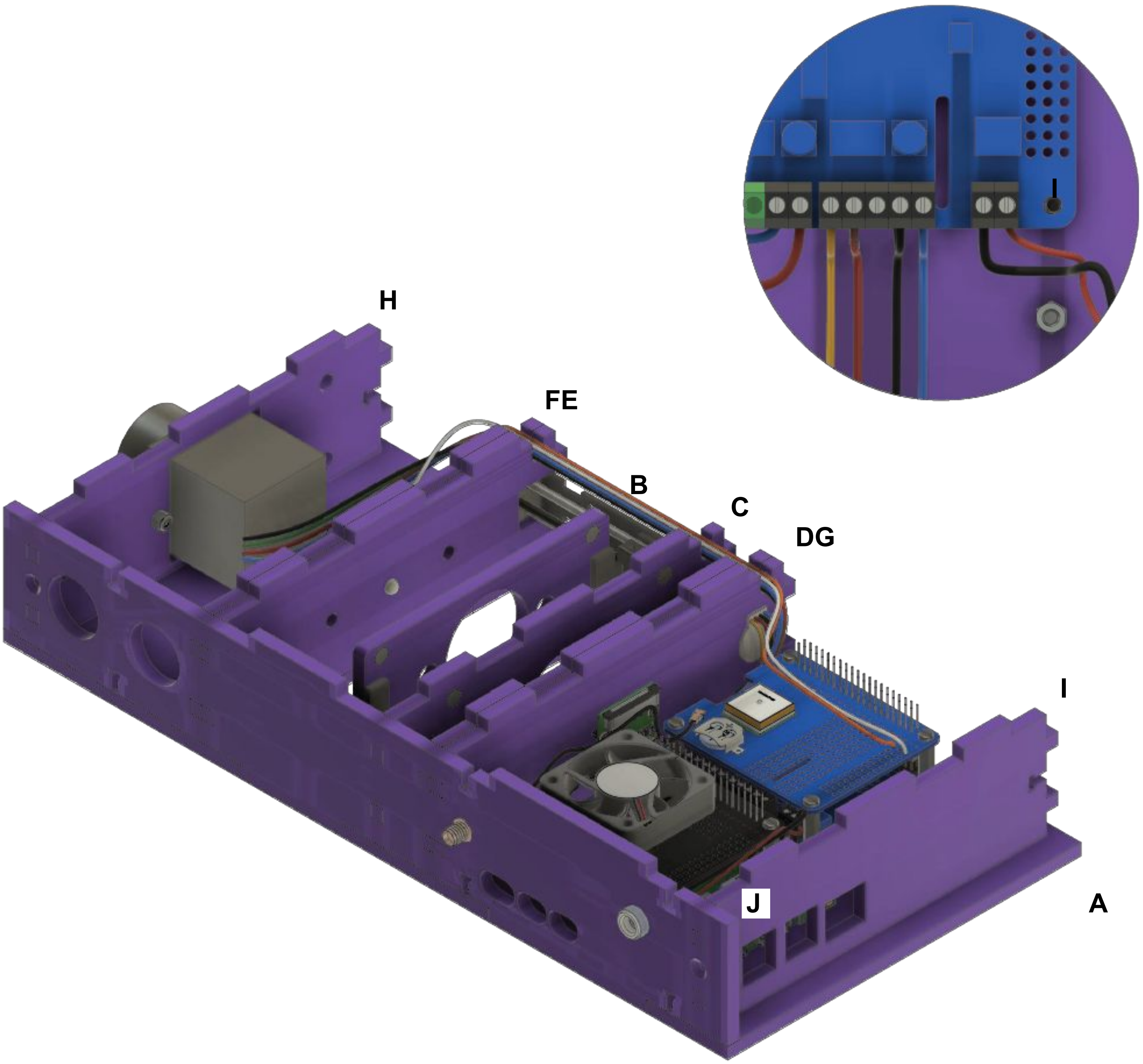


Secure the wires into their designated terminal mounts, check stable connection with a slight tug

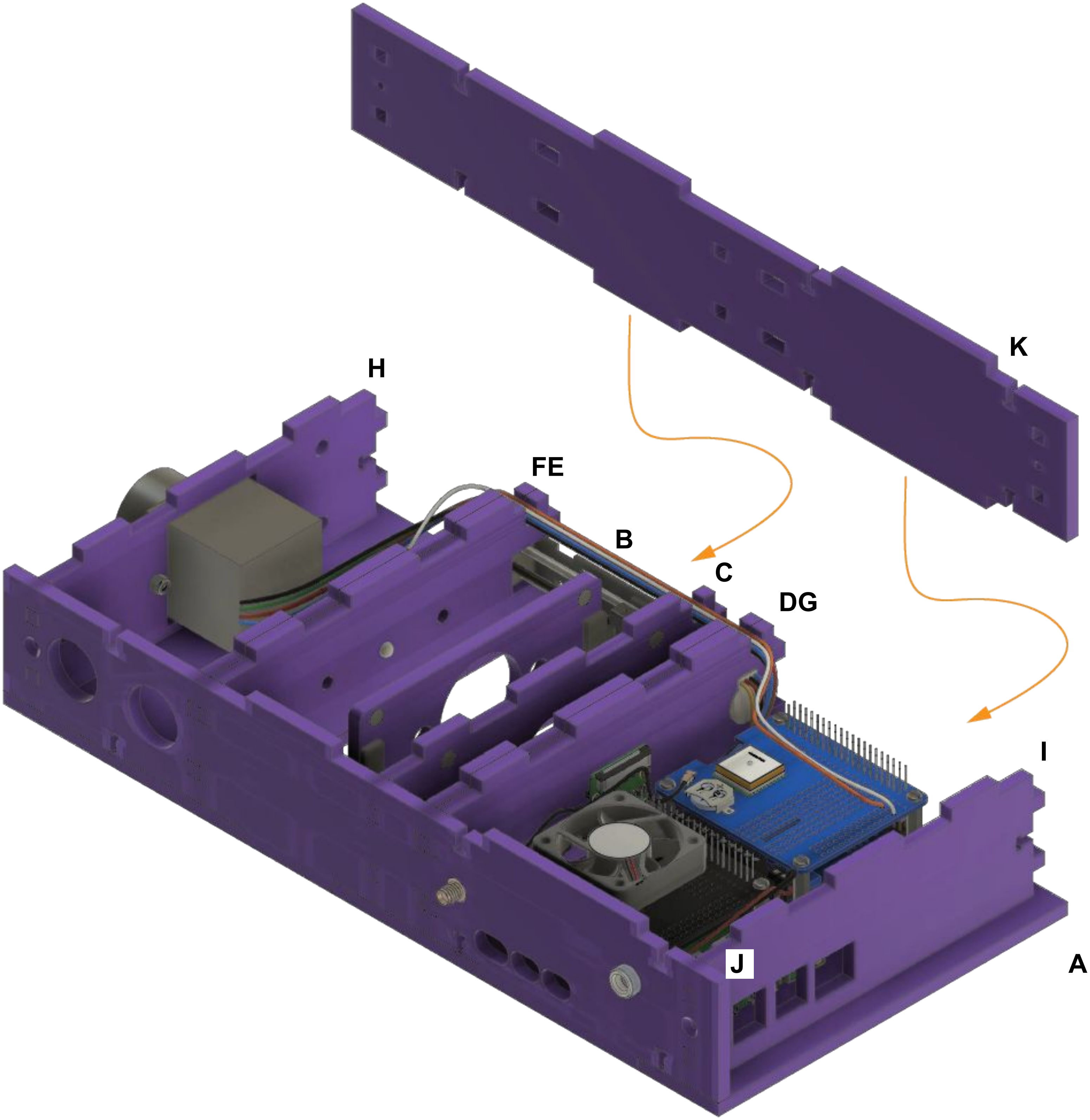


Mount the side plate **K** on base **A** using the assigned slots

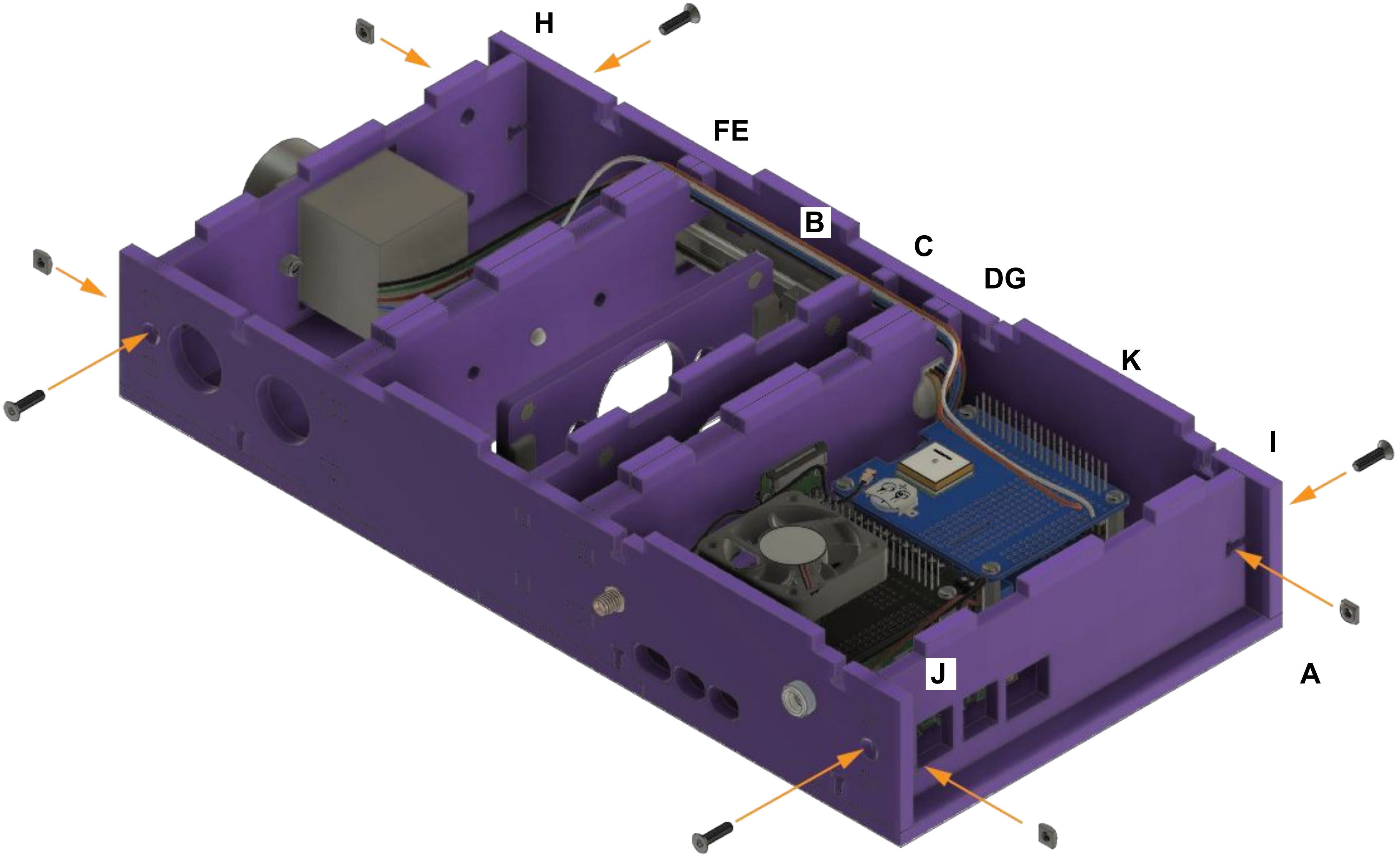


Secure the laser cut sides with the screws and nuts

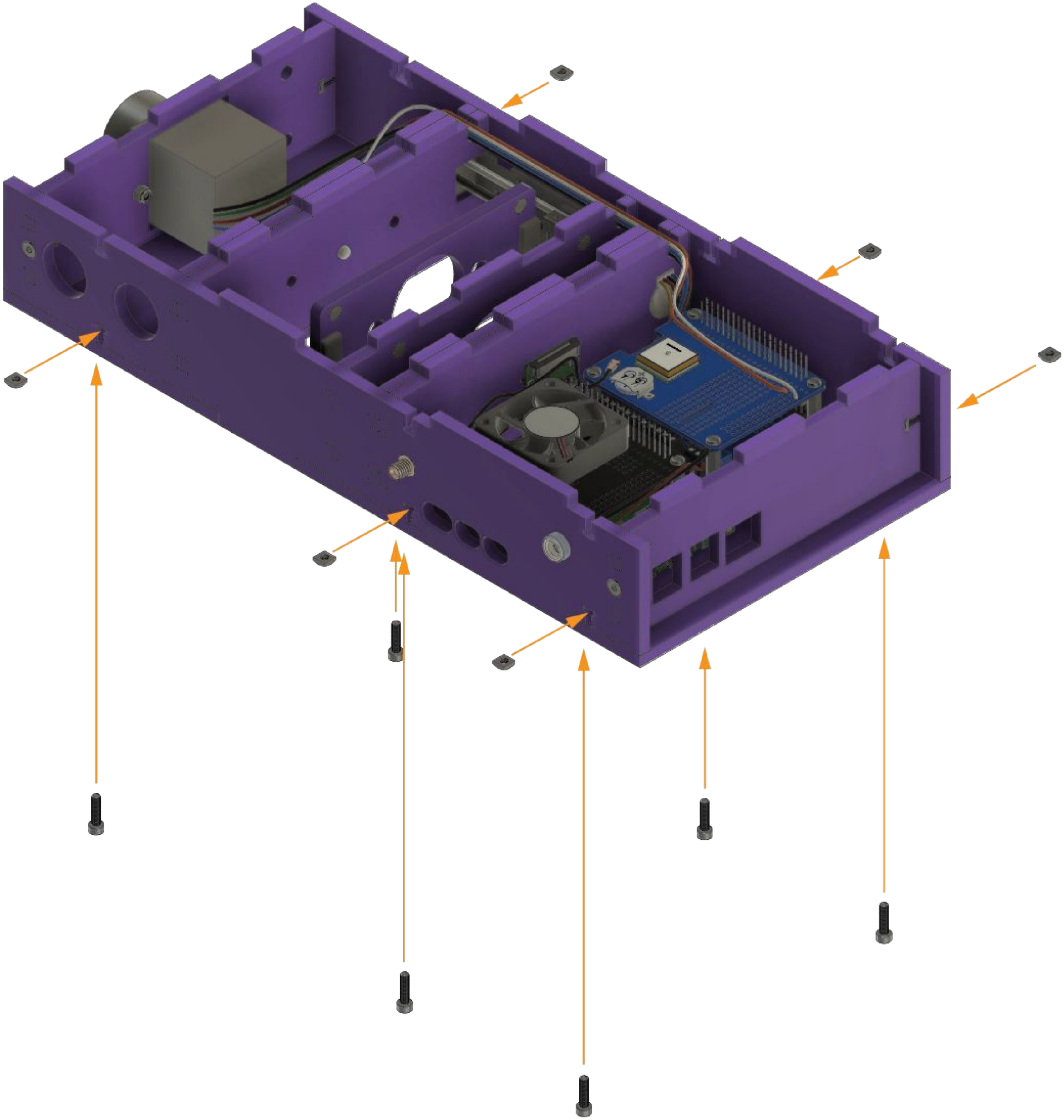

Secure the laser cut sides with the screws and nuts

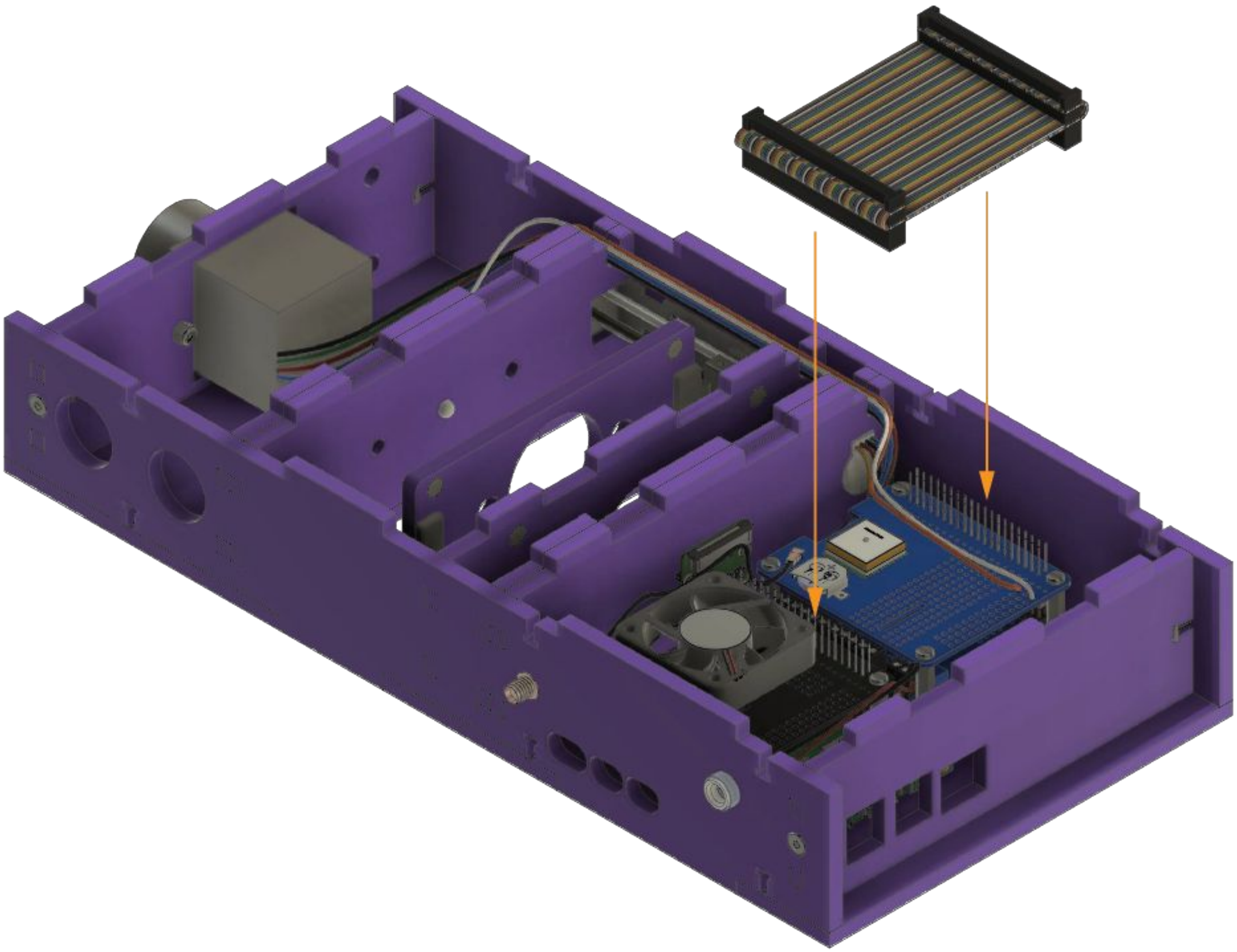

Attach the GPIO ribbon to connect the cooling fan HAT to the GPS HAT

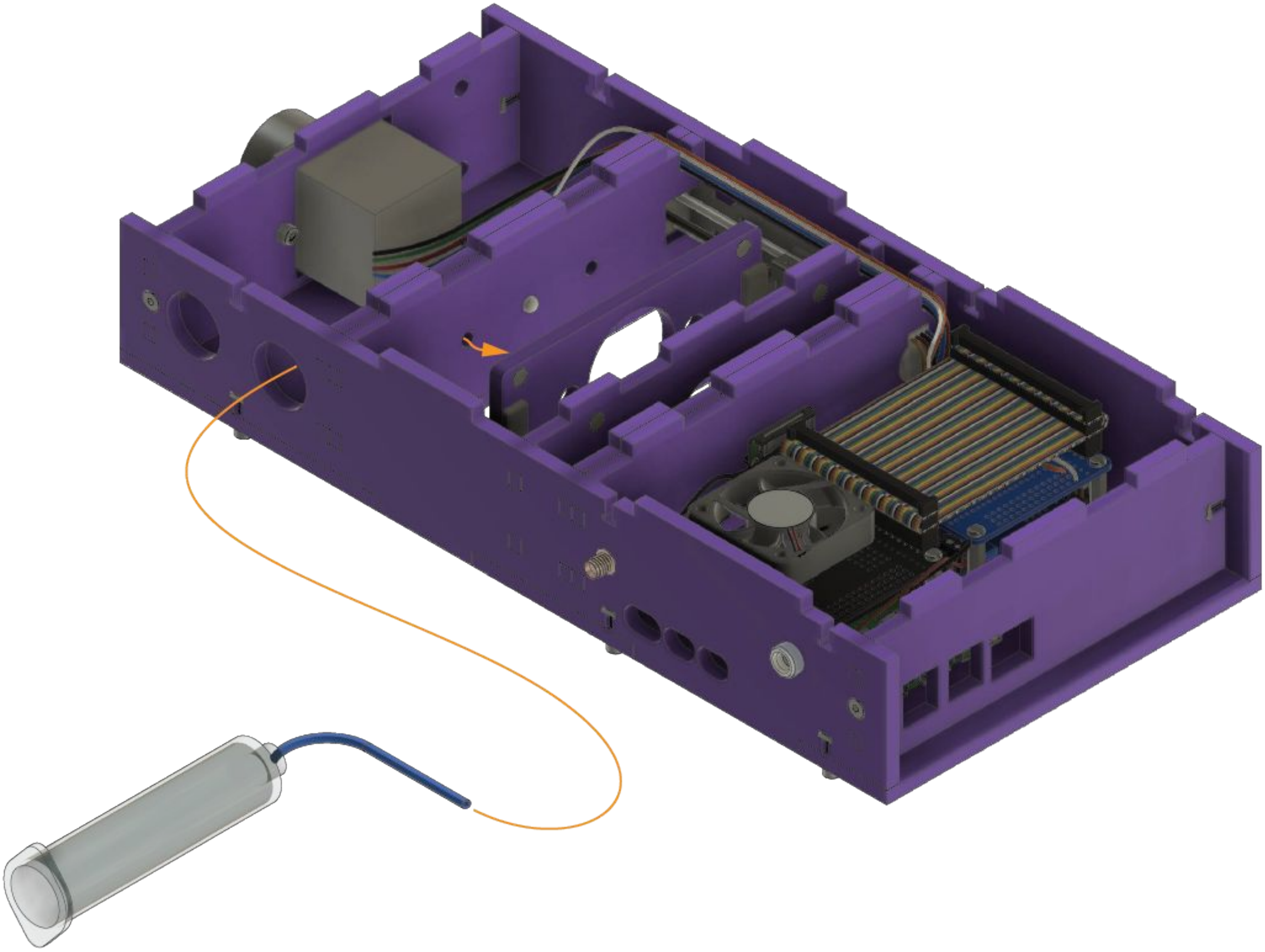

Feed in the tubing from syringe 1 to form the fluidic path as shown

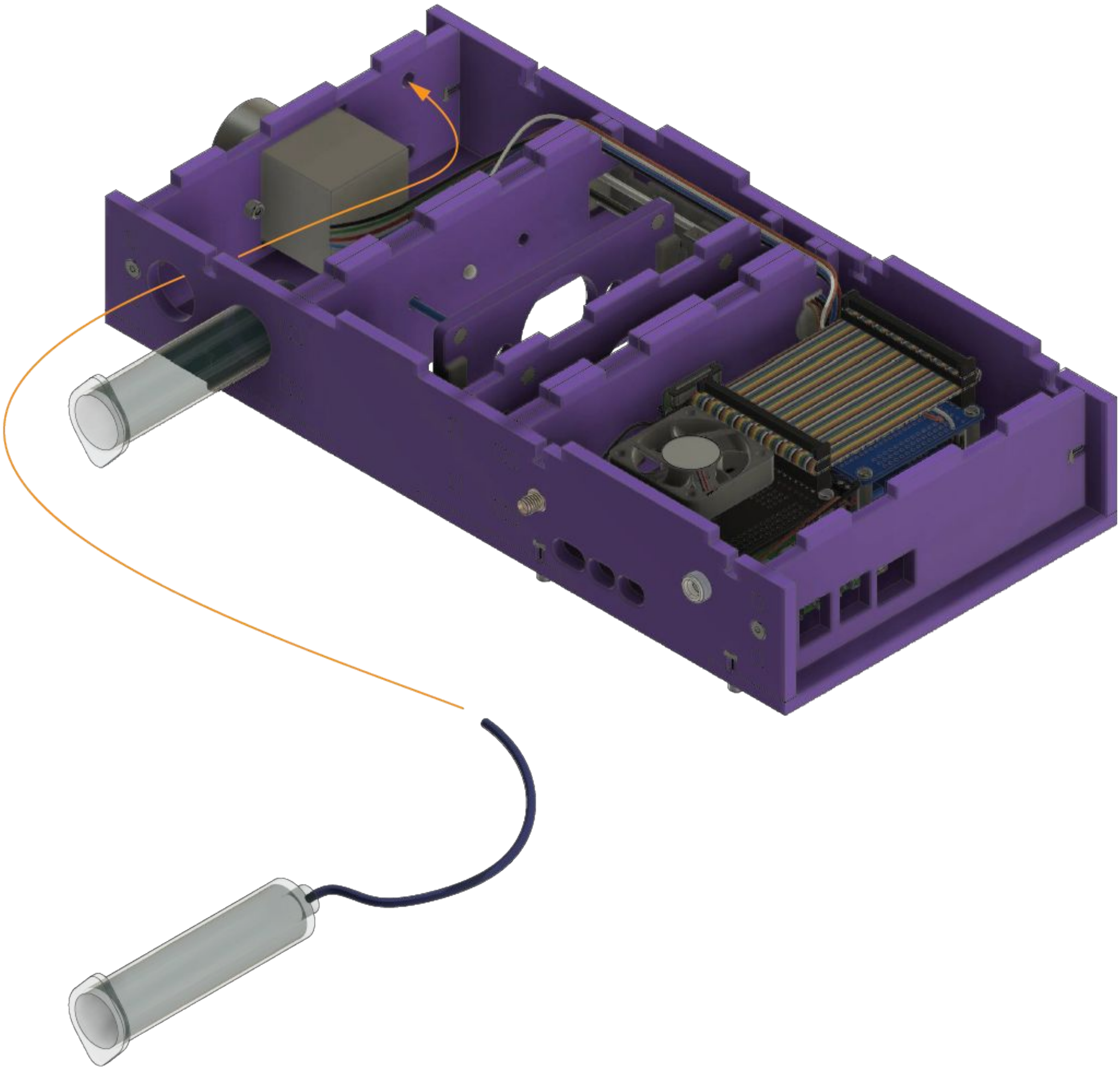

Feed in the tubing from syringe 2 to form the fluidic path as shown

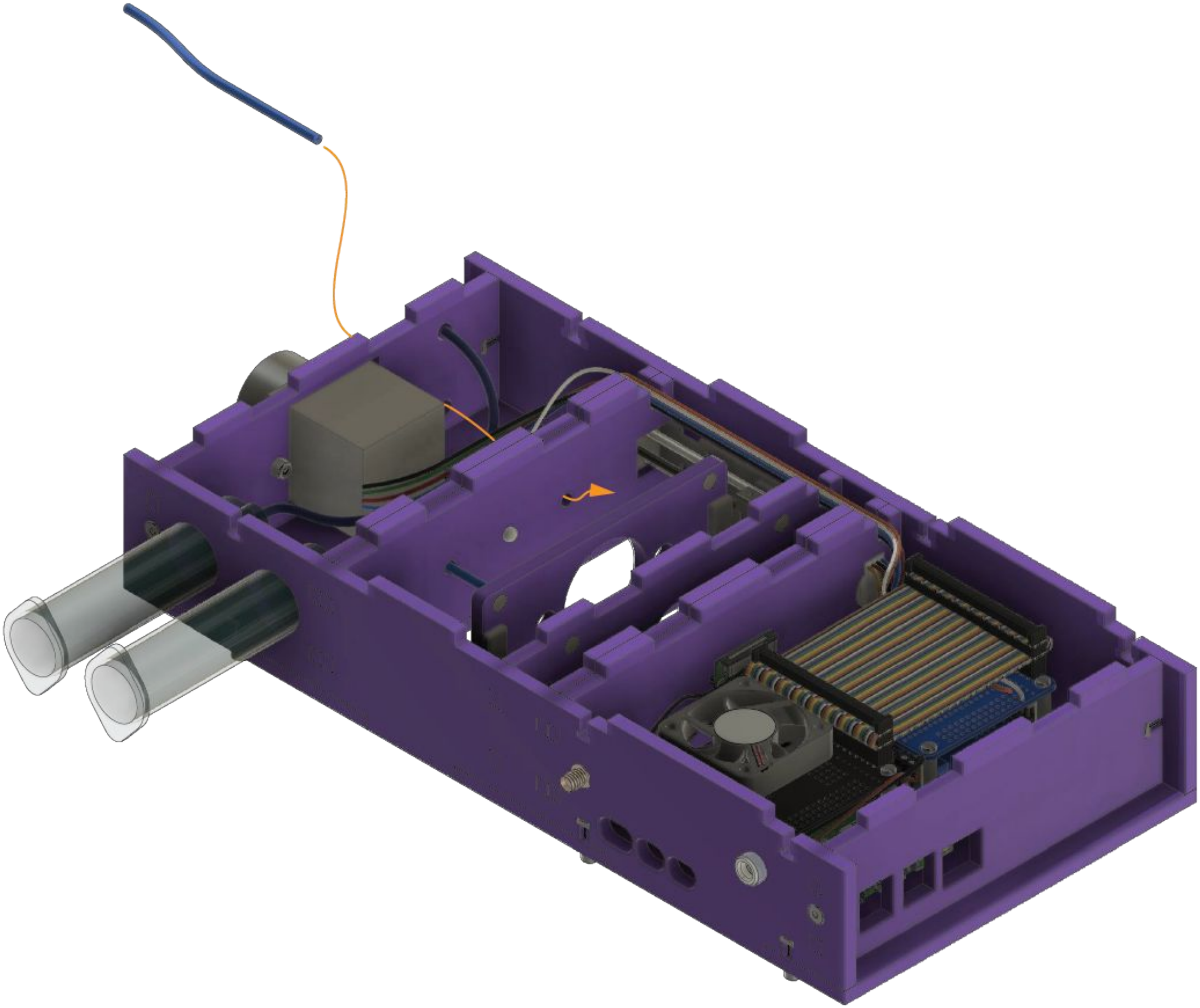

Feed in a length of tubing as shown through motor mount **H** and illumination mount **FE**

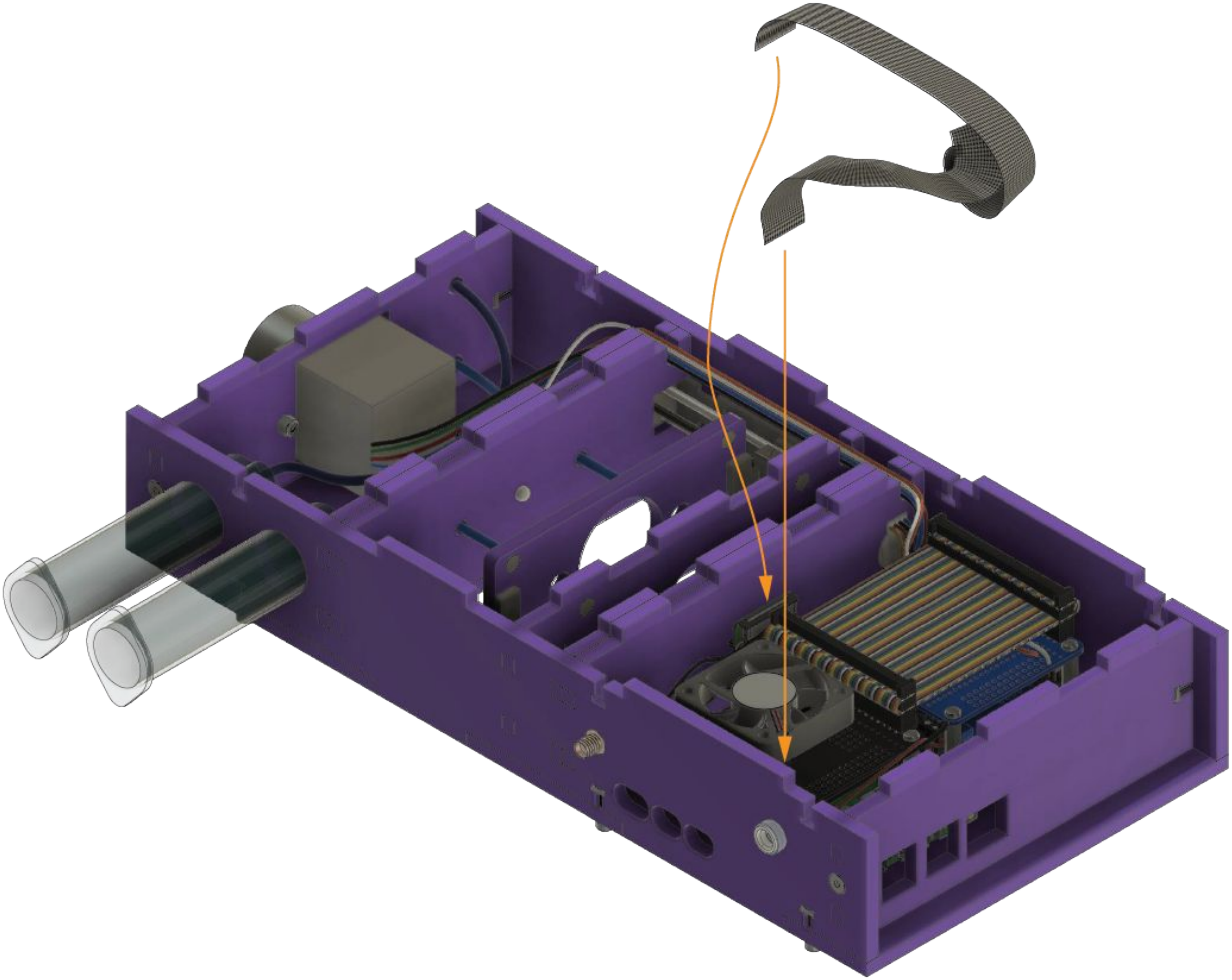

Connect the PiCam with the ribbon cable

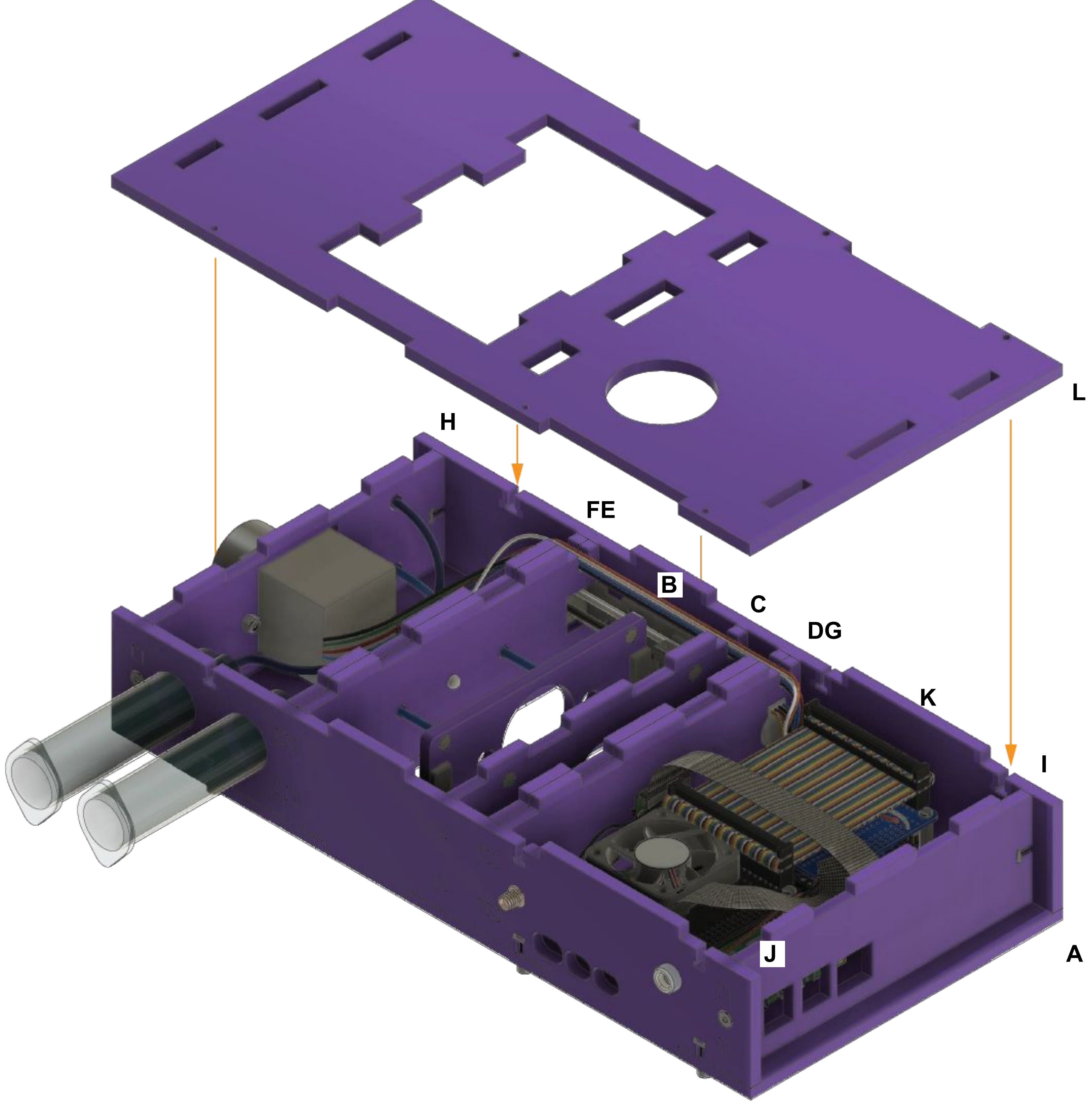


Place the top **L** into the slots on the Planktonscope body

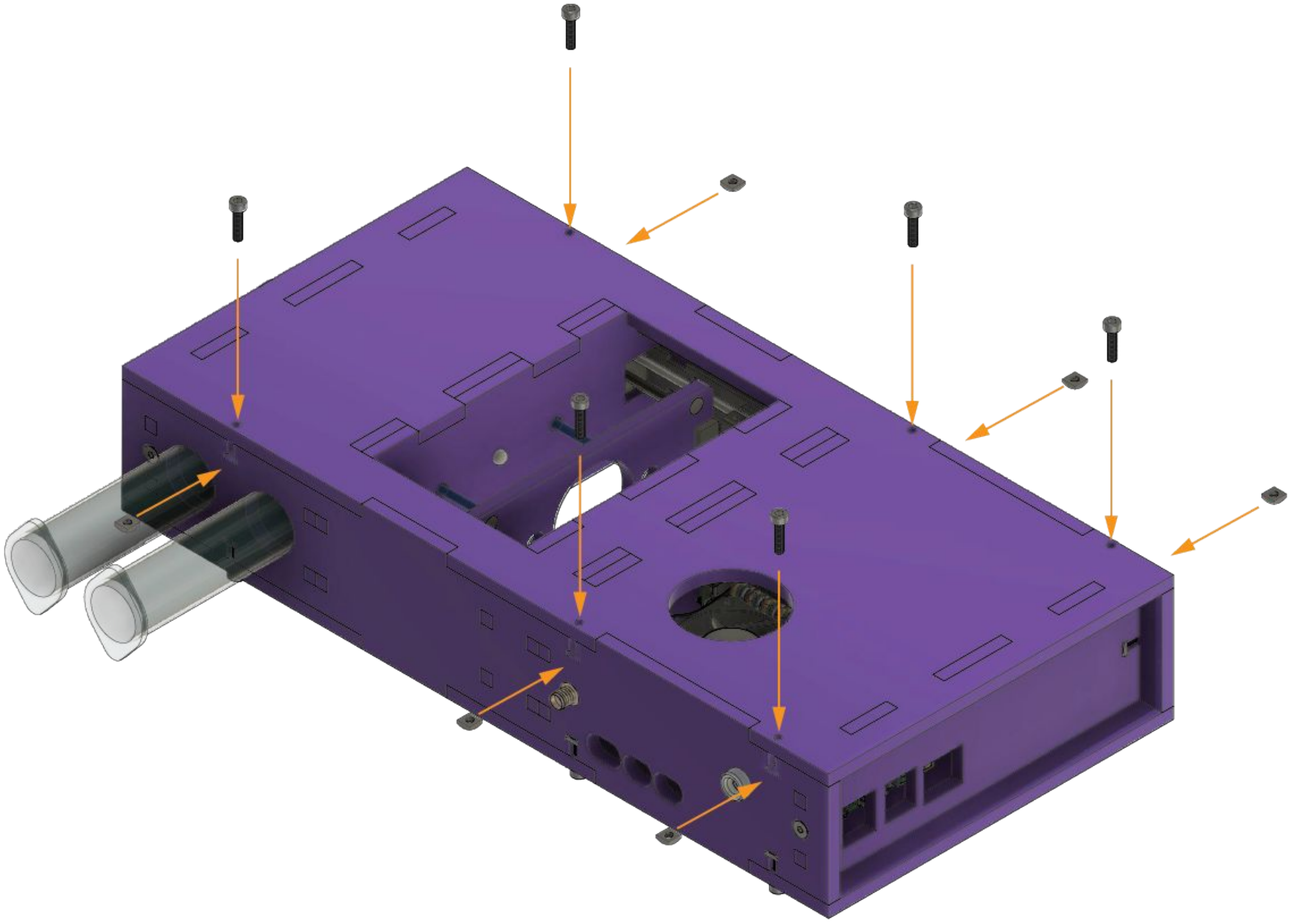

Secure the laser cut top to the side plates with the screws and nuts

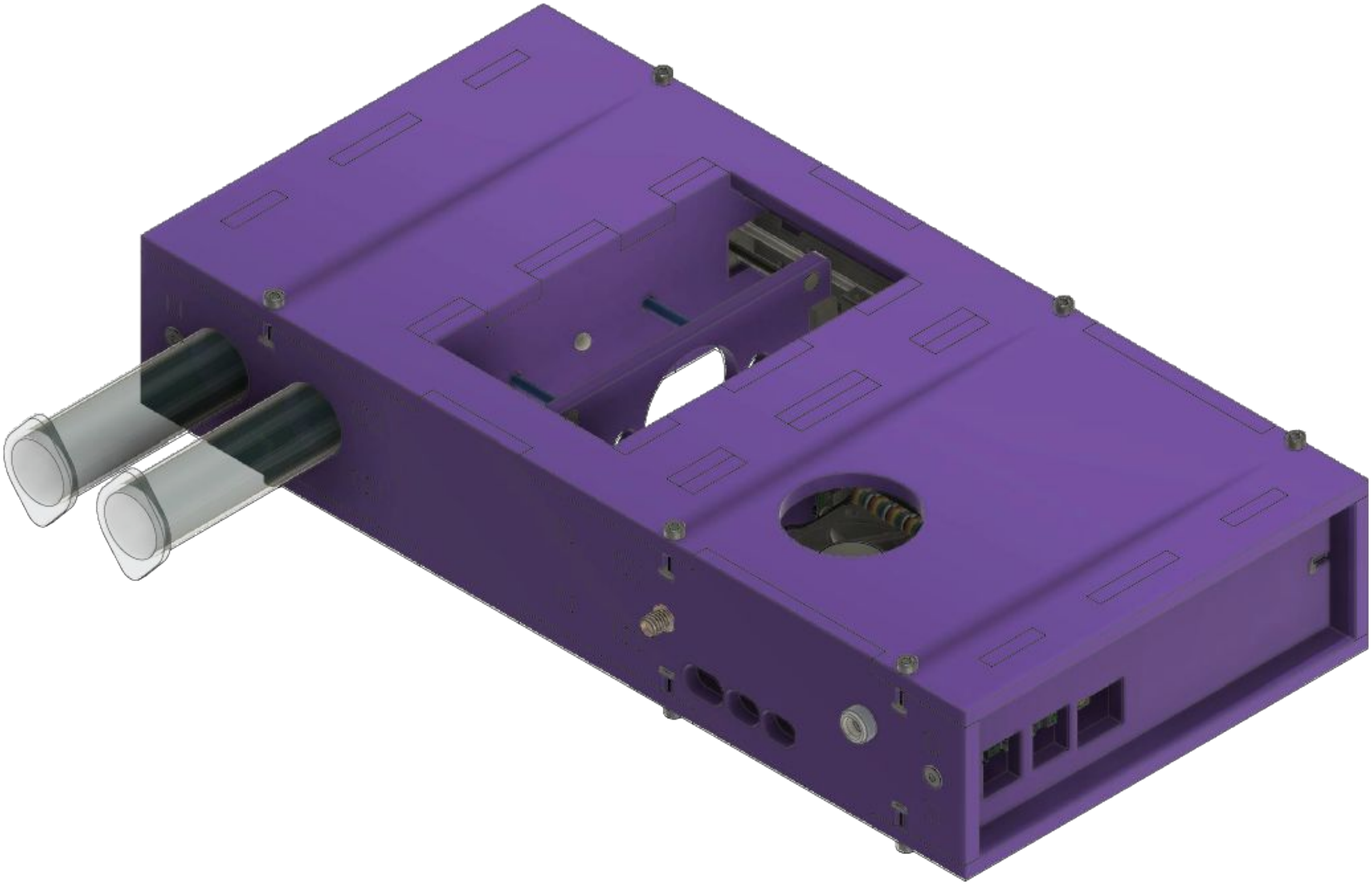

#### You have now assembled the Planktonscope !

The next step is enabling functionality by setting up the software.

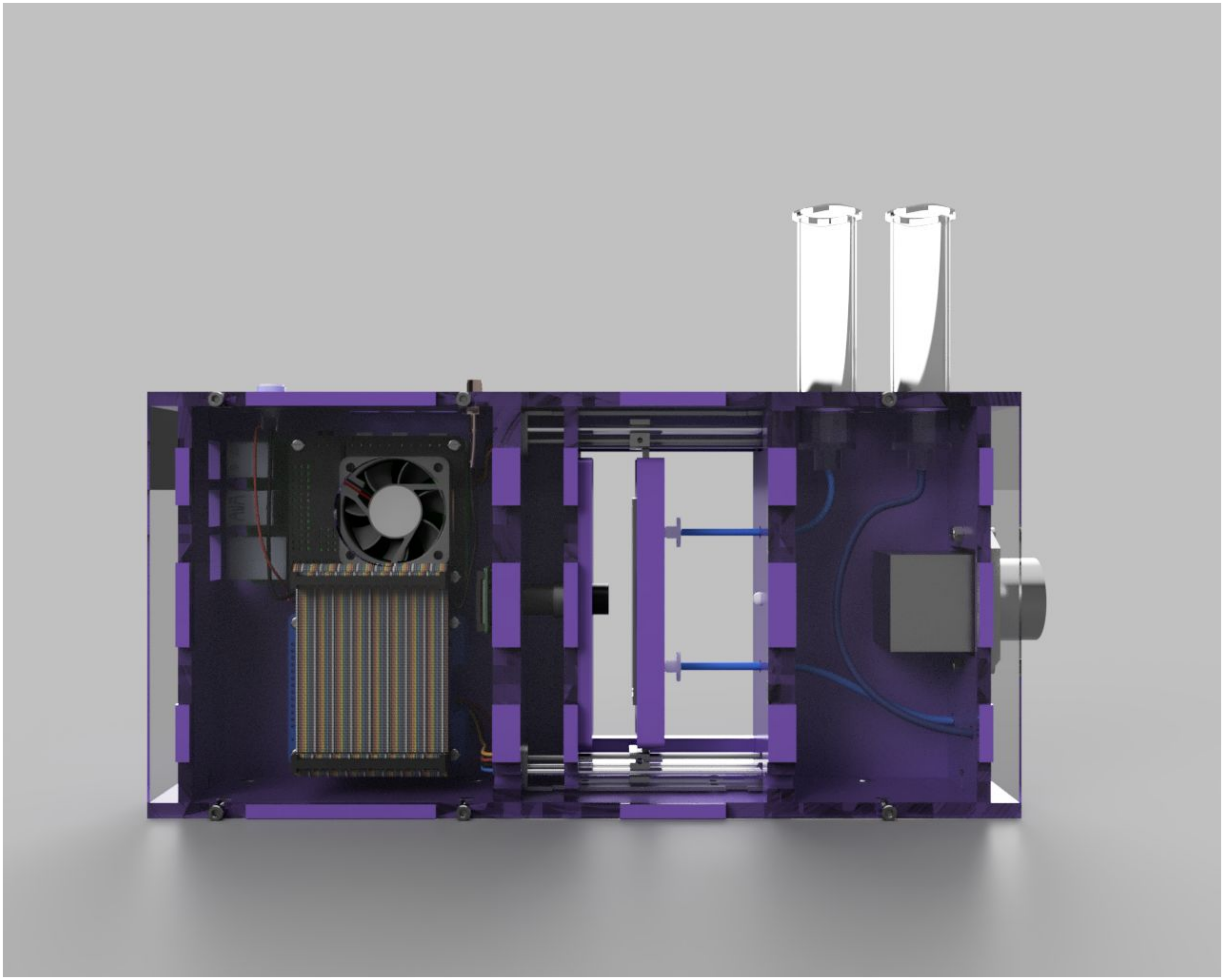
